# Neurocomputational mechanisms underlying the distinct motivational influences of reward and punishment on cognitive control

**DOI:** 10.1101/2025.10.17.682886

**Authors:** Debbie M. Yee, Mahalia Prater Fahey, Ziwei Cheng, Xiamin Leng, Joonhwa Kim, Maisy Tarlow, Kaitlyn Mundy, Samuel Nevins, Amitai Shenhav

## Abstract

Human motivation is fundamentally shaped by one’s expectations of the reward they could earn for good performance or the punishment they would receive for poor performance. However, the extent to which distinct brain regions are selectively associated with specific incentives and/or their corresponding influence on control strategy remains unclear. Using model-based fMRI and a novel multi-incentive control task, we observed distinct neural patterns by incentive valence, with ventral striatum and caudal subregion of dorsal anterior cingulate cortex showing greater sensitivity to rewards whereas inferior frontal gyrus and rostral subregion of dorsal anterior cingulate cortex showing greater sensitivity to penalties. Reward-sensitive regions were associated with increased efficiency (e.g., faster responding with moderate decreases in accuracy) whereas penalty-sensitive regions were associated with increased caution (e.g., slower responding with increased accuracy). We disentangled the global and selective influences of motivation on control processes and subjective experience, providing novel insight into the neurocomputational mechanisms of how effort is determined by expected reward and punishment.

**Teaser:** Distinct dACC sub-networks guide dissociable control strategies, revealing how dissociable mental effort profiles are determined by reward and punishment.

## Introduction

A hallmark of adaptive cognition is the capacity to mobilize cognitive resources in accordance with shifting motivational demands.^1,2^ For instance, both humans and animals will adjust their effort when motivated at the prospect of earning rewards (e.g., monetary bonus, social praise) or receiving punishments (e.g., pay reduction, social admonishment).^3,4^ Recent work has shown that humans are more likely to prioritize efficiency (responding quickly) when rewarded for good performance, and more likely to prioritize caution (responding slowly and accurately) when penalized for poor performance.^5–8^ These differential control strategies arise from a common cost-benefit analysis, according to which potential future outcomes are weighed against the cost of effort exertion in order to simultaneously configure multiple forms of cognitive control assuming one’s goal of maximizing expected reward rate.^9–11^ However, while the reward rate optimization model predicts distinct influences of reward and penalty on multivariate control configuration,^12^ the neural underpinnings of how diverse incentives drive this dissociation remains poorly understood.

Converging evidence implicates a distributed network of brain regions in linking motivation and cognitive control, comprising dorsal anterior cingulate (dACC), ventral striatum (VS), anterior insula (AI), dorsolateral prefrontal cortex (DLPFC), and inferior frontal gyrus (IFG).^13–15^ The Expected Value of Control (EVC) Theory organizes this neural circuit around a core set of computations that distinguish between temporally distinct sub-processes related to evaluating potential outcomes and their performance contingencies, and identifying and signaling the optimal control strategies in the current situation.^16,17^ The EVC theory has been used to hypothesize about the functions of specific brain regions within this circuit – for instance, proposing that dACC may integrate information about costs and benefits to identify optimal control settings, and that these control strategies may be executed in regions of LPFC and IFG – and to develop experimental and computational tools to manipulate and measure the process by which motivational incentives are translated into adaptive control. To this end, we recently developed the Multi-Incentive Control Task (MICT), which examines how different combinations of incentives (e.g., monetary rewards and penalties tied to performance) translate into different combinations of control strategies.^5,6^ Using a normative process model of the task based on EVC, we showed that people engage distinct control strategies to different degrees based on their optimality under a given incentive – most notably, with higher rewards participants prioritize efficiency (parameterized as higher rates of evidence accumulation and lower decision thresholds); with higher penalties participants prioritize caution (higher decision thresholds).

Here, we probe the neural circuits responsible for enabling the evaluation and integration of incentives, selection of control strategies, and the motivation—and ultimately execution and monitoring of—control. We had participants (N=100) perform the MICT while undergoing fMRI. We observed that distinct brain regions were associated with the evaluation of positive and negative incentive value, finding that caudal dACC and VS were selectively sensitive to greater reward, whereas rostral dACC and IFG were more sensitive to greater penalty. Cue-related activity in regions also dissociated across control configurations that determined performance in the forthcoming intervals. Model-based fMRI analyses (e.g., including cue-related and interval-related activity as neural regressors in our hierarchical Bayesian estimation of drift diffusion model parameters^18–20)^ revealed that caudal dACC, VS, and DLPFC were more strongly associated with higher drift rate (e.g., greater response efficiency), whereas rostral dACC and IFG were more strongly associated with higher threshold (e.g., greater response caution). Incentives were also found to exert a global motivational influence on these brain regions, with increased activity associated with larger reward-related decreases in threshold and larger penalty-related increases in threshold.

The present study establishes a novel link between computational models of incentive-driven control and their neural substrates, as well as the brain signals that represent how incentive motivation shapes one’s effort profile and affect. To more fully capture the motivational processes elicited by these incentives, we examined whether subjective variability in the affective and motivational ratings was associated with control strategies and neural activity. We found that greater differentiation in motivation was positively associated with greater reward-related adjustments in control strategy. Furthermore, we estimated the positive and negative arousal to quantify the ‘anticipatory affect’ associated with each incentive cue.^21,22^ We observed positive arousal was associated with reward-related regions during the cue phase (e.g., ventral striatum, caudal dACC), whereas negative arousal was associated with penalty-related regions during the interval phase (e.g., rostral dACC, IFG), revealing a distinction in affective states associated with dissociable reward and penalty influences on effort profile. Our data show convergence between neural circuits supporting behavioral signatures and subjective evaluation of incentive-modulated adaptive control, ultimately providing greater mechanistic insight into how multiple incentives dynamically modulate the flexible recruitment of cognitive control in real-world effortful choices.

## Results

The overall aim of our study was to examine the neural mechanisms of mental effort allocation and to determine the extent to which differential brain networks underlie how reward vs. penalty incentives drive dissociable control strategies. Participants (N=94) performed our Multi-Incentive Control Task which varied the amount of monetary reward earned for performing well and the amount of monetary loss penalized for performing poorly (Figure 1). We implemented model-based fMRI analyses, including neural regressors from *a priori* ROIs during cue-related and task execution phases and performed hierarchical Bayesian estimation of drift diffusion model parameters. We sought to address two primary questions: first, whether brain regions involved in the evaluation of positive and negative incentives were overlapping or distinct from brain regions processing their influence on effort exertion; and second, whether distinct neural circuits underlie the degree to which specific positive vs. negative incentives promote distinct control strategies for allocating mental effort.

**Fig 1.**
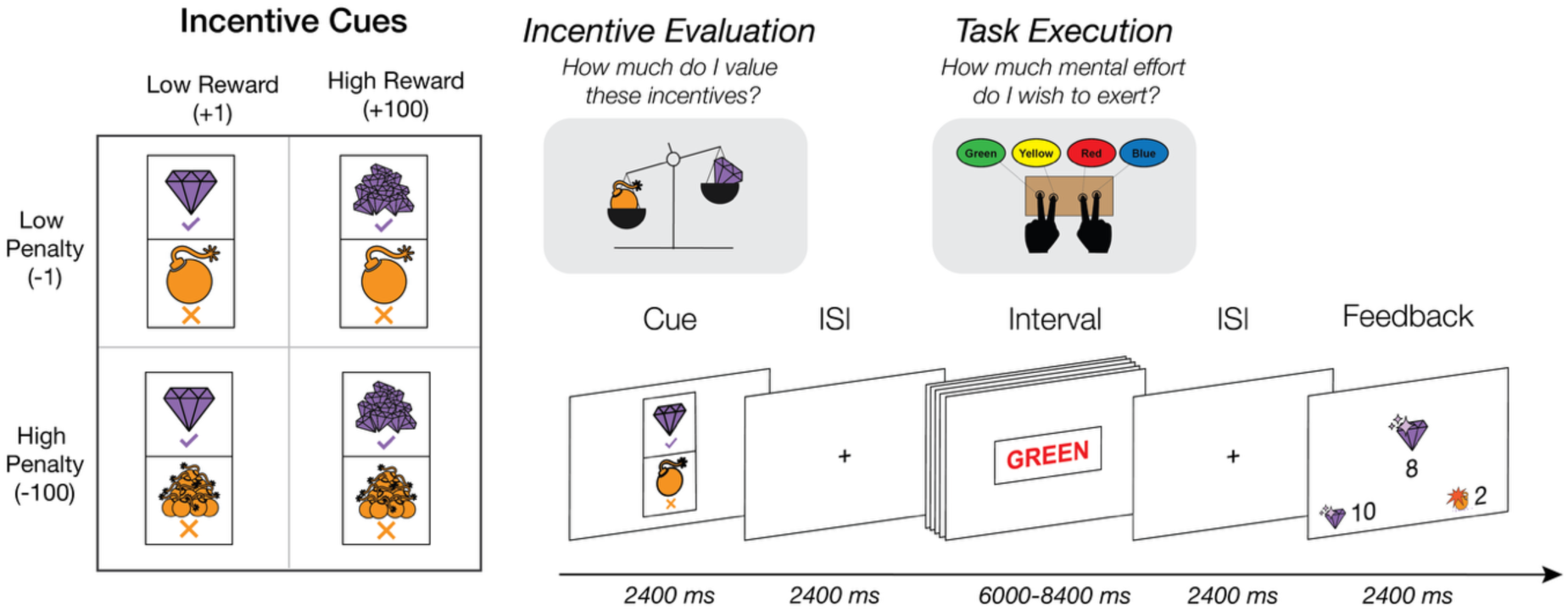
Multi-Incentive Control Task. At the beginning of each self-paced interval (incentive evaluation phase), a visual cue indicates the potential bonus to be earned for each correct response (e.g., earn 1 vs 100 points) and potential loss penalized for each incorrect response (e.g., lose 1 vs. 100 points). During the interval (task execution phase), the participant can decide how many Stroop trials they would like to complete and could choose to respond more efficiently (faster, maintain accuracy) or more cautiously (slower, increase accuracy), depending on how much monetary bonus they would like to earn vs. avoid losing. At the end of each interval, participants are given feedback about their net accrued points (e.g., gems earned minus bombs detonated). The length of the task execution phase was jittered (6000-8400 ms) to ensure that participants could not anticipate the duration and to increase design efficacy by decorrelating intervals.

### Brain networks for incentive evaluation and control allocation

To identify neural activity associated with incentive processing during cue presentation, we first focused on cued information that was novel for a given interval (Figure 2a). Since we held either reward or penalty level constant for a given interval (thereby making that information predictable prior to cue presentation), we examined the relationship between cue-related neural activity and incentive levels that varied over the course of a given block (e.g. high vs. low reward for blocks where penalty level was held constant). Examining cue-related activity within our *a priori* ROIs, we found that caudal dACC (t=3.46, p=.001) and VS (t=2.63, p=.009) selectively signaled higher levels of expected reward, and rostral dACC showed a non-significant trend towards signaling higher expected penalty (t=1.39, p=0.165). When collapsing across fixed and varying blocks, we found similar patterns in many of these regions (Supplemental Tables S1-S6, Figures S1-S2), with even stronger effects of expected penalty appearing in IFG (t=2.85, p=.004), DLPFC (t=2.61, p=.009), rostral dACC (t=2.14, p=.032), AI (t=2.03, p=.042), and VS (t=3.02, p=.003), suggesting that incentive levels are processed during cue presentation in these regions regardless of fixed or varying context. Several regions showed interactions between reward and penalty, with AI showing greatest sensitivity to low reward cues during high penalty conditions (t=-2.51, p=.012), caudal dACC showing greatest sensitivity to high reward cues during low penalty conditions (t=-2.86, p=.004), and DLPFC showing greater modulation during low reward compared to high reward incentive conditions (t=-2.79, p=.005).

**Fig 2.**
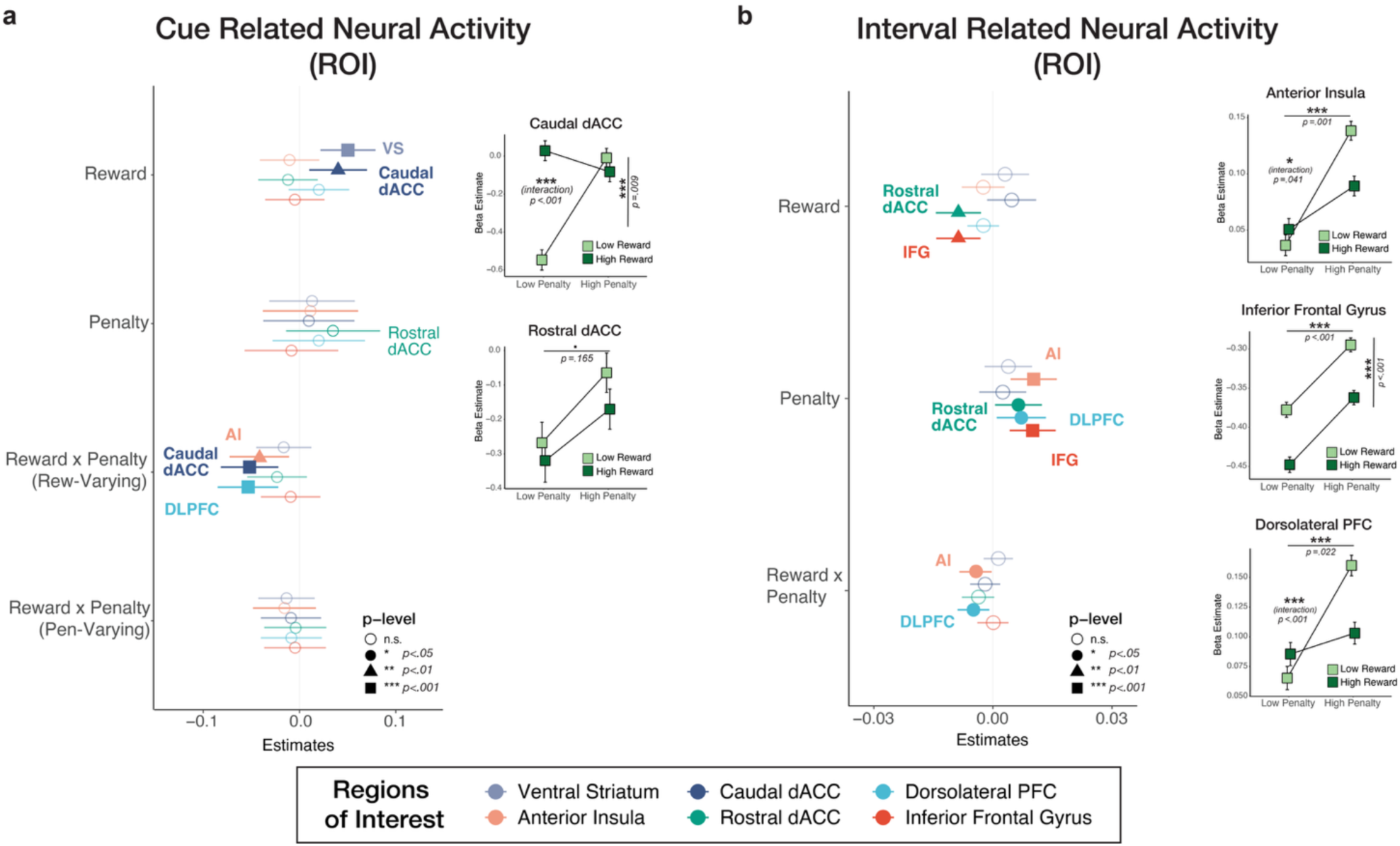
Cue and Interval Related ROI Activity. **(A)** ROI activity during cue phase of the task for reward-varying and penalty-varying trials, as incentive effects were most prominent in intervals when incentive level varied. Caudal dACC and VS selectively signaled higher levels of expected reward, whereas rostral dACC showed a trend towards signaling higher levels of expected penalty. Reward-by-penalty interactions revealed that Caudal dACC was sensitive to high reward cues during low penalty conditions, DLPFC was more modulated by low reward cues compared to high reward cues, and AI was sensitive to low reward during high penalty conditions. **(B)** ROI activity during the interval phase. IFG and rostral dACC both signal higher penalty and lower reward. DLPFC and AI both signal high penalty, and reward-by-penalty interactions in both regions show greater sensitivity to low reward during high penalty conditions. Error bars in the forest plot represent 95% CI, and error bars in the scatterplots within each forest plot represent standard error.

Distinct patterns of incentive processing emerge during task execution (interval phase). Examining neural activity across all intervals (Figure 2b), we find that both IFG and rostral dACC show increased activity during low-reward (vs. high-reward) intervals (IFG: t=-3.05, p=.002, rdACC: t=-2.99, p=.003) and during high-penalty (vs. low-penalty) intervals (IFG: t=3.39, p=.001, rdACC: t=2.14, p=.032). Higher penalty was also associated with increased interval-related activity in DLPFC (t=2.27, p=.023), and AI (t=3.46, p=.001). We found reward-by-penalty interactions in DLPFC (t=-2.41, p=.016), and AI (t=-2.05, p=.041), with sensitivity to low reward cues being enhanced during high penalty intervals (Figure S3). These analyses collapse across fixed and varying blocks because there was no *a priori* reason to restrict them (since both incentive attributes would be known at this stage), but supplementary analyses show that effects are qualitatively similar when restricting to varying blocks only (Supplemental Tables S1-S6).

Exploratory whole-brain analyses corroborated our ROI analyses, revealing similar patterns of neural activity during cue presentation and task execution (interval). With respect to cue-related neural activity, higher levels of reward were associated with greater activity in caudal dACC, whereas higher levels of penalty were associated with greater activation in rostral dACC. We also observed reward-related activations in the ventral striatum (consistent with ROI analyses), and overlapping activation for both incentives in regions of occipital and ventral temporal cortex (e.g., V4). During the intervals, we only observed reward-related activation in dACC, lateral prefrontal cortex, anterior insula, as well as in regions of occipital and ventral temporal cortex. We observed overall greater activation for reward compared to penalty incentives during both cue and interval phases, with greater activation in visual cortex during the cue phase and greater activation for reward in the left lateral prefrontal cortex and motor cortex during the interval phase (Supplemental Figure 4).

Interval-related neural activity also tracked performance during a given task interval, with faster RT associated with increased activity in supplementary motor area (SMA) and pre-SMA; slower RT associated with increased activity in regions of medial prefrontal cortex (including dACC) and lateral prefrontal cortex, as well as temporal and parietal cortices. Higher accuracy associated with increased activity in ventromedial prefrontal cortex, posterior parietal cortex, as well as in occipital and temporal cortices (Supplemental Figures S5-S6). Unless otherwise specified, all reported activations survived whole-brain correction using threshold-free cluster enhancement^23^ with familywise error correction at p<.05.

### Neural activity predicts distinct control strategies for mental effort allocation

Consistent with previous studies using this task^5,6^, we found that reward and punishment promoted distinct effort profiles. Higher levels of expected reward were associated with more efficient responding, reflected in faster responses (t=-7.47, p<.001) and moderate decreases in accuracy (t=-7.38, p<.001), ultimately achieving a higher number of correct responses per second (t=6.98, p<.001). By contrast, higher levels of expected punishment were associated with more cautious responding, reflected in slower (t=7.28, p<.001) and more accurate responding (t=11.75, p<.001), ultimately achieving fewer correct responses per second (t =-5.95, p<.001). Fitting task behavior to a drift diffusion model, we also replicated our previous finding that these two types of incentives are linked to dissociable control strategies: higher levels of expected reward are selectively associated with increased drift rate (potentially reflecting selective attention) and decreased threshold, whereas higher levels of expected punishment are selectively associated with increased threshold (Figure 3; Supplemental Tables S7-S9).

**Fig 3.**
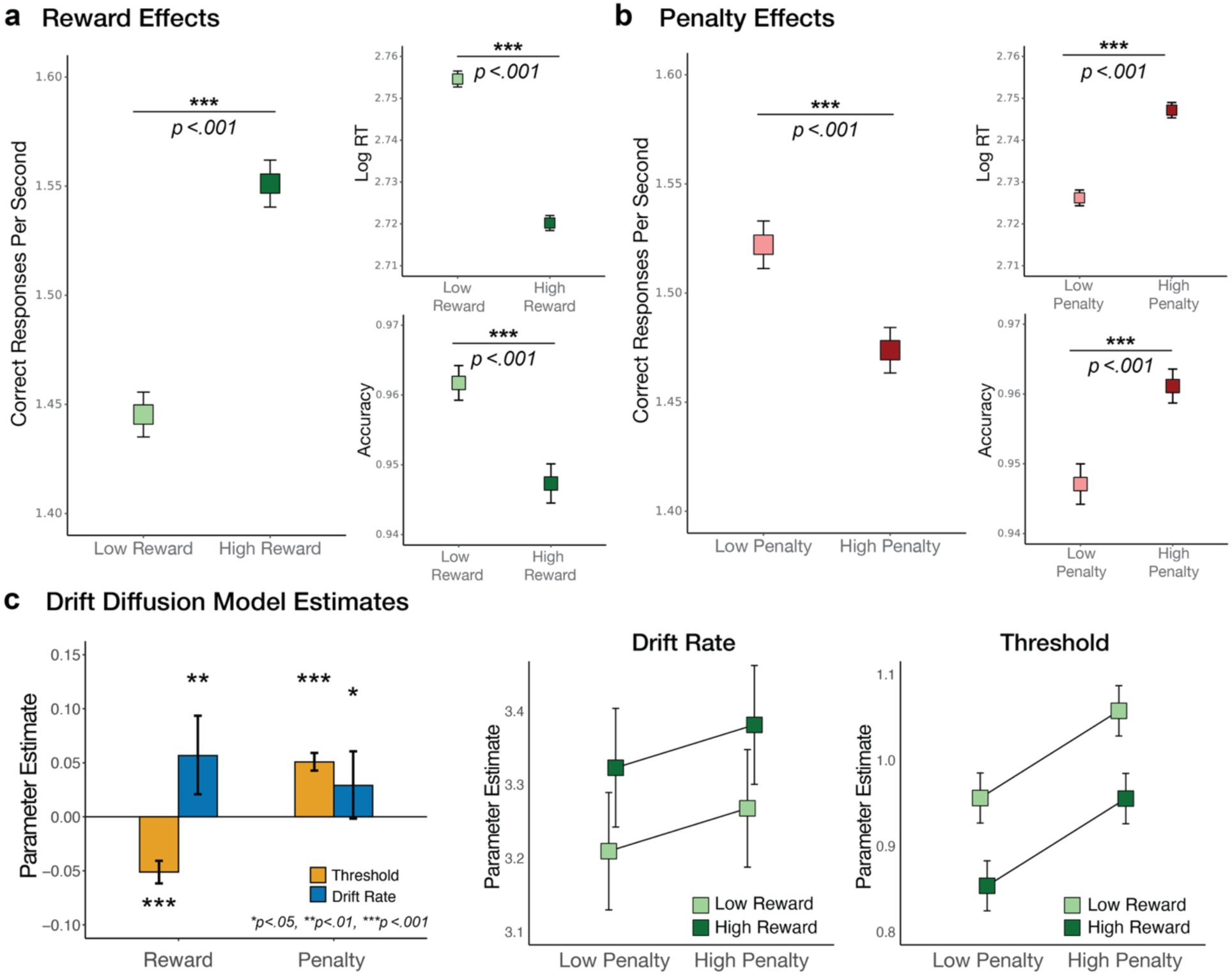
Behavior. **(A)** Reward enhanced response efficiency, reflected in faster responding and moderate decreases in accuracy. **(B)** Penalties promoted response caution, reflected in slower but more accurate performance. Error bars for behavioral measures represent 95% confidence intervals. c) Empirical model fits were consistent with normative model predictions: reward-related effects were associated with increased drift rate and reduced threshold, whereas penalty-related effects were associated with increased threshold (and to a less extent increased drift rate). Error bars of the DDM parameter estimates represent 95% credible intervals.

This dissociation in control signals engaged by distinct incentives enabled us to examine the extent to which different brain regions are selectively associated with specific incentives, control strategies, or both. To identify links between our *a priori* regions of interest and putative control strategies, we first examined neural activity during cue presentation across all trials, extracting beta estimates for each cue irrespective of the incentive combination. These regions dissociated across control configurations that guided predictions about the forthcoming interval. Specifically, caudal dACC, VS, and DLPFC were more strongly associated with increases in drift rate (p’s<.001), whereas rostral dACC and IFG were more associated with increases in response threshold (rdACC: p=.048), though this was only at a trend level in IFG (p=.066).

We found a different set of associations when examining neural activity during the task interval. IFG was again associated with higher threshold (p=.003) and VS with higher drift rate (p<.001). However, both VS and caudal dACC were now associated with lower threshold (VS: p<.001, caudal dACC: p=.034), and AI, rostral dACC, and DLFPC were associated with lower drift rate (p’s<.003). While these findings suggest that these regions may align with different control configurations at different stages of the task, it is also important to note that interval-related neural activity (unlike cue-related activity) can also reflect performance monitoring, such that neural signatures of lowered control (e.g., lower drift rates) can also reflect signatures of performance decrements (and/or corresponding control adjustments). Posterior estimates for the DDMs with cue and interval neural regressors are shown in Figure 4, Supplemental Tables S10-S21, and Figure S7.

**Fig 4.**
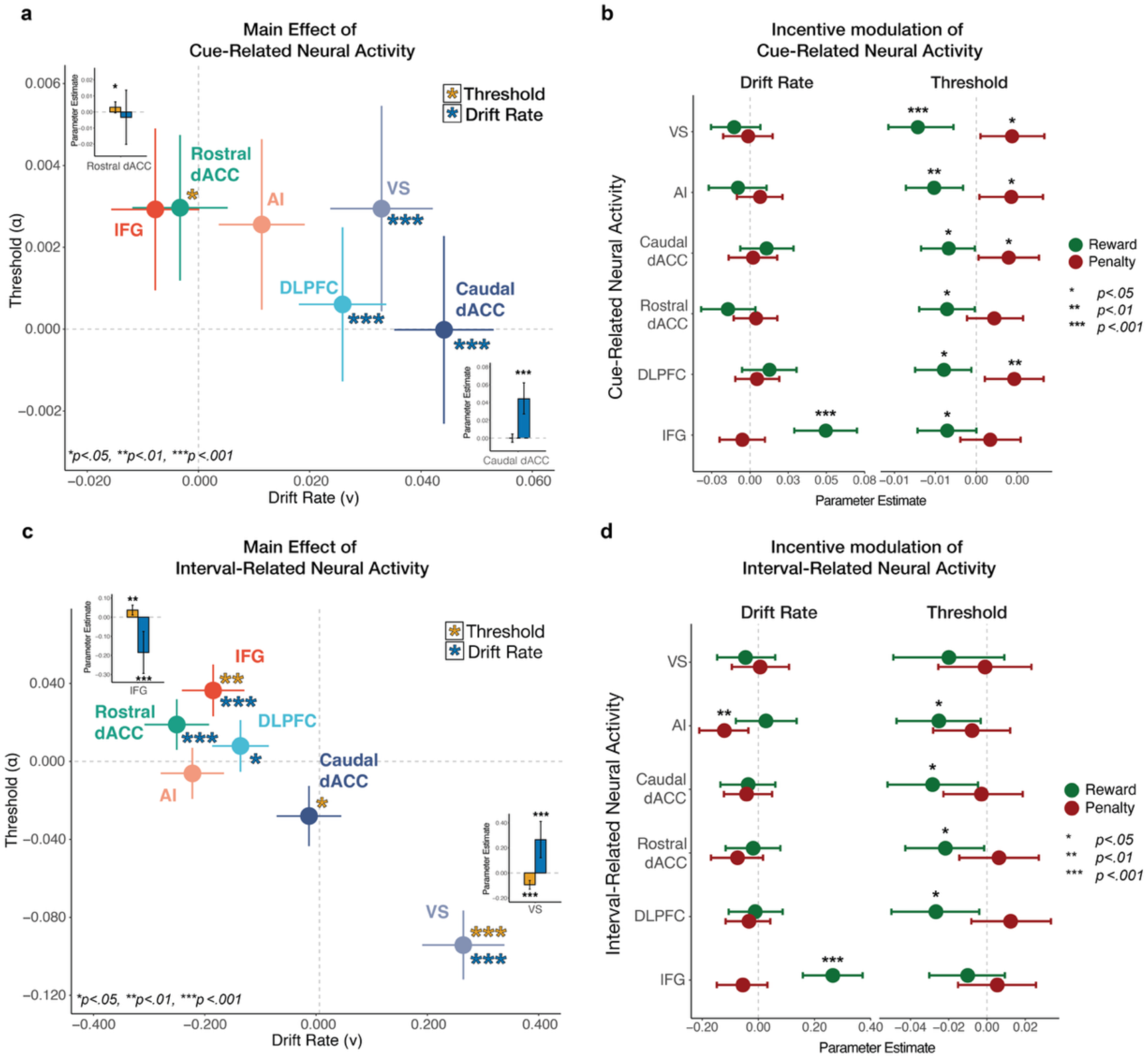
Model-based fMRI analyses. **(A)** Posterior estimates of main effects of cue-related neural activity on drift rate and threshold. VS, caudal dACC, DLPFC were associated with increased drift rate (p’s<.001), whereas rostral dACC was associated with increased threshold. **(B)** Posterior estimates of interactive modulation of cue-related neural activity on drift rate and threshold. We observed a specific modulatory effect of reward on IFG activity to promote increased drift rate. We observe a more general modulatory effect of both incentives on cue-related activity to promote greater adjustments in decision threshold (e.g., reward promotes lower threshold, penalty promotes higher threshold). **(C)** Posterior estimates of the interval-related neural activity on drift rate and threshold. VS was associated with increased drift rate and lowered threshold (p’s<.001), and IFG was associated with increased threshold and lowered drift rate (p’s<.01). AI, rostral dACC, and DLFPC were associated with lowered drift rate (p’s<.01), and caudal dACC was associated with lowered threshold (p<.05). **(D)** Posterior estimates of interactive modulation of interval-related neural activity on drift rate and threshold. We observed a specific modulatory effect of penalty on interval-related AI activity to lower drift rate. We observe a general modulatory effect of reward on interval-related activity to lower threshold in AI, Caudal dACC, Rostral dACC, and DLPFC. Error bars in the scatterplots represent standard deviation, and error bars in the barplots within the scatterplots represent 95% credible intervals. Error bars in the forest plots of the incentive x ROI interactions represent 95% credible intervals.

A separate set of analyses confirmed that the relationships we observed between neural activity and model parameters broadly mirror the patterns observed when relating neural activity to model-agnostic patterns of behavior (i.e., variability in accuracy and RT; Supp Tables S22-S27).

### Selective and global influences of motivation on neural mechanisms of adaptive control

Having established that our ROIs were engaged in both incentive processing and downstream control strategies, we next asked whether these processes converged within the same region. Specifically, we tested whether neural activity in each region modulated the effect of reward and penalty incentives on drift rate and threshold (cf. Figure 4b, 4d). We anticipated two possible modulatory relationships. First, if a region is involved in global task engagement or arousal, its activity should enhance the influence of *both* incentive types on their corresponding control strategies (e.g., strengthening reward’s influence on drift rate/threshold and penalty’s influence on threshold). Second, if a region preferentially encodes or supports control in response to a particular type of incentive, then its activity should selectively enhance the influence of that incentive (e.g., reward’s effect on drift rate/threshold) but not the other (e.g. penalty’s influence on threshold).

Focusing first on cue-related activity, we observed a global motivation effect across nearly all our regions of interest: increased activity predicted stronger reward-related decreases in threshold (p’s<.05) and stronger punishment-related increases in threshold (p’s<.05 in all except IFG and rostral dACC, which demonstrate non-significant trends in the same direction) (Figure 4b). We also found that IFG activity was selectively associated with enhanced reward-related increases in drift rate (p<.001), a relationship that we had not predicted. During the interval, we found that IFG again enhanced the relationship between reward and drift rate (p<.001), and that caudal dACC, rostral dACC, DLPFC, and AI enhanced the relationship between reward and decreased threshold (p’s<.05) (Figure 4d). We also found that the relationship between punishment and decreased drift rate was enhanced by activity in AI (p=.002), and to a lesser extent rostral dACC (p=.058) and IFG (p=.118), though this is not a directional relationship anticipated by our normative predictions or empirical findings. As in the case of the reward-related associations during the interval, it is important to again note that these relationships may reflect consequences of control allocation (e.g., enhanced signals of poor performance when expected punishment is high) rather than the control adjustments themselves.

### Subjective evaluation of motivational incentives predicts adaptive control

We have so far identified neural and behavioral signatures of reward and punishment within individuals. However, these analyses do not fully capture variability in motivational processes elicited by these incentives, as participants also differed substantially in how affectively salient or motivating they found these incentives. To capture this subjective variability, participants rated the pleasantness, arousal, motivation, effort, attention, and difficulty associated with each of the four incentive conditions (e.g., high/low reward/punishment) at the end of the session.

As in previous behavioral studies using this task^5,6^, we found that participants experienced high reward conditions as more pleasant (t=22.92, p<.001) and high punishment conditions as less pleasant (t=-13.59, p<.001). Self-reported arousal increased when either incentive was high (t*_Rew_*=4.95, p<.001; t*_Pen_*=5.68, p<.001) (Supplemental Table S29). Projecting these estimates onto a two-dimensional affective circumplex^21,22^ (Figure 5a), we find that higher rewards were associated with higher positive arousal (t=16.12, p<.001) and higher punishment with higher negative arousal (t=14.64, p<.001; Supplemental Table S31). Following previous work^6^, we averaged ratings of motivation, effort, and attention into a single ‘motivation index’. This motivation index reflected an interaction between reward and punishment (t=-11.02, p<.001), with motivation levels lowest for the low-reward/low-punishment condition, and motivation levels similarly high for all three other conditions (Figure 5b). Also consistent with previous results, despite task difficulty (e.g., proportion of congruent vs. incongruent trials) being held constant across conditions, participants rated conditions involving higher expected punishment as subjectively more difficult (t=6.85, p<.001; Supplemental Table S29; Figure S5).

**Fig 5.**
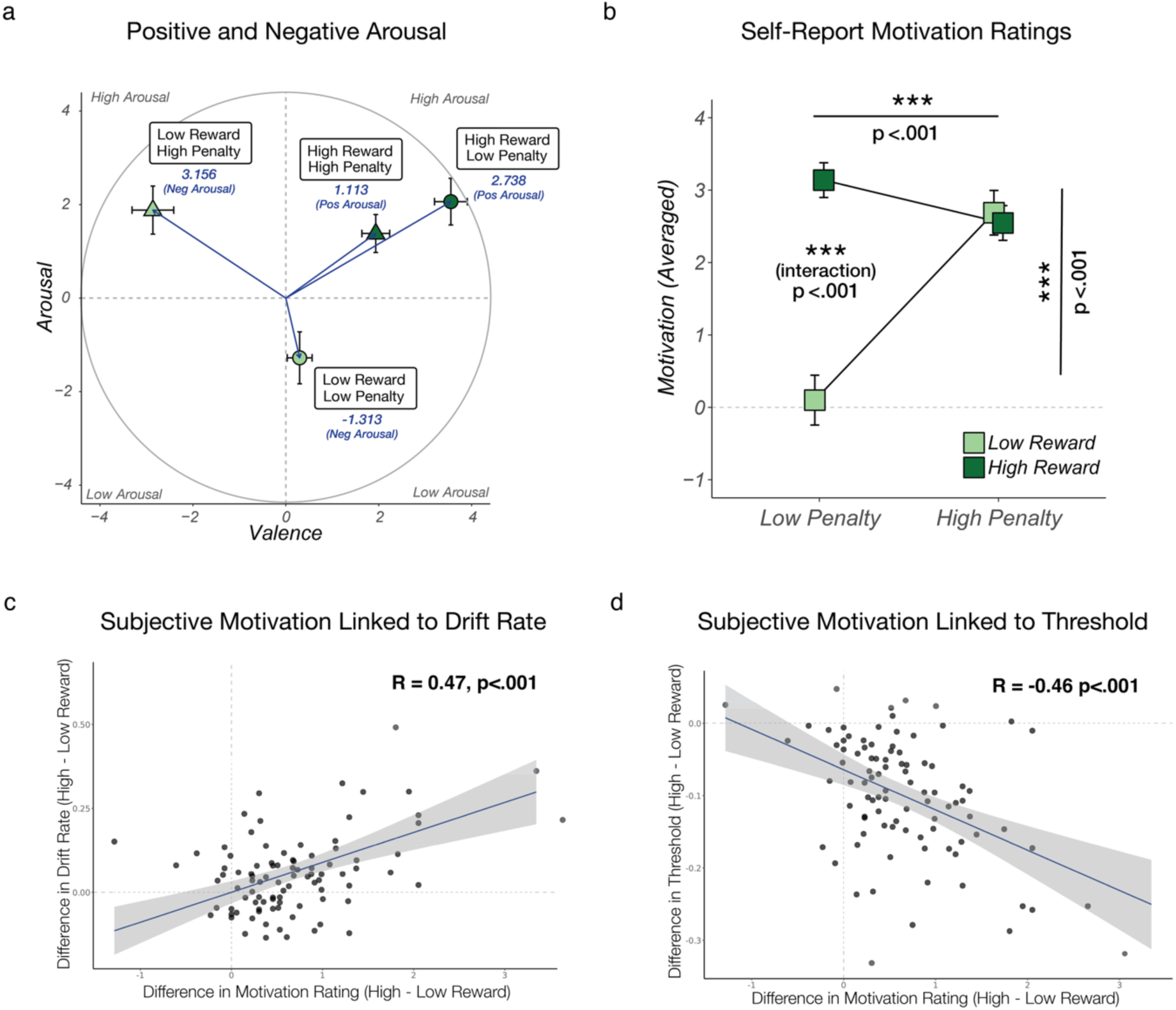
Self-report affect and motivation ratings. **(A)** Positive and Negative Arousal estimates for incentive cues. Participants found high reward cues generally positive and arousing, and showed greater differentiation in the low reward cues, with low reward / high penalty cues as more arousing and more negative whereas low reward / low penalty cues were less arousing and slightly positive. **(B)** Pleasantness, arousal, motivation (averaged across motivation, attention, and effort), and difficulty ratings by reward and penalty cues. Participants reported higher pleasantness, arousal, and motivation for high reward conditions. They reported lower pleasantness for high penalty conditions, as well as higher arousal, motivation, and difficulty. Significant interactions revealed these differences were greatest for pleasantness during high penalty conditions, and for arousal and motivation during low penalty conditions. The significant interaction in difficulty revealed that subjective difficulty was more modulated for low reward (relative to high reward) conditions. Error bars for the self-report ratings represent 95% confidence intervals. **(C,D)** Individuals who reported greater differentiation in subjective motivation for high vs. low reward also demonstrated greater degree of reward modulation of both drift rate and threshold (i.e., greater influence of reward on adjustments in adaptive control). P-values are Benjamini-Hochberg corrected.

We found that individual differences in self-reported reward motivation predicted how participants adjusted control in response to rewards. Participants who reported larger motivation difference between high and low reward also showed greater increases in drift rate (R=0.47, p<.001; Benjamini-Hochberg corrected; Figure 5c) and greater decreases in threshold (R=-0.46, p<.001; Benjamini-Hochberg corrected; Figure 5d). By contrast, self-reported motivation for penalty incentives was not associated with threshold adjustments, although participants who were more penalty-motivated exhibited higher thresholds overall (R=-.35, p=.007; Benjamini-Hochberg corrected; Supplementary Table S33, Figure S9).

### Subjective evaluation of motivational incentives predicts neural activity

Having observed differentiation in arousal and motivation at the self-report level, we next examined whether these subjective states were captured in neural variability across participants. These analyses allowed us to test whether incentive-driven experiences map onto the neural processes that support adaptive control.

Positive arousal was associated with activity in reward-related regions. At cue onset, higher positive arousal corresponded to greater activity in VS (t=2.06, p=.040) and caudal dACC (t=2.08, p=.041; Figure 6a; Supplemental Table S33). During the interval, it remained positively associated with VS (t=2.05, p=.005) but was negatively associated with activity in rostral dACC (t=-2.65, p=.008) and IFG (t=-2.82, p=.005; Figure 6b; Supplemental Table S34). We additionally observed interactions between positive arousal and reward in caudal dACC (t=2.20, p=.028), rostral dACC (t=2.58, p=.010), and IFG (t=1.96, p=.050), as well as between positive arousal and penalty in VS (t=3.11, p=.002), rostral dACC (t=2.02, p=.044), and IFG (t=3.26, p=.001). Negative arousal was associated with greater cue-related activity in VS (t=2.93, p=.003) and greater interval-related activity in IFG (t=4.28, p<.001), rostral dACC (t=4.31, p<.001), AI (t=3.79, p<.001), DLPFC (t=3.20, p=.001), and VS (t=2.44, p=.015). Incentive interactions further revealed that the relationship between negative arousal and IFG differed under high vs. low reward conditions (t=-2.05, p=.041), whereas the relationship between negative arousal and all ROIs were broadly enhanced under penalty (t’s<-1.95, p’s<.048).

**Fig 6.**
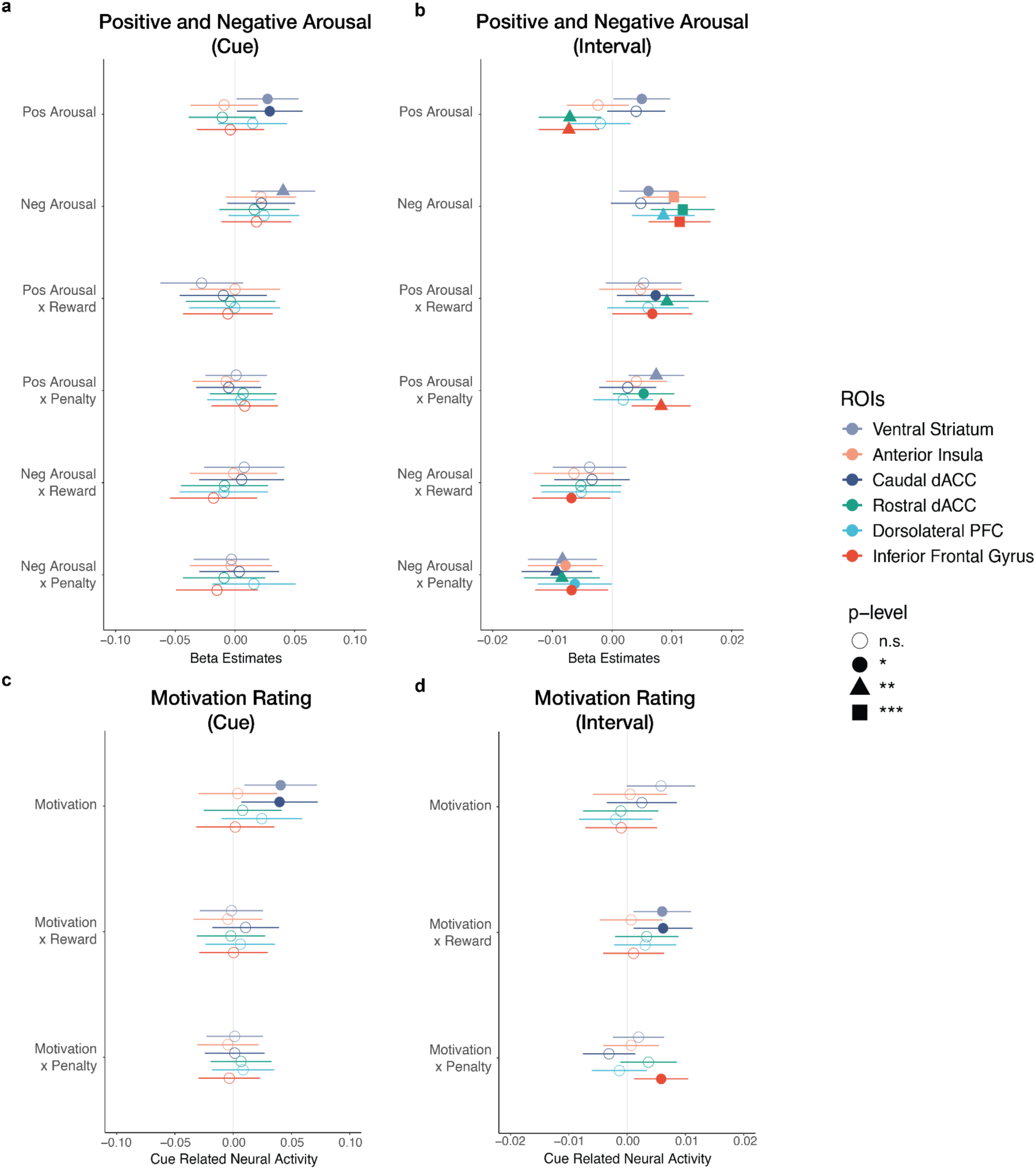
Self-report ratings predict neural activity. Linear mixed models tested whether neural activity were predicted by positive arousal, negative arousal, and motivation ratings (Supplemental Tables S33-S36). Forest plots show beta estimates with 95% confidence intervals. **(A)** At cue onset, caudal dACC predicted positive arousal, and VS predicted both positive and negative arousal. **(B)** During the interval, VS tracked positive arousal, whereas rostral dACC and IFG tracked higher negative and lower positive arousal; AI and DLPFC only tracked negative arousal. Reward interacted with caudal dACC, rostral dACC, and IFG, whereas penalty interacted with VS, rostral dACC, and IFG in predicting higher positive arousal. **(C)** Cue-related VS and caudal dACC activity predicted higher motivation. **(D)** During the interval, interactions between reward and VS and caudal dACC predicted higher motivation, and interactions with penalty and IFG predicted higher motivation.

Higher ratings of motivation were associated with greater cue-related activity in VS (t=2.56, p=.010) and caudal dACC (t=2.23, p=.015) (Figure 6c; Supplementary Table S36), consistent with findings reported above (Figure 5a) suggesting that these regions at minimum reflect the global motivation to perform the task. Motivation level was associated with greater reward-related activity in VS (t=2.39, p=.017) and caudal dACC (t=2.40, p=.017), and greater penalty-related activity in IFG (t=2.46, p=.014), suggesting that motivation level is predictive how responsive these regions are to reward and penalty conditions (Figure 6d; Supplementary Table S37).

Although we did not have *a priori* predictions about the relationship between subjective difficulty and neural activity, for completeness we tested for associations with this variable. We found that ratings of subjective difficulty were positively associated with cue-related activity in VS (t=3.99, p<.001), AI (t=2.68, p=.007), caudal dACC (t=2.09, p=.037), rostral dACC (t=2.38, p=.018), and DLPFC (t=2.72, p=.007), as well as with interval-related activity in AI (t=3.47, p=.001), IFG (t=2.65, p=.008), DLPFC (t=2.99, p=.003), VS (t=2.19, p=.028), and caudal dACC (t=2.15, p=.032; Supplemental Tables S37-S38, Figure S10). Additional analyses with pleasantness and arousal ratings are shown in Supplemental Tables S39-S42 and Figure S10.

## Discussion

In the current study, we investigated the neurocomputational mechanisms that underlie the influence of reward and punishment on mental effort allocation. Whereas past research has shown that cognitive control–the ability to process information to support adaptive goal-directed behavior–is modulated separately by reward and punishment,^24–27^ the utilization our recently developed Multi-Incentive Control Task combined with our reward-rate optimization model^5,6^ provided a unique opportunity to illuminate *how* distributed neural circuits are organized when both reward and penalty incentives are used to exert dissociable influences on one’s adaptive control strategy.

We identified a network of brain regions sensitive to evaluation of reward and penalty incentives and observed distinct neural regions that may underlie their differing effects on control allocation. Our neural dissociation by valence is consistent with past work showing ventral striatum (VS) signaling associated with reward-related enhancements vs. anterior insula (AI) signaling associated with penalty-related enhancements in decision-making.^28–31^ Moreover, our finding that dorsal anterior cingulate cortex (dACC) and lateral prefrontal cortex regions are associated with incentive-related control adjustments are consistent with past research highlighting regions as modulated by reward and punishment.^32–36^ Interestingly, although the neural patterns underpinning interactions between motivation and cognitive control networks are still somewhat inconsistent, our subdivision of these frontal regions (caudal vs. rostral dACC, dorsolateral prefrontal cortex (DLPFC) vs. inferior frontal gyrus (IFG)) illustrates one way in which these distributed neural networks are systematically modulated and organized by positive vs. negative incentive valence.

Our model-based fMRI analyses revealed two distinct neural profiles underlying the dissociable adaptive control strategies based on our normative model (efficiency vs. caution). We observe a consistent pattern across caudal dACC, ventral striatum (VS), and dorsolateral prefrontal cortex (DLPFC), in which cue-related activity predicted increased drift rate. These regions may be upregulated during the cue phase to promote efficient responding when expected reward is high, which is the optimal strategy according to our normative reward rate model.^37^ Conversely, cue-related activity in rostral dACC predicted an increased decision threshold, revealing that this latter region may be linked in upregulating the strategy for cautious responding when expected penalty is high. Intriguingly, this dissociation is still prominent with respect to interval-related activity of VS and IFG (but not dACC), revealing putative downstream regions involved with the execution and/or monitoring of corresponding control strategies.

A key feature of our task was the inclusion of bundled reward and penalty incentives, which allowed us to evaluate whether these brain regions represent the *specific* influence of given incentive, or instead represented the *global* influence of both reward and punishment on cognitive control allocation. We find evidence that both incentives interacted with cue-related activity in all our *a priori* ROIs to promote incentive-relevant control strategy adjustments (e.g., reward lowering decision threshold, penalty increasing decision threshold). One caveat is that is although penalty-related increases in threshold did not reach statistical significance in rostral dACC and IFG (though these effects were in the same direction as predicted by the model), is it possible we observed marginal interactive effects of penalty on threshold given the already strong main effect of both these regions on increased threshold (see Fig 4a). We also observed that interval-related neural activity in caudal dACC, rostral dACC, DLPFC, and AI interacted with reward to lower threshold, which can reflect modulation of the engaged control signals themselves or monitoring of resulting performance (e.g., slow/error-prone responding) and/or implications for cumulative expected reward.^38^ Additionally, we observed a specific interaction between reward and IFG (during cue and interval) resulting in increased drift rate, as well as an interaction between penalty and AI (during interval) resulting in lowered threshold, though these effects are difficult to interpret as they were not specifically predicted.

We also found a relationship between our behavioral effects and self-reported affect and motivation, revealing important insight into how motivational and affective states modulate (or are modulated by) neural activity. For instance, associations with cue-related neural activity may reflect a proactive influence, (e.g., greater arousal or motivation), whereas associations with interval-related activity may reflect a retrospective experience shaped by one’s performance during that task interval (e.g., positive arousal may be associated with the excitement of earning reward for performing well, negative arousal may be associated with the fear of being penalized for mistakes).^39,40^ We observed a distinction in cue-related activity between caudal dACC, which was sensitive to the positively valenced cues, and VS, which was sensitive to higher-incentive cues regardless of valence. Both caudal dACC and VS were also associated with higher motivation, consistent with prior work showing convergence between incentive-related modulation and subjective evaluation of motivational state during effort allocation.^35^ In contrast, interval-related activity showed broader associations with both positive and negative arousal, as well as interactions with incentives. Rostral dACC and IFG were more strongly associated with higher negative arousal and lower positive arousal, consistent with past work linking both regions to negative affect.^41–43^ We show that incentives strengthened the association between ROIs and positive and negative arousal, which suggest affective modulation of these ROIs by incentive cues. However, we cannot rule out the possibility that these effects may also be driven by introspective evaluation of task performance (e.g., ratings were collected at the end of the study). Future studies that account variability in these affective ratings during the task (e.g., within intervals) would clarify the temporal dynamics of affective modulation within these brain regions during adaptive control.

The observed associations between self-report motivation ratings task performance reveal an intriguing link between the subjective evaluation of incentives and incentive-related adjustments in control strategy. Participants who reported that they were more motivated by high (relative to low) rewards also adjusted showed greater adjustments in their control strategy (e.g., people increased their drift rate and lowered their threshold in response to higher reward value). This finding highlights reward-sensitivity as an important individual difference measure in cognitive control tasks, which provides an additional relevant dimension for understanding individual differences in cognitive control capacity.^44–47^ Future work accounting for how variability in both motivational and affective processes may interact with cognitive control ability will provide more comprehensive understanding of the repertoire of the possible ways that incentives shape adaptive behavior.^48^

Together, our results facilitate a deeper understanding of the neural circuits and computations that underlie how reward and penalty incentives elicit dissociable strategies for cognitive control allocation. Specifically, our finding that distinct brain networks associated with reward-related enhancement of response efficiency (e.g., caudal dACC, VS) vs. penalty-related enhancement of response caution (e.g., rostral dACC, IFG) reveals novel insight into the neural mechanisms that underlie incentive-modulated adaptive control. Our functional division of caudal and rostral dACC subregions through distinct resting-state network affiliations^49–52^ contributes to our broadening understanding of dACC’s putative role in dynamically integrating information and transmitting downstream signals about the type of adaptive control.^53–60^ However, it remains unclear to what extent these distinct neural profiles are maintained when outcomes are no longer performance-contingent (i.e., points or money are no longer yoked to correct or incorrect responses or are probabilistically determined). Past work has shown that one’s performance efficacy interacts with reward to modulate neural signals, yet how one’s effort is altered in stressful or uncontrollable environments, or alternatively in the face of diverse types of aversive outcomes (e.g., monetary loss, physical pain, unpleasant sounds, social rejection) remains unclear.^24,61–63^ Future work that delineates the motivational context of incentives (i.e., whether an incentive promotes reinforcement or punishment, and/or whether the outcome is controllable or not) as well how these neural patterns map onto with well-validated neural signatures of affect, motivation, and cognitive control will help us glean further insight into how cognitive control networks are activated by diverse motivational incentives (e.g., money, pain, threat).^64,65^

Finally, while our study provides novel insight into the neural signals that underlie the normative assumption of reward rate maximization from the EVC model, there are several limitations to the interpretation of our results. For instance, it is possible that these control strategies may not only reflect one’s desire to pursue extrinsic incentives (i.e., monetary reward) but rather one’s internal state. Recent theoretical frameworks have suggested that motivated behavior serves to promote anticipated affective states, with individuals gravitating towards situations that evoke positive affect and avoiding those that evoke negative affect.^66,67^ Our data are consistent with these theoretical accounts, revealing how these same neural circuits that predict behavior also predict affective processes (e.g., positive and negative arousal, motivation). Future work should seek to examine how adaptive changes in motivated behavior unfold in more naturalistic settings in which potential drivers of positive and negative affect change, and/or are modulated by more ecologically relevant incentives commonly used in human and animal paradigms (e.g., electric shocks, food, or drink). Tackling such questions would further benefit from examining the interoceptive states associated with the physiological signatures of motivated behavior (e.g., hunger, satiety, stress, arousal).^68,69^ Ultimately, this multimodal combination of naturalistic and ecological experimental paradigms ^70^with theoretically-informed neurocomputational models (alongside other real-world longitudinal data) would be a crucial step towards translating computational insights to predict motivational and affective impairments in neuropsychiatric disorder.^71–74^

## Materials and Methods

### Experimental Procedures

All demographic and self-report data were collected and managed using Qualtrics, a secure web-based application hosted at Brown University. Prior to the beginning of the study, participants were screened by a researcher and determined to be eligible for the study if they were currently enrolled at a four-year college/university in the Rhode Island area, never transferred colleges/universities, lived consecutively in the United States for greater than 15 years, and did not have color blindness or dyslexia. Participants were additionally screened for MRI-specific criteria to determine whether they were safe to enter the MRI environment. Once participants were enrolled in the study, they completed several Qualtrics surveys to provide information about their home environment, upload a student transcript, and answer several self-report questionnaires about their childhood environment, personality measures, clinical measures, and COVID impact. More details about the questionnaires are detailed below.

During the scanning session, participants arrived and were instructed to practice the task they would perform in the MRI. Participants first completed several practice blocks of the Multi-Incentive Control Task (MICT) in a testing room. They were told that they could earn a monetary bonus in the MRI based on their performance and completed four practice blocks with the different combinations of bundled cues. Next, participants were prepared to enter the MRI scanning environment and were instructed on how to use the button box (4 Button Curve Right Fiber Topic Response Pad). Participants performed eight blocks of the task in the MRI (about 1.5 hours). After the scan, participants completed a Qualtrics survey and provided Likert ratings of the cue types (e.g., low reward/low penalty, low reward/high penalty, high reward/low penalty, high reward/high penalty). Finally, participants were debriefed and compensated for completing the study plus their monetary bonus up to $20 based on their task performance.

### Participants

A total of 100 college-student participants (58 female; 18-23 years, mean=20.1, SD=1.25) with normal or corrected-to-normal vision participated in the experiment. All participants provided written consent approved by the Brown University Institutional Review Board and received payment for their participation ($10 per hour for completing the Qualtrics survey at home, $15 per hour for the MRI session), plus additional earnings of up to $20 based on task performance. One participant was excluded from the analyses because of a technical error during data collection. Another 5 participants were excluded because of model convergence issues during fMRI preprocessing, limiting our ability to make inferences about their data. The final analysis sample consisted of 94 participants (55 female; 18-23 years, mean=20, SD=1.24).

### Multi-Incentive Control Task (MICT)

Participants performed a self-paced incentivized Stroop task where they could earn monetary rewards (1 gem or 100 gems) for each correct response and were penalized with monetary losses (1 bomb or 100 bombs) for each incorrect response. A key feature of the experimental design was the self-paced nature of effort allocation, as participants could choose to complete as many or as few Stroop trials as desired within a time limited (6-8.4 seconds) interval. Incentive cues were bundled and randomly varied across intervals (e.g., within each block, one incentive value maintained while the other varied between high and low value), allowing parametric modulation of the expected value for reward and penalty cues on cognitive control allocation on an interval-by-interval basis.

### Self-Report Likert Ratings

After each MRI session, participants completed a post-scan survey where they rated the four incentive cues (low reward/low penalty, low reward/high penalty, high reward/low penalty, high reward/high penalty) on various dimensions on a scale from -5 to 5. They rated the cues based on their pleasantness, arousal, motivation, difficultly, effort, and attention.

**Table 1.** Self-Report Likert Ratings at end of Experimental Session.

| Dimension | Question |
| --- | --- |
| Pleasantness | How **unpleasant/negative** or **pleasant/positive** it made you feel |
| Arousal | How **relaxed/calm** or **stimulated/aroused** it made you feel |
| Motivation | How **MOTIVATED** you were |
| Difficulty | How **DIFFICULT** the task was |
| Effort | How much **EFFORT** you exerted |
| Attention | How much you **paid attention** |

### Behavioral Analyses

Task performance was analyzed at the level of a given interval and the level of responses to individual Stroop stimuli within that interval. For the interval-based analyses, the last trial of each interval was excluded. We analyzed participant’s interval-level performance by fitting a linear mixed model with maximum likelihood using the lme4^75^ and lmerTest packages^76^ in R to estimate the correct responses per second as a function of contrast-coded reward and punishment levels (High Reward=1, Low Reward=-1, High Punishment=1, Low Punishment=-1) as well as their interaction. The model controlled for gender, and proportion of congruent stimuli, interval length, interval session number, and included specified random effects for reward and punishment. Log-transformed reaction time trial-wise data (correct responses only) was analyzed with a linear mixed effects model, controlling for stimulus congruency, interval length, trial session number, and included specified random effects for reward and punishment. Accuracy trial-wise data was analyzed with a generalized linear mixed effect model, controlling for stimulus congruency, interval length, trial session number, and include specified random effects for reward and punishment. The random effects for each model were determined by a model fitting procedure starting with maximally random effect structure and performing backward stepwise elimination based on the change in log-likelihood^77^, using the buildmer^78^ package in R. To ensure model convergence, we used the *nlminb* optimization routine from the optimx^79,80^ package in R.

### Computational Modeling Analyses

#### Model fitting

We parameterized participants’ responses to on the Stroop task as correct (e.g., responding to the ink color) or incorrect (e.g., responding to the word) using the drift diffusion model (DDM), a sequential sampling model that characterizes speeded decision-making as a set of constituent processes that reflect evidence accumulation towards one of two boundaries^18,81^. Following previous work^5,6^, we only included automatic errors (e.g., incorrectly responding to the word during an incongruent trial), and excluded random errors (e.g., incorrectly responding to a color not associated with the color stimulus), as we are only interested in characterizing the decision about cognitive control allocation and sought to reduce noise during the model fitting process. This did not significantly impact the dataset, since random errors only consisted of 3.28% of total trials.

We performed hierarchical Bayesian estimation of DDM parameters (e.g., drift rate *v*, response threshold *a*, prepotent bias *z*) using the *HDDMregressor* function in the HDDM package^19,82^. We estimated the parameters of drift rate and threshold as a function of reward level (high reward = 1, low reward = -1), and penalty level (high punishment = 1, low punishment = -1). In our regression models, we accounted for the collapsing response deadline (i.e., decision threshold) associated with the reduced amount of time to complete a given trial within each interval^82,83^. For example, for a given 8-second interval, if the participant took 500 ms to respond to the first stimulus followed by a 250 ms interstimulus interval, then the participant would have 7250 ms left within the interval to complete additional trials within that interval. To approximate this interval-level collapsing bound in our threshold model, we included a variable called “scaled linear running time,” which tracked the amount of time that had passed with each interval (z-scored). To account for congruency effects, the model for drift rate included trial congruency (congruent = 1, incongruent = 0). We fixed the starting point at the mid-point between the two boundaries as there was no prior bias toward a specific response in the task and controlled for congruency. The non-decision time was fitted as a free parameter. The models additionally included intertrial variability in drift and non-decision time and included p_outlier to downweight the influence of extreme RTs. We ran 5 parallel chains for 10000 iterations each, discarded the first 7000 warm-up samples, and kept the remaining 3000 samples as posterior estimates. Model convergence was tested using the Gelman-Rubin convergence diagnostic R-hat^84,85^, which compares between-chain and within-chain variances for model parameters across the Markov chains (R-hats close to 1 indicates convergence; we considered a model successfully converged with R-hats <= 1.01). Model comparison was based on deviance information criterion (BPIC; lower is better) to identify the best model for the behavioral data.

#### Posterior Predictive Checks

Posterior prediction checks were performed to validate the model fitting procedure^86^. We generated 500 independent samples from the posterior distribution of fitted parameters and then simulated the reaction time distribution for each posterior sample. The predicted reaction time distribution matches with the actual reaction time distribution and error rate for each condition (Supplemental Figure S11).

### fMRI Data Acquisition

The MRI data were acquired on a 3-Tesla Siemens Prisma scanner equipped with a 64-channel head coil. A T1-weighed MPRAGE scan was acquired for each participant (TR = 2400 ms, TE = 1.86 ms, flip angle = 7 degrees, slices per slab = 192, thickness = 0.80 mm, FOB = 205 x 205 mm). For each EPI bold run (8 total), 360 volumes were acquired with 2.4 mm isotropic voxels (TR = 1200 ms, TE = 33.0 ms, flip angle = 62 degrees, 60 slices, interleaved series, slice thickness = 2.4 mm, FOV = 211 x 211 mm, acquisition matrix 88 x 88 yielding an in-plane resolution of 2.4×2.4 mm). Finally, a field map was acquired for each participant (TR = 588 ms, TE = 4.92 ms, flip angle = 60 degrees, 60 slices, interleaved series, slice thickness = 2.4 mm, FOV = 211 x 211 mm, acquisition matrix 88 x 88 yielding an in-plane resolution of 2.4×2.4 mm).

### fMRI Data Preprocessing

Preprocessing was performed using fMRIPrep versions 20.2.6^87,88^, which is based on Nipype 1.7.0^88,89^ and uses many internal operations of Nilearn 0.6.2^38^, mostly within the functional processing workflow.

#### Anatomical data processing

The T1-weighted (T1w) image was corrected for intensity non-uniformity (INU) with N4BiasFieldCorrection^90^, distributed with ANTs 2.3.3^91^ [RRID:SCR_004757], and used as T1w-reference throughout the workflow. The T1w-reference was then skull-stripped with a Nipype implementation of the antsBrainExtraction.sh workflow (from ANTs), using OASIS30ANTs as target template. Brain surfaces were reconstructed using recon-all^92^ (FreeSurfer 6.0.1, RRID:SCR_001847), and the brain mask estimated previously was refined with a custom variation of the method to reconcile ANTs-derived and FreeSurfer-derived segmentations of the cortical gray-matter of Mindboggle^93^ (RRID:SCR_002438). Volume-based spatial normalization to two standard spaces (MNI152NLin2009cAsym, MNI152NLin6Asym) was performed through nonlinear registration with antsRegistration (ANTs 2.3.3), using brain-extracted versions of both T1w reference and the T1w template. The following templates were selected for spatial normalization: ICBM 152 Nonlinear Asymmetrical template version 2009c^70^ [RRID:SCR_008796; TemplateFlow ID: MNI152NLin2009cAsym], FSL’s MNI ICBM 152 non-linear 6th Generation Asymmetric Average Brain Stereotaxic Registration Model^94^ [RRID:SCR_002823; TemplateFlow ID: MNI152NLin6Asym].

#### Functional data processing

For each of the 8 BOLD runs found per subject (across all tasks and sessions), the following preprocessing was performed. First, a reference volume and its skull-stripped version were generated using a custom methodology of fMRIPrep. A B0-nonuniformity map (or fieldmap) was estimated based on a phase-difference map calculated with a dual-echo GRE (gradient-recall echo) sequence, processed with a custom workflow of SDCFlows inspired by the *epidewarp.fsl* script and further improvements in HCP Pipelines^95^. The fieldmap was then co-registered to the target EPI (echo-planar imaging) reference run and converted to a displacements field map (amenable to registration tools such as ANTs) with FSL’s fugue and other *SDCflows* tools. Based on the estimated susceptibility distortion, a corrected EPI (echo-planar imaging) reference was calculated for a more accurate co-registration with the anatomical reference. The BOLD reference was then co-registered to the T1w reference using bbregister (FreeSurfer) which implements boundary-based registration^96^. Co-registration was configured with six degrees of freedom. Head-motion parameters with respect to the BOLD reference (transformation matrices, and six corresponding rotation and translation parameters) are estimated before any spatiotemporal filtering using mcflirt^97^ (FSL 5.0.9). BOLD runs were slice-time corrected to 0.551s (0.5 of slice acquisition range 0s-1.1s) using 3dTshift from AFNI 20160207^98^ (RRID:SCR_005927). The BOLD time-series (including slice-timing correction when applied) were resampled onto their original, native space by applying a single, composite transform to correct for head-motion and susceptibility distortions. These resampled BOLD time-series will be referred to as preprocessed BOLD in original space, or just preprocessed BOLD. The BOLD time-series were resampled into standard space, generating a preprocessed BOLD run in MNI152NLin2009cAsym space. First, a reference volume and its skull-stripped version were generated using a custom methodology of fMRIPrep. Automatic removal of motion artifacts using independent component analysis^99^ (ICA-AROMA) was performed on the preprocessed BOLD on MNI space time-series after removal of non-steady state volumes and spatial smoothing with an isotropic, Gaussian kernel of 6mm FWHM (full-width half-maximum).

### ROI selection

We selected regions of interest (ROIs) based on prior meta-analyses of motivation and cognitive control, the subjective valuation of positive and negative motivational incentives in decision-making contexts, and effort-based decision-making^13–15^. These included the ventral striatum (VS), anterior insula (AI), two subregions of dorsal anterior cingulate cortex (caudal vs. rostral dACC), dorsolateral prefrontal cortex (DLPFC), and inferior frontal gyrus (IFG). Below we describe how each of these ROIs were selected and/or extracted.

#### Ventral Striatum

We identified the bilateral ventral striatum ROI from the different fMRI dataset in the laboratory using a different task with similar cue and interval structure, and the CuexReward contrast was thresholded (cluster size k>25, p<.05) and masked by an anatomical atlas (AAL) of the striatum from the Bartra meta-analysis, and we subtracted voxels that overlapped with a bilateral insula mask (cluster size k>5, p<.20) from the Harvard Oxford cortical atlas (fsleyes version 1.6.1).

#### Anterior Insula

We identified a bilateral anterior insula ROI from the conjunction of positive and negative effects from Bartra meta-analysis, and was additionally thresholded (size k>25, p<.05).

#### Caudal and Rostral Dorsal Anterior Cingulate Cortex

To account for the functional heterogeneity of dACC^53–56^, we used an analysis mask that provided broad coverage across the medial frontal cortex (e.g., anterior cingulate cortex and preSMA).^100^ This mask was constructed by overlapping cingulate responses co-occurring with ‘cognitive control’ (Neurosynth uniformity test)^101^ and selected parcels from the whole brain Schaefer parcellation (400 parcels, 17 networks, 2mm voxels)^49,50^ that had a 50 voxel overlap with the meta-analytic mask. These parcels were categorized by membership in their corresponding functional networks (caudal dACC: salience attention network, rostral dACC: frontoparietal control network).

#### Dorsolateral Prefrontal Cortex

We identified the bilateral DLPFC ROI using a similar conjunction procedure as with the dACC, and we constructed this mask by overlapping dorsolateral prefrontal cortex responses and parcels from the Schaefer parcellation with a 50 voxel overlap with the meta-analytic mask.

#### Inferior Frontal Gyrus

We constructed the bilateral inferior frontal gyrus mask using the Harvard Oxford cortical atlas (from fsleyes version 1.6.1), based upon the Inferior Frontal Gyrus pars triangularis, which was additionally thresholded (p<.025) and resampled to 2mm voxels to match the dimensions of the BOLD images.

**Table 2.** Regions of Interest.

| Region of Interest | Source |
| --- | --- |
| Ventral Striatum | Bartra et al., 2013 |
| Dorsal Striatum | Bartra et al., 2013 |
| Anterior Insula | Bartra et al., 2013 |
| Dorsal ACC (Salience) | Schaefer et al., 2017; Kong et al., 2021; Ritz et al., 2024 |
| Dorsal ACC (Control) | Schaefer et al., 2017; Kong et al., 2021; Ritz et al., 2024 |
| Dorsolateral Prefrontal Cortex | Schaefer et al., 2017; Kong et al., 2021; Ritz et al., 2024 |
| Inferior Frontal Gyrus | Harvard-Oxford atlas (from FSL) |

### fMRI General Linear Models

#### Trialwise Estimates

Preprocessed data were submitted to linear mixed effects analyses using a two-step procedure. In the first step, we computed first level general linear models (GLM) in SPM to generate BOLD signal change estimates for each trial and participant. The GLM modeled a stick function at the onset of each cue and a response window at the start of each interval for the duration of that interval. The model also included nuisance regressors and modeled a stick function at the onset of each feedback and response window for error trials modulated by the duration of that trial (e.g., response time). Trials were concatenated across the eight task blocks and additional regressors were included to model within-block means and linear trends. GLMs were estimated using a reweighted least squares approach RobustWLS Toolbox^102^ to minimize the influence of outlier time-points (e.g., due to motion) and obtain optimal model estimates. We applied the ROI masks (ventral striatum, anterior insula, caudal and rostral dorsal anterior cingulate cortex, dorsolateral prefrontal cortex, and inferior frontal gyrus; see *ROI Selection* above) to the obtained estimates and calculated the mean activation for each ROI. These ROI estimates were analyzed using linear mixed effects models (model-free analyses), predicting neural activity, task performance, and subjective motivation and affect ratings. They were additionally included as neural regressors in model-based fMRI analyses (detailed below).

#### Whole-Brain Univariate Analyses

We complemented ROI analyses with whole-brain GLMs. For these analyses, we computed first-level GLMs, modeling the stick function at cues and intervals (as well as their interactions with reward and penalty incentives). Several of these GLMs included parametric regressors for reward and penalty incentives, RT and accuracy task performance, as well as for nuisance regressors such as interval number, interval length, and mean congruency. Regressors were de-orthogonalized to allow them to compete for variance^103^. As above, trials were concatenated across all eight task blocks, additional regressors were included to model within-block means and linear trends, and GLMs were estimated using RobustWLS. Second-level random effects analyses on first-level estimates were performed in SPM, and statistical inference was conducted using threshold-free cluster enhancement (TFCE) with familywise error correction across the whole brain (p<.05).^23^ Results were visualized using FSLeyes.

### Model-Based fMRI Analyses

Model fitting was performed in the same manner as the behavioral data (Supplemental Table S43), except now additionally including neural regressors to predict drift rate and threshold parameters. Trialwise beta estimates of cue-related and interval-related activity for each ROI were z-scored and included as neural regressors in our drift diffusion models to perform joint modeling of neural activity and task performance. Due to the high multicollinearity between ROIs, we separately included cue-related and interval-related activity in each model. To account for the temporal dynamics of the hemodynamic response function in fMRI BOLD signals, each trial contained the neural regressor from the relevant interval (e.g., all trials within the first interval were linked with cue- or interval-related estimates associated with that interval).

## Acknowledgments

This research was supported by NSF CAREER Award 2046111 and Carney Innovation Award to A.S., and a Brown OVPR Seed Award to A.S. and D.M.Y. DMY was supported by NIH Awards T32-MH12638, R25-NS124530, and K99-MH133912. X.L. was supported by NIH Training Fellowship T32-MH115895, and M.P.F. was supported by a NSF Graduate Research Fellowship. Advanced computing resources from the Brain Science Computer Cluster were supported by NIH Award S10-OD025181, and neuroimaging resources from the Behavior and Neurodata Core were supported by 5P30GM149405-02. We would like to thank Carolyn Dean Wolf for her assistance with design of the task cues, and members of the Shenhav Lab for invaluable discussions and feedback throughout the course of this project.

## Funding

National Science Foundation CAREER Award 2046111 (AS)

National Institutes of Health grant K99-MH133912 (DMY)

National Institutes of Health grant T32-MH12638 (DMY)

National Institutes of Health grant R25-NS124530 (DMY)

National Science Foundation Graduate Research Fellowship (MPF)

National Institutes of Health grant T32-MH115895 (XL)

National Institutes of Health grant S10-OD025181

National Institutes of Health grant 5P30GM149405-02

## Author contributions

Conceptualization: DMY, MPF, AS

Methodology: DMY, MPF, XL

Investigation: DMY, MPF, XL, ZC, JK, MT, KM, SN

Visualization: DMY

Supervision: DMY, AS

Writing—original draft: DMY

Writing—review & editing: DMY, MPF, XL, ZC, JK, MT, KM, SN

## Competing interests

Authors declare that they have no competing interests.

## Data and materials availability

All data, code, and materials are available on the OSF repository (https://doi.org/10.17605/OSF.IO/HJZ6V) and GitHub repository (https://github.com/debyee/TCB_fMRI)

## Notes

https://doi.org/10.17605/OSF.IO/HJZ6V

### Competing Interest Statement

The authors have declared no competing interest.

### Summary of Updates

This version of the manuscript has been revised to update the whole brain figures.

## References

1. Botvinick, M. & Braver, T. Motivation and Cognitive Control: From Behavior to Neural Mechanism. Annu. Rev. Psychol. 66, 83–113 (2015).

2. Yee, D. M. & Braver, T. S. Interactions of motivation and cognitive control. Curr. Opin. Behav. Sci. 19, 83–90 (2018).

3. Hu, H. Reward and Aversion. Annu. Rev. Neurosci. 39, 297–324 (2016).

4. Ben-Zion, Z. & Levy, I. Representation of Anticipated Rewards and Punishments in the Human Brain. Annu. Rev. Psychol. 76, 197–226 (2025).

5. Leng, X., Yee, D., Ritz, H. & Shenhav, A. Dissociable influences of reward and punishment on adaptive cognitive control. PLOS Comput. Biol. 17, e1009737 (2021).

6. Prater Fahey, M., Yee, D. M., Leng, X., Tarlow, M. & Shenhav, A. Motivational context determines the impact of aversive outcomes on mental effort allocation. Cognition 254, 105973 (2025).

7. Yee, D. M., Leng, X., Shenhav, A. & Braver, T. S. Aversive motivation and cognitive control. Neurosci. Biobehav. Rev. 133, 104493 (2022).

8. Manohar, S. G. et al. Reward Pays the Cost of Noise Reduction in Motor and Cognitive Control. Curr. Biol. 25, 1707–1716 (2015).

9. Simen, P. et al. Reward rate optimization in two-alternative decision making: Empirical tests of theoretical predictions. J. Exp. Psychol. Hum. Percept. Perform. 35, 1865–1897 (2009).

10. Bogacz, R., Brown, E., Moehlis, J., Holmes, P. & Cohen, J. D. The physics of optimal decision making: A formal analysis of models of performance in two-alternative forced-choice tasks. Psychol. Rev. 113, 700–765 (2006).

11. Musslick, S., Botvinick, M. M., Shenhav, A. & Cohen, J. D. A computational model of control allocation based on the Expected Value of Control. 6 (2015).

12. Ritz, H., Leng, X. & Shenhav, A. Cognitive Control as a Multivariate Optimization Problem. J. Cogn. Neurosci. 34, 569–591 (2022).

13. Parro, C., Dixon, M. L. & Christoff, K. The neural basis of motivational influences on cognitive control. Hum. Brain Mapp. 39, 5097–5111 (2018).

14. Lopez-Gamundi, P., et al. The neural basis of effort valuation: A meta-analysis of functional magnetic resonance imaging studies. bioRxiv 2021.01.08.425909 (2021) doi:10.1101/2021.01.08.425909.

15. Bartra, O., McGuire, J. T. & Kable, J. W. The valuation system: A coordinate-based meta-analysis of BOLD fMRI experiments examining neural correlates of subjective value. NeuroImage 76, 412–427 (2013).

16. Shenhav, A. et al. Toward a Rational and Mechanistic Account of Mental Effort. Annu. Rev. Neurosci. 40, 99–124 (2017).

17. Shenhav, A., Botvinick, M. M. & Cohen, J. D. The Expected Value of Control: An Integrative Theory of Anterior Cingulate Cortex Function. Neuron 79, 217–240 (2013).

18. Ratcliff, R., Smith, P. L., Brown, S. D. & McKoon, G. Diffusion Decision Model: Current Issues and History. Trends Cogn. Sci. 20, 260–281 (2016).

19. Wiecki, T. V., Sofer, I. & Frank, M. J. HDDM: Hierarchical Bayesian estimation of the Drift-Diffusion Model in Python. *Front*. Neuroinformatics 7, (2013).

20. Turner, B. M., Rodriguez, C. A., Norcia, T. M., McClure, S. M. & Steyvers, M. Why more is better: Simultaneous modeling of EEG, fMRI, and behavioral data. NeuroImage 128, 96–115 (2016).

21. Knutson, B. & Greer, S. M. Anticipatory affect: neural correlates and consequences for choice. Philos. Trans. R. Soc. B Biol. Sci. 363, 3771–3786 (2008).

22. Knutson, B., Katovich, K. & Suri, G. Inferring affect from fMRI data. Trends Cogn. Sci. 18, 422–428 (2014).

23. Smith, S. & Nichols, T. Threshold-free cluster enhancement: Addressing problems of smoothing, threshold dependence and localisation in cluster inference. NeuroImage 44, 83–98 (2009).

24. Frömer, R., Lin, H., Dean Wolf, C. K., Inzlicht, M. & Shenhav, A. Expectations of reward and efficacy guide cognitive control allocation. Nat. Commun. 12, 1030 (2021).

25. Krebs, R. M., Boehler, C. N. & Woldorff, M. G. The influence of reward associations on conflict processing in the Stroop task. Cognition 117, 341–347 (2010).

26. Braem, S., Duthoo, W. & Notebaert, W. Punishment Sensitivity Predicts the Impact of Punishment on Cognitive Control. PLoS ONE 8, e74106 (2013).

27. Yang, Q., Xing, J., Braem, S. & Pourtois, G. The selective use of punishments on congruent versus incongruent trials in the Stroop task. Neurobiol. Learn. Mem. 193, 107654 (2022).

28. Krebs, R. M., Boehler, C. N., Roberts, K. C., Song, A. W. & Woldorff, M. G. The Involvement of the Dopaminergic Midbrain and Cortico-Striatal-Thalamic Circuits in the Integration of Reward Prospect and Attentional Task Demands. Cereb. Cortex 22, 607–615 (2012).

29. Pessiglione, M. & Delgado, M. R. The good, the bad and the brain: neural correlates of appetitive and aversive values underlying decision making. Curr. Opin. Behav. Sci. 5, 78–84 (2015).

30. Spielberg, J. M. et al. A brain network instantiating approach and avoidance motivation. Psychophysiology 49, 1200–1214 (2012).

31. Palminteri, S. et al. Critical Roles for Anterior Insula and Dorsal Striatum in Punishment-Based Avoidance Learning. Neuron 76, 998–1009 (2012).

32. Kouneiher, F., Charron, S. & Koechlin, E. Motivation and cognitive control in the human prefrontal cortex. Nat. Neurosci. 12, 939–945 (2009).

33. Etzel, J. A., Cole, M. W., Zacks, J. M., Kay, K. N. & Braver, T. S. Reward Motivation Enhances Task Coding in Frontoparietal Cortex. Cereb. Cortex 26, 1647–1659 (2016).

34. Cubillo, A., Makwana, A. B. & Hare, T. A. Differential modulation of cognitive control networks by monetary reward and punishment. Soc. Cogn. Affect. Neurosci. 14, 305–317 (2019).

35. Yee, D. M., Crawford, J. L., Lamichhane, B. & Braver, T. S. Dorsal Anterior Cingulate Cortex Encodes the Integrated Incentive Motivational Value of Cognitive Task Performance. J. Neurosci. 41, 3707–3720 (2021).

36. Hampshire, A., Chamberlain, S. R., Monti, M. M., Duncan, J. & Owen, A. M. The role of the right inferior frontal gyrus: inhibition and attentional control. NeuroImage 50, 1313–1319 (2010).

37. Leng, X., Yee, D., Ritz, H. & Shenhav, A. Dissociable Influences of Reward and Punishment on Adaptive Cognitive Control. http://biorxiv.org/lookup/doi/10.1101/2020.09.11.294157 (2020) doi:10.1101/2020.09.11.294157.

38. Ullsperger, M., Danielmeier, C. & Jocham, G. Neurophysiology of Performance Monitoring and Adaptive Behavior. Physiol. Rev. 94, 35–79 (2014).

39. Chiew, K. S. Revisiting positive affect and reward influences on cognitive control. Curr. Opin. Behav. Sci. 39, 27–33 (2021).

40. Fröber, K. & Dreisbach, G. The differential influences of positive affect, random reward, and performance-contingent reward on cognitive control. Cogn. Affect. Behav. Neurosci. 14, 530–547 (2014).

41. Shackman, A. J. et al. The integration of negative affect, pain and cognitive control in the cingulate cortex. Nat. Rev. Neurosci. 12, 154–167 (2011).

42. Tolomeo, S. et al. A causal role for the anterior mid-cingulate cortex in negative affect and cognitive control. Brain 139, 1844–1854 (2016).

43. Braem, S. et al. The Role of Anterior Cingulate Cortex in the Affective Evaluation of Conflict. J. Cogn. Neurosci. 29, 137–149 (2017).

44. Engle, R. W. Role of Working-Memory Capacity in Cognitive Control. Curr. Anthropol. 51, S17–S26 (2010).

45. Braver, T. S. The variable nature of cognitive control: a dual mechanisms framework. Trends Cogn. Sci. 16, 106–113 (2012).

46. Musslick, S. & Cohen, J. D. Rationalizing Constraints on the Capacity for Cognitive Control. https://psyarxiv.com/vtknh/ (2020) doi:10.31234/osf.io/vtknh.

47. Miyake, A. & Friedman, N. P. The Nature and Organization of Individual Differences in Executive Functions: Four General Conclusions. Curr. Dir. Psychol. Sci. 21, 8–14 (2012).

48. Cheng, Z., Leng, X. & Shenhav, A. Incentive Effects Capture Variability in Task-General Control Allocation. Proc. 47th Annu. Meet. Cogn. Sci. Soc. (2025).

49. Schaefer, A. et al. Local-Global Parcellation of the Human Cerebral Cortex from Intrinsic Functional Connectivity MRI. Cereb. Cortex 28, 3095–3114 (2018).

50. Kong, R. et al. Individual-Specific Areal-Level Parcellations Improve Functional Connectivity Prediction of Behavior. Cereb. Cortex 31, 4477–4500 (2021).

51. Heilbronner, S. R. & Hayden, B. Y. Dorsal Anterior Cingulate Cortex: A Bottom-Up View. Annu. Rev. Neurosci. 39, 149–170 (2016).

52. Monosov, I. E., Haber, S. N., Leuthardt, E. C. & Jezzini, A. Anterior Cingulate Cortex and the Control of Dynamic Behavior in Primates. Curr. Biol. 30, R1442–R1454 (2020).

53. Vogt, B. A., Finch, D. M. & Olson, C. R. Functional Heterogeneity in Cingulate Cortex: The Anterior Executive and Posterior Evaluative Regions. Cereb. Cortex 2, 435–443 (1992).

54. Nee, D. E., Kastner, S. & Brown, J. W. Functional heterogeneity of conflict, error, task-switching, and unexpectedness effects within medial prefrontal cortex. NeuroImage 54, 528–540 (2011).

55. Kragel, P. A. et al. Generalizable representations of pain, cognitive control, and negative emotion in medial frontal cortex. Nat. Neurosci. 21, 283–289 (2018).

56. De La Vega, A., Chang, L. J., Banich, M. T., Wager, T. D. & Yarkoni, T. Large-Scale Meta-Analysis of Human Medial Frontal Cortex Reveals Tripartite Functional Organization. J. Neurosci. 36, 6553–6562 (2016).

57. Shenhav, A., Cohen, J. D. & Botvinick, M. M. Dorsal anterior cingulate cortex and the value of control. Nat. Neurosci. 19, 1286–1291 (2016).

58. Brockett, A. T. & Roesch, M. R. Anterior cingulate cortex and adaptive control of brain and behavior. in International Review of Neurobiology vol. 158 283–309 (Elsevier, 2021).

59. Shenhav, A., Straccia, M. A., Musslick, S., Cohen, J. D. & Botvinick, M. M. Dissociable neural mechanisms track evidence accumulation for selection of attention versus action. Nat. Commun. 9, 2485 (2018).

60. Venkatraman, V., Rosati, A. G., Taren, A. A. & Huettel, S. A. Resolving Response, Decision, and Strategic Control: Evidence for a Functional Topography in Dorsomedial Prefrontal Cortex. J. Neurosci. 29, 13158–13164 (2009).

61. Grahek, I., Frömer, R., Prater Fahey, M. & Shenhav, A. Learning when effort matters: neural dynamics underlying updating and adaptation to changes in performance efficacy. Cereb. Cortex 33, 2395–2411 (2023).

62. Ligneul, R., Mainen, Z. F., Ly, V. & Cools, R. Stress-sensitive inference of task controllability. *Nat*. Hum. Behav. 6, 812–822 (2022).

63. Woo, C.-W. et al. Separate neural representations for physical pain and social rejection. Nat. Commun. 5, 5380 (2014).

64. Speer, S. P. H. et al. A multivariate brain signature for reward. NeuroImage 271, 119990 (2023).

65. Coll, M.-P., et al. The neural signature of the decision value of future pain. Proc. Natl. Acad. Sci. 119, e2119931119 (2022).

66. Shenhav, A. The affective gradient hypothesis: An affect-centered account of motivated behavior. Preprint at 10.31234/osf.io/68yta (2024).

67. Eldar, E., Pessiglione, M. & van Dillen, L. Positive affect as a computational mechanism. Curr. Opin. Behav. Sci. 39, 52–57 (2021).

68. Tsakiris, M. & Critchley, H. Interoception beyond homeostasis: affect, cognition and mental health. Philos. Trans. R. Soc. B Biol. Sci. 371, 20160002 (2016).

69. Weber, L. A., Yee, D. M., Small, D. M. & Petzschner, F. H. The interoceptive origin of reinforcement learning. Trends Cogn. Sci. 29, 840–854 (2025).

70. Fonov, V., Evans, A., McKinstry, R., Almli, C. & Collins, D. Unbiased nonlinear average age-appropriate brain templates from birth to adulthood. NeuroImage 47, S102 (2009).

71. Huys, Q. J. M., Maia, T. V. & Frank, M. J. Computational psychiatry as a bridge from neuroscience to clinical applications. Nat. Neurosci. 19, 404–413 (2016).

72. Yip, S. W. et al. From Computation to Clinic. Biol. Psychiatry Glob. Open Sci. 3, 319–328 (2023).

73. Berwian, I. M., Hitchock, P., Pisupati, S., Schoen, G. & Niv, Y. Using computational models of learning to advance cognitive behavioral therapy. Commun. Psychol. 3, 72 (2025).

74. Monosov, I. E., Zimmermann, J., Frank, M. J., Mathis, M. W. & Baker, J. T. Ethological computational psychiatry: Challenges and opportunities. Curr. Opin. Neurobiol. 86, 102881 (2024).

75. Bates, D., Mächler, M., Bolker, B. & Walker, S. Fitting Linear Mixed-Effects Models Using **lme4**. J. Stat. Softw. 67, (2015).

76. Kuznetsova, A., Brockhoff, P. B. & Christensen, R. H. B. **lmerTest** Package: Tests in Linear Mixed Effects Models. J. Stat. Softw. 82, (2017).

77. Barr, D. J., Levy, R., Scheepers, C. & Tily, H. J. Random effects structure for confirmatory hypothesis testing: Keep it maximal. J. Mem. Lang. 68, 255–278 (2013).

78. Voeten, C. C. buildmer: Stepwise Elimination and Term Reordering for Mixed-Effects Regression. 2.12 10.32614/CRAN.package.buildmer (2019).

79. Nash, J. C. On Best Practice Optimization Methods in *R*. J. Stat. Softw. 60, (2014).

80. Nash, J. C. & Varadhan, R. Unifying Optimization Algorithms to Aid Software System Users: **optimx** for *R*. J. Stat. Softw. 43, (2011).

81. Ratcliff, R. & McKoon, G. The Diffusion Decision Model: Theory and Data for Two-Choice Decision Tasks. Neural Comput. 20, 873–922 (2008).

82. Fengler, A., Bera, K., Pedersen, M. L. & Frank, M. J. Beyond Drift Diffusion Models: Fitting a Broad Class of Decision and Reinforcement Learning Models with HDDM. J. Cogn. Neurosci. 34, 1780–1805 (2022).

83. Palestro, J. J., Weichart, E., Sederberg, P. B. & Turner, B. M. Some task demands induce collapsing bounds: Evidence from a behavioral analysis. Psychon. Bull. Rev. 25, 1225–1248 (2018).

84. Gelman, A. & Rubin, D. B. Inference from Iterative Simulation Using Multiple Sequences. Stat. Sci. 7, (1992).

85. Baribault, B. & Collins, A. G. E. Troubleshooting Bayesian cognitive models. Psychol. Methods 30, 128–154 (2025).

86. Wilson, R. C. & Collins, A. G. Ten simple rules for the computational modeling of behavioral data. eLife 8, e49547 (2019).

87. Esteban, O. et al. fMRIPrep: a robust preprocessing pipeline for functional MRI. Nat. Methods 16, 111–116 (2019).

88. Esteban, O., et al. nipy/nipype: 1.9.1. Zenodo 10.5281/ZENODO.596855 (2025).

89. Gorgolewski, K. et al. Nipype: A Flexible, Lightweight and Extensible Neuroimaging Data Processing Framework in Python. Front. Neuroinformatics 5, (2011).

90. Tustison, N. J. et al. N4ITK: Improved N3 Bias Correction. IEEE Trans. Med. Imaging 29, 1310–1320 (2010).

91. Avants, B., Epstein, C., Grossman, M. & Gee, J. Symmetric diffeomorphic image registration with cross-correlation: Evaluating automated labeling of elderly and neurodegenerative brain. Med. Image Anal. 12, 26–41 (2008).

92. Dale, A. M., Fischl, B. & Sereno, M. I. Cortical Surface-Based Analysis. NeuroImage 9, 179–194 (1999).

93. Klein, A. et al. Mindboggling morphometry of human brains. PLOS Comput. Biol. 13, e1005350 (2017).

94. Evans, A. C., Janke, A. L., Collins, D. L. & Baillet, S. Brain templates and atlases. NeuroImage 62, 911–922 (2012).

95. Glasser, M. F. et al. The minimal preprocessing pipelines for the Human Connectome Project. NeuroImage 80, 105–124 (2013).

96. Greve, D. N. & Fischl, B. Accurate and robust brain image alignment using boundary-based registration. NeuroImage 48, 63–72 (2009).

97. Jenkinson, M., Bannister, P., Brady, M. & Smith, S. Improved Optimization for the Robust and Accurate Linear Registration and Motion Correction of Brain Images. NeuroImage 17, 825–841 (2002).

98. Cox, R. W. & Hyde, J. S. Software tools for analysis and visualization of fMRI data. NMR Biomed. 10, 171–178 (1997).

99. Pruim, R. H. R. et al. ICA-AROMA: A robust ICA-based strategy for removing motion artifacts from fMRI data. NeuroImage 112, 267–277 (2015).

100. Ritz, H. & Shenhav, A. Orthogonal neural encoding of targets and distractors supports multivariate cognitive control. *Nat*. Hum. Behav. 8, 945–961 (2024).

101. Yarkoni, T., Poldrack, R. A., Nichols, T. E., Van Essen, D. C. & Wager, T. D. Large-scale automated synthesis of human functional neuroimaging data. Nat. Methods 8, 665–670 (2011).

102. Diedrichsen, J. & Shadmehr, R. Detecting and adjusting for artifacts in fMRI time series data. NeuroImage 27, 624–634 (2005).

103. Mumford, J. A., Poline, J.-B. & Poldrack, R. A. Orthogonalization of Regressors in fMRI Models. PLOS ONE 10, e0126255 (2015).

