## Supplemental Material for "Neurocomputational mechanisms underlying the distinct motivational influences of reward and punishment on cognitive control"

**Table S1:** *Ventral Striatum Activity During the Cue and Interval Phase*. Linear mixed models of ROI during cue and interval task phases by reward and penalty incentive conditions, during all conditions, reward-varying conditions, and penalty-varying conditions.

**Ventral Striatum Cue**

| | ROI - All Trials | | | | ROI - Vary Reward | | | | ROI - Vary Penalty | | | |
|---|---|---|---|---|---|---|---|---|---|---|---|---|
| *Predictors* | *Estimates* | *CI* | *Statistic* | *p* | *Estimates* | *CI* | *Statistic* | *p* | *Estimates* | *CI* | *Statistic* | *p* |
| (Intercept) | 0.06 * | 0.00 – 0.11 | 2.12 | **0.034** | 0.06 | -0.01 – 0.12 | 1.72 | 0.085 | 0.07 * | 0.01 – 0.13 | 2.17 | **0.030** |
| Reward | 0.02 | -0.01 – 0.04 | 1.46 | 0.145 | 0.05 *** | 0.02 – 0.08 | 3.46 | **0.001** | -0.02 | -0.07 – 0.03 | -0.91 | 0.363 |
| Penalty | 0.05 ** | 0.02 – 0.08 | 3.02 | **0.003** | 0.06 ** | 0.02 – 0.10 | 2.63 | **0.009** | 0.01 | -0.03 – 0.06 | 0.57 | 0.571 |
| Interval Congruency | -0.00 | -0.02 – 0.02 | -0.18 | 0.859 | -0.00 | -0.03 – 0.02 | -0.32 | 0.751 | 0.00 | -0.03 – 0.03 | 0.15 | 0.878 |
| Interval Length | -0.01 | -0.04 – 0.01 | -1.18 | 0.239 | -0.01 | -0.04 – 0.02 | -0.77 | 0.440 | 0.01 | -0.03 – 0.05 | 0.51 | 0.611 |
| Interval Session Number | 0.01 | -0.01 – 0.03 | 1.21 | 0.226 | 0.03 | -0.00 – 0.06 | 1.87 | 0.062 | -0.02 | -0.06 – 0.01 | -1.52 | 0.129 |
| Reward:Penalty | -0.01 | -0.04 – 0.01 | -1.39 | 0.164 | -0.02 | -0.04 – 0.01 | -1.12 | 0.261 | -0.01 | -0.04 – 0.02 | -0.91 | 0.362 |
| **Random Effects** | | | | | | | | | | | | |
| $\sigma^2$ | 1.30 | | | | 1.27 | | | | 1.23 | | | |
| $\tau_{00}$ | 0.05 $_{\text{SubID}}$ | | | | 0.08 $_{\text{SubID}}$ | | | | 0.07 $_{\text{SubID}}$ | | | |
| $\tau_{11}$ | 0.01 $_{\text{SubID.catPenaltyLevel1}}$ | | | | 0.03 $_{\text{SubID.catPenaltyLevel1}}$ | | | | 0.04 $_{\text{SubID.catRewardLevel1}}$ | | | |
| $\rho_{01}$ | -0.10 $_{\text{SubID}}$ | | | | 0.13 $_{\text{SubID}}$ | | | | -0.09 $_{\text{SubID}}$ | | | |
| N | 94 $_{\text{SubID}}$ | | | | 94 $_{\text{SubID}}$ | | | | 94 $_{\text{SubID}}$ | | | |
| Observations | 11551 | | | | 5998 | | | | 5553 | | | |
| Marginal $R^2$ / Conditional $R^2$ | 0.002 / 0.049 | | | | 0.005 / 0.082 | | | | 0.001 / 0.080 | | | |

** p<0.05 ** p<0.01 *** p<0.001*

**Ventral Striatum Interval**

| | ROI - All Trials | | | | ROI - Vary Reward | | | | ROI - Vary Penalty | | | |
|---|---|---|---|---|---|---|---|---|---|---|---|---|
| *Predictors* | *Estimates* | *CI* | *Statistic* | *p* | *Estimates* | *CI* | *Statistic* | *p* | *Estimates* | *CI* | *Statistic* | *p* |
| (Intercept) | 0.19 *** | 0.17 – 0.20 | 21.39 | **<0.001** | 0.18 *** | 0.17 – 0.20 | 18.40 | **<0.001** | 0.07 * | 0.01 – 0.13 | 2.17 | **0.030** |
| Reward | 0.00 | -0.00 – 0.01 | 1.00 | 0.317 | 0.01 * | 0.00 – 0.01 | 2.06 | **0.040** | -0.02 | -0.07 – 0.03 | -0.91 | 0.363 |
| Penalty | 0.00 | -0.00 – 0.01 | 1.28 | 0.201 | 0.00 | -0.00 – 0.01 | 1.00 | 0.318 | 0.01 | -0.03 – 0.06 | 0.57 | 0.571 |
| Interval Congruency | -0.00 | -0.00 – 0.00 | -0.46 | 0.642 | -0.00 | -0.01 – 0.00 | -1.54 | 0.123 | 0.00 | -0.03 – 0.03 | 0.15 | 0.878 |
| Interval Length | -0.00 | -0.01 – 0.00 | -0.77 | 0.441 | -0.00 | -0.01 – 0.00 | -0.70 | 0.482 | 0.01 | -0.03 – 0.05 | 0.51 | 0.611 |
| Interval Session Number | 0.01 ** | 0.00 – 0.01 | 2.70 | **0.007** | 0.01 *** | 0.01 – 0.02 | 4.09 | **<0.001** | -0.02 | -0.06 – 0.01 | -1.52 | 0.129 |
| Reward:Penalty | 0.00 | -0.00 – 0.01 | 0.72 | 0.472 | 0.00 | -0.00 – 0.01 | 1.77 | 0.077 | -0.01 | -0.04 – 0.02 | -0.91 | 0.362 |
| **Random Effects** | | | | | | | | | | | | |
| $\sigma^2$ | 0.04 | | | | 0.04 | | | | 1.23 | | | |
| $\tau_{00}$ | 0.01 $_{\text{SubID}}$ | | | | 0.01 $_{\text{SubID}}$ | | | | 0.07 $_{\text{SubID}}$ | | | |
| $\tau_{11}$ | 0.00 $_{\text{SubID.catPenaltyLevel1}}$ | | | | 0.00 $_{\text{SubID.catPenaltyLevel1}}$ | | | | 0.04 $_{\text{SubID.catRewardLevel1}}$ | | | |
| | 0.00 $_{\text{SubID.catRewardLevel1}}$ | | | | 0.00 $_{\text{SubID.catRewardLevel1}}$ | | | | | | | |
| $\rho_{01}$ | -0.24 | | | | 0.01 | | | | -0.09 $_{\text{SubID}}$ | | | |
| | 0.25 | | | | 0.58 | | | | | | | |
| N | 94 $_{\text{SubID}}$ | | | | 94 $_{\text{SubID}}$ | | | | 94 $_{\text{SubID}}$ | | | |
| Observations | 11551 | | | | 5998 | | | | 5553 | | | |
| Marginal $R^2$ / Conditional $R^2$ | 0.001 / 0.162 | | | | 0.005 / 0.216 | | | | 0.001 / 0.080 | | | |

** p<0.05 ** p<0.01 *** p<0.001*

**Table S2:** *Anterior Insula Activity During the Cue and Interval Phase*. Linear mixed models of ROI during cue and interval task phases by reward and penalty incentive conditions, during all conditions, reward-varying conditions, and penalty-varying conditions.

**AI Cue**

| | ROI - All Trials | | | | ROI - Vary Reward | | | | ROI - Vary Penalty | | | |
|---|---|---|---|---|---|---|---|---|---|---|---|---|
| *Predictors* | *Estimates* | *CI* | *Statistic* | *p* | *Estimates* | *CI* | *Statistic* | *p* | *Estimates* | *CI* | *Statistic* | *p* |
| (Intercept) | -0.14 *** | -0.19 – -0.09 | -5.05 | **<0.001** | -0.14 *** | -0.21 – -0.07 | -3.87 | **<0.001** | -0.13 *** | -0.19 – -0.08 | -4.60 | **<0.001** |
| Reward | -0.02 | -0.04 – 0.01 | -1.51 | 0.130 | -0.01 | -0.04 – 0.02 | -0.65 | 0.514 | -0.03 | -0.07 – 0.02 | -1.08 | 0.282 |
| Penalty | 0.03 * | 0.00 – 0.06 | 2.03 | **0.042** | 0.04 | -0.01 – 0.08 | 1.64 | 0.101 | 0.01 | -0.04 – 0.06 | 0.44 | 0.657 |
| Interval Congruency | 0.00 | -0.02 – 0.02 | 0.18 | 0.856 | -0.00 | -0.03 – 0.03 | -0.05 | 0.958 | 0.01 | -0.03 – 0.04 | 0.39 | 0.698 |
| Interval Length | -0.00 | -0.02 – 0.02 | -0.05 | 0.960 | 0.00 | -0.03 – 0.03 | 0.12 | 0.905 | 0.01 | -0.04 – 0.06 | 0.53 | 0.596 |
| Interval Session Number | -0.04 *** | -0.07 – -0.02 | -3.68 | **<0.001** | -0.05 ** | -0.08 – -0.02 | -2.87 | **0.004** | -0.04 * | -0.07 – -0.00 | -2.03 | **0.043** |
| Reward:Penalty | -0.03 * | -0.05 – -0.01 | -2.51 | **0.012** | -0.04 ** | -0.07 – -0.01 | -2.63 | **0.009** | -0.02 | -0.05 – 0.02 | -0.95 | 0.343 |
| **Random Effects** | | | | | | | | | | | | |
| $\sigma^2$ | 1.55 | | | | 1.49 | | | | 1.53 | | | |
| $\tau_{00}$ | 0.06 SubID | | | | 0.10 SubID | | | | 0.05 SubID | | | |
| $\tau_{11}$ | 0.01 SubID.catPenaltyLevel1 | | | | 0.03 SubID.catPenaltyLevel1 | | | | 0.03 SubID.catRewardLevel1 | | | |
| $\rho_{01}$ | -0.01 SubID | | | | 0.26 SubID | | | | 0.11 SubID | | | |
| N | 94 SubID | | | | 94 SubID | | | | 94 SubID | | | |
| Observations | 11551 | | | | 5998 | | | | 5553 | | | |
| Marginal $R^2$ / Conditional $R^2$ | 0.002 / 0.043 | | | | 0.004 / 0.081 | | | | 0.002 / 0.052 | | | |

** p<0.05   ** p<0.01   *** p<0.001*

**AI Interval**

| | ROI - All Trials | | | | ROI - Vary Reward | | | | ROI - Vary Penalty | | | |
|---|---|---|---|---|---|---|---|---|---|---|---|---|
| *Predictors* | *Estimates* | *CI* | *Statistic* | *p* | *Estimates* | *CI* | *Statistic* | *p* | *Estimates* | *CI* | *Statistic* | *p* |
| (Intercept) | -0.13 *** | -0.14 – -0.11 | -15.25 | **<0.001** | -0.12 *** | -0.14 – -0.11 | -13.13 | **<0.001** | -0.13 *** | -0.14 – -0.11 | -14.46 | **<0.001** |
| Reward | -0.00 | -0.01 – 0.00 | -0.89 | 0.375 | 0.00 | -0.01 – 0.01 | 0.17 | 0.869 | -0.01 | -0.01 – 0.00 | -1.20 | 0.231 |
| Penalty | 0.01 *** | 0.00 – 0.02 | 3.46 | **0.001** | 0.01 ** | 0.00 – 0.02 | 2.65 | **0.008** | 0.01 | -0.00 – 0.01 | 1.18 | 0.238 |
| Interval Congruency | 0.00 | -0.00 – 0.01 | 1.31 | 0.191 | -0.00 | -0.01 – 0.01 | -0.06 | 0.955 | 0.01 * | 0.00 – 0.01 | 2.16 | **0.031** |
| Interval Length | -0.00 | -0.01 – 0.00 | -1.20 | 0.232 | -0.00 | -0.01 – 0.00 | -0.97 | 0.332 | 0.00 | -0.01 – 0.01 | 0.28 | 0.782 |
| Interval Session Number | -0.02 *** | -0.02 – -0.02 | -10.01 | **<0.001** | -0.02 *** | -0.02 – -0.01 | -5.71 | **<0.001** | -0.02 *** | -0.03 – -0.02 | -7.10 | **<0.001** |
| Reward:Penalty | -0.00 * | -0.01 – -0.00 | -2.05 | **0.041** | -0.00 | -0.01 – 0.00 | -1.49 | 0.137 | -0.00 | -0.01 – 0.00 | -1.49 | 0.136 |
| **Random Effects** | | | | | | | | | | | | |
| $\sigma^2$ | 0.05 | | | | 0.05 | | | | 0.05 | | | |
| $\tau_{00}$ | 0.01 SubID | | | | 0.01 SubID | | | | 0.01 SubID | | | |
| $\tau_{11}$ | 0.00 SubID.catPenaltyLevel1 | | | | 0.00 SubID.catPenaltyLevel1 | | | | 0.00 SubID.catRewardLevel1 | | | |
| | 0.00 SubID.catRewardLevel1 | | | | | | | | 0.00 SubID.catPenaltyLevel1 | | | |
| $\rho_{01}$ | 0.12 | | | | 0.18 SubID | | | | -0.20 | | | |
| | -0.01 | | | | | | | | -0.27 | | | |
| N | 94 SubID | | | | 94 SubID | | | | 94 SubID | | | |
| Observations | 11551 | | | | 5998 | | | | 5553 | | | |
| Marginal $R^2$ / Conditional $R^2$ | 0.010 / 0.127 | | | | 0.009 / 0.161 | | | | 0.012 / 0.148 | | | |

** p<0.05   ** p<0.01   *** p<0.001*

**Table S3:** *Caudal dACC Activity During the Cue and Interval Phase*. Linear mixed models of ROI during cue and interval task phases by reward and penalty incentive conditions, during all conditions, reward-varying conditions, and penalty-varying conditions.

**Caudal dACC Cue**

| | ROI - All Trials | | | | ROI - Vary Reward | | | | ROI - Vary Penalty | | | |
|---|---|---|---|---|---|---|---|---|---|---|---|---|
| *Predictors* | *Estimates* | *CI* | *Statistic* | *p* | *Estimates* | *CI* | *Statistic* | *p* | *Estimates* | *CI* | *Statistic* | *p* |
| (Intercept) | 0.07 * | 0.02 – 0.12 | 2.54 | **0.011** | 0.06 | -0.01 – 0.13 | 1.69 | 0.091 | 0.08 ** | 0.02 – 0.15 | 2.72 | **0.007** |
| Reward | 0.02 | -0.01 – 0.04 | 1.11 | 0.269 | 0.04 ** | 0.01 – 0.07 | 2.63 | **0.009** | -0.01 | -0.06 – 0.03 | -0.53 | 0.599 |
| Penalty | 0.03 | -0.00 – 0.06 | 1.69 | 0.091 | 0.03 | -0.01 – 0.08 | 1.44 | 0.151 | 0.01 | -0.04 – 0.06 | 0.40 | 0.692 |
| Interval Congruency | 0.01 | -0.02 – 0.03 | 0.55 | 0.582 | 0.00 | -0.03 – 0.03 | 0.16 | 0.869 | 0.01 | -0.02 – 0.04 | 0.51 | 0.611 |
| Interval Length | 0.01 | -0.02 – 0.03 | 0.70 | 0.485 | 0.01 | -0.02 – 0.04 | 0.60 | 0.546 | 0.02 | -0.02 – 0.07 | 0.95 | 0.340 |
| Interval Session Number | -0.00 | -0.02 – 0.02 | -0.19 | 0.852 | -0.02 | -0.05 – 0.02 | -1.03 | 0.303 | 0.01 | -0.03 – 0.04 | 0.40 | 0.690 |
| Reward:Penalty | -0.03 ** | -0.05 – -0.01 | -2.86 | **0.004** | -0.05 *** | -0.08 – -0.02 | -3.39 | **0.001** | -0.01 | -0.04 – 0.02 | -0.57 | 0.571 |
| **Random Effects** | | | | | | | | | | | | |
| $\sigma^2$ | 1.44 | | | | 1.40 | | | | 1.40 | | | |
| $\tau_{00}$ | 0.06 SubID | | | | 0.10 SubID | | | | 0.07 SubID | | | |
| $\tau_{11}$ | 0.01 SubID.catPenaltyLevel1 | | | | 0.03 SubID.catPenaltyLevel1 | | | | 0.03 SubID.catRewardLevel1 | | | |
| | 0.01 SubID.catRewardLevel1 | | | | | | | | | | | |
| $\rho_{01}$ | -0.05 | | | | 0.10 SubID | | | | -0.31 SubID | | | |
| | -0.15 | | | | | | | | | | | |
| N | 94 SubID | | | | 94 SubID | | | | 94 SubID | | | |
| Observations | 11551 | | | | 5998 | | | | 5553 | | | |
| Marginal $R^2$ / Conditional $R^2$ | 0.002 / 0.052 | | | | 0.004 / 0.090 | | | | 0.001 / 0.066 | | | |

** p<0.05 ** p<0.01 *** p<0.001*

**Caudal dACC Interval**

| | ROI - All Trials | | | | ROI - Vary Reward | | | | ROI - Vary Penalty | | | |
|---|---|---|---|---|---|---|---|---|---|---|---|---|
| *Predictors* | *Estimates* | *CI* | *Statistic* | *p* | *Estimates* | *CI* | *Statistic* | *p* | *Estimates* | *CI* | *Statistic* | *p* |
| (Intercept) | 0.24 *** | 0.22 – 0.25 | 25.42 | **<0.001** | 0.23 *** | 0.21 – 0.25 | 22.95 | **<0.001** | 0.24 *** | 0.22 – 0.26 | 24.47 | **<0.001** |
| Reward | 0.00 | -0.00 – 0.01 | 1.50 | 0.133 | 0.01 ** | 0.00 – 0.01 | 2.66 | **0.008** | 0.00 | -0.01 – 0.01 | 0.17 | 0.865 |
| Penalty | 0.00 | -0.00 – 0.01 | 0.82 | 0.412 | 0.00 | -0.01 – 0.01 | 0.53 | 0.599 | 0.00 | -0.01 – 0.01 | 0.30 | 0.764 |
| Interval Congruency | -0.00 | -0.00 – 0.00 | -0.48 | 0.630 | -0.00 | -0.01 – 0.00 | -1.48 | 0.140 | 0.00 | -0.00 – 0.01 | 0.96 | 0.335 |
| Interval Length | -0.01 ** | -0.01 – -0.00 | -3.27 | **0.001** | -0.01 ** | -0.01 – -0.00 | -2.61 | **0.009** | -0.01 | -0.01 – 0.00 | -1.33 | 0.184 |
| Interval Session Number | -0.02 *** | -0.02 – -0.01 | -8.78 | **<0.001** | -0.02 *** | -0.02 – -0.01 | -6.00 | **<0.001** | -0.02 *** | -0.03 – -0.01 | -6.47 | **<0.001** |
| Reward:Penalty | -0.00 | -0.01 – 0.00 | -1.01 | 0.312 | -0.00 | -0.01 – 0.00 | -0.29 | 0.775 | -0.00 | -0.01 – 0.00 | -1.15 | 0.250 |
| **Random Effects** | | | | | | | | | | | | |
| $\sigma^2$ | 0.04 | | | | 0.04 | | | | 0.04 | | | |
| $\tau_{00}$ | 0.01 SubID | | | | 0.01 SubID | | | | 0.01 SubID | | | |
| $\tau_{11}$ | 0.00 SubID.catRewardLevel1 | | | | 0.00 SubID.catPenaltyLevel1 | | | | 0.00 SubID.catRewardLevel1 | | | |
| | 0.00 SubID.catPenaltyLevel1 | | | | 0.00 SubID.catRewardLevel1 | | | | 0.00 SubID.catPenaltyLevel1 | | | |
| $\rho_{01}$ | 0.15 | | | | -0.06 | | | | -0.00 | | | |
| | -0.15 | | | | 0.37 | | | | -0.32 | | | |
| N | 94 SubID | | | | 94 SubID | | | | 94 SubID | | | |
| Observations | 11551 | | | | 5998 | | | | 5553 | | | |
| Marginal $R^2$ / Conditional $R^2$ | 0.007 / 0.175 | | | | 0.009 / 0.208 | | | | 0.008 / 0.214 | | | |

** p<0.05 ** p<0.01 *** p<0.001*

**Table S4:** *Rostral dACC Activity During the Cue and Interval Phase*. Linear mixed models of ROI during cue and interval task phases by reward and penalty incentive conditions, during all conditions, reward-varying conditions, and penalty-varying conditions.

**Rostral dACC Cue**

| | ROI - All Trials | | | | ROI - Vary Reward | | | | ROI - Vary Penalty | | | |
|---|---|---|---|---|---|---|---|---|---|---|---|---|
| *Predictors* | *Estimates* | *CI* | *Statistic* | *p* | *Estimates* | *CI* | *Statistic* | *p* | *Estimates* | *CI* | *Statistic* | *p* |
| (Intercept) | -0.07 ** | -0.12 – -0.02 | -2.63 | **0.009** | -0.07 | -0.14 – 0.00 | -1.94 | 0.052 | -0.07 * | -0.12 – -0.01 | -2.24 | **0.025** |
| Reward | -0.01 | -0.04 – 0.02 | -0.83 | 0.404 | -0.01 | -0.04 – 0.02 | -0.75 | 0.451 | -0.01 | -0.07 – 0.04 | -0.53 | 0.595 |
| Penalty | 0.03 * | 0.00 – 0.06 | 2.14 | **0.032** | 0.04 | -0.00 – 0.08 | 1.79 | 0.074 | 0.03 | -0.01 – 0.08 | 1.39 | 0.165 |
| Interval Congruency | 0.01 | -0.01 – 0.03 | 0.77 | 0.440 | -0.00 | -0.03 – 0.03 | -0.03 | 0.979 | 0.02 | -0.01 – 0.05 | 1.27 | 0.204 |
| Interval Length | 0.00 | -0.02 – 0.02 | 0.06 | 0.950 | 0.01 | -0.02 – 0.05 | 0.89 | 0.375 | -0.01 | -0.06 – 0.03 | -0.53 | 0.593 |
| Interval Session Number | -0.06 *** | -0.08 – -0.03 | -4.88 | **<0.001** | -0.09 *** | -0.12 – -0.05 | -5.13 | **<0.001** | -0.04 * | -0.07 – -0.00 | -2.00 | **0.045** |
| Reward:Penalty | -0.01 | -0.04 – 0.01 | -1.25 | 0.211 | -0.02 | -0.05 – 0.01 | -1.47 | 0.142 | -0.00 | -0.04 – 0.03 | -0.26 | 0.797 |
| **Random Effects** | | | | | | | | | | | | |
| $\sigma^2$ | 1.53 | | | | 1.49 | | | | 1.49 | | | |
| $\tau_{00}$ | 0.05 $_{SubID}$ | | | | 0.10 $_{SubID}$ | | | | 0.05 $_{SubID}$ | | | |
| $\tau_{11}$ | 0.01 $_{SubID.catRewardLevel1}$ | | | | 0.02 $_{SubID.catPenaltyLevel1}$ | | | | 0.05 $_{SubID.catRewardLevel1}$ | | | |
| | 0.01 $_{SubID.catPenaltyLevel1}$ | | | | | | | | | | | |
| $\rho_{01}$ | -0.08 | | | | 0.19 $_{SubID}$ | | | | -0.13 $_{SubID}$ | | | |
| | 0.07 | | | | | | | | | | | |
| N | 94 $_{SubID}$ | | | | 94 $_{SubID}$ | | | | 94 $_{SubID}$ | | | |
| Observations | 11551 | | | | 5998 | | | | 5553 | | | |
| Marginal $R^2$ / Conditional $R^2$ | 0.003 / 0.046 | | | | 0.006 / 0.078 | | | | 0.002 / 0.066 | | | |

** p<0.05 ** p<0.01 *** p<0.001*

**Rostral dACC Interval**

| | ROI - All Trials | | | | ROI - Vary Reward | | | | ROI - Vary Penalty | | | |
|---|---|---|---|---|---|---|---|---|---|---|---|---|
| *Predictors* | *Estimates* | *CI* | *Statistic* | *p* | *Estimates* | *CI* | *Statistic* | *p* | *Estimates* | *CI* | *Statistic* | *p* |
| (Intercept) | -0.14 *** | -0.15 – -0.12 | -14.74 | **<0.001** | -0.13 *** | -0.15 – -0.11 | -12.96 | **<0.001** | -0.14 *** | -0.15 – -0.12 | -14.85 | **<0.001** |
| Reward | -0.01 ** | -0.01 – -0.00 | -2.99 | **0.003** | -0.01 * | -0.01 – -0.00 | -2.45 | **0.014** | -0.01 | -0.02 – 0.00 | -1.92 | 0.054 |
| Penalty | 0.01 * | 0.00 – 0.01 | 2.14 | **0.032** | 0.01 | -0.00 – 0.02 | 1.47 | 0.142 | 0.01 | -0.00 – 0.02 | 1.35 | 0.178 |
| Interval Congruency | 0.00 | -0.00 – 0.01 | 0.54 | 0.586 | -0.00 | -0.01 – 0.00 | -1.02 | 0.309 | 0.01 * | 0.00 – 0.01 | 2.03 | **0.043** |
| Interval Length | 0.00 | -0.00 – 0.01 | 0.48 | 0.629 | 0.00 | -0.00 – 0.01 | 0.55 | 0.583 | 0.00 | -0.01 – 0.01 | 0.19 | 0.848 |
| Interval Session Number | -0.01 ** | -0.01 – -0.00 | -2.98 | **0.003** | -0.01 * | -0.01 – -0.00 | -1.97 | **0.048** | -0.01 ** | -0.02 – -0.00 | -2.95 | **0.003** |
| Reward:Penalty | -0.00 | -0.01 – 0.00 | -1.74 | 0.081 | -0.00 | -0.01 – 0.00 | -1.29 | 0.197 | -0.00 | -0.01 – 0.00 | -1.26 | 0.206 |
| **Random Effects** | | | | | | | | | | | | |
| $\sigma^2$ | 0.05 | | | | 0.05 | | | | 0.05 | | | |
| $\tau_{00}$ | 0.01 $_{SubID}$ | | | | 0.01 $_{SubID}$ | | | | 0.01 $_{SubID}$ | | | |
| $\tau_{11}$ | 0.00 $_{SubID.catRewardLevel1}$ | | | | 0.00 $_{SubID.catPenaltyLevel1}$ | | | | 0.00 $_{SubID.catRewardLevel1}$ | | | |
| | 0.00 $_{SubID.catPenaltyLevel1}$ | | | | 0.00 $_{SubID.catRewardLevel1}$ | | | | 0.00 $_{SubID.catPenaltyLevel1}$ | | | |
| $\rho_{01}$ | 0.18 | | | | 0.04 | | | | -0.12 | | | |
| | -0.10 | | | | 0.69 | | | | -0.31 | | | |
| N | 94 $_{SubID}$ | | | | 94 $_{SubID}$ | | | | 94 $_{SubID}$ | | | |
| Observations | 11551 | | | | 5998 | | | | 5553 | | | |
| Marginal $R^2$ / Conditional $R^2$ | 0.003 / 0.145 | | | | 0.003 / 0.182 | | | | 0.005 / 0.156 | | | |

** p<0.05 ** p<0.01 *** p<0.001*

**Table S5:** *Dorsolateral PFC Activity During the Cue and Interval Phase*. Linear mixed models of ROI during cue and interval task phases by reward and penalty incentive conditions, during all conditions, reward-varying conditions, and penalty-varying conditions.

**Dorsolateral PFC Cue**

| | ROI - All Trials | | | | ROI - Vary Reward | | | | ROI - Vary Penalty | | | |
|---|---|---|---|---|---|---|---|---|---|---|---|---|
| *Predictors* | *Estimates* | *CI* | *Statistic* | *p* | *Estimates* | *CI* | *Statistic* | *p* | *Estimates* | *CI* | *Statistic* | *p* |
| (Intercept) | -0.02 | -0.08 – 0.04 | -0.76 | 0.446 | -0.02 | -0.10 – 0.06 | -0.42 | 0.677 | -0.02 | -0.09 – 0.05 | -0.61 | 0.543 |
| Reward | -0.00 | -0.03 – 0.02 | -0.42 | 0.677 | 0.02 | -0.01 – 0.05 | 1.26 | 0.207 | -0.04 | -0.09 – 0.02 | -1.27 | 0.206 |
| Penalty | 0.04 ** | 0.01 – 0.08 | 2.61 | **0.009** | 0.06 * | 0.01 – 0.11 | 2.31 | **0.021** | 0.02 | -0.03 – 0.07 | 0.81 | 0.417 |
| Interval Congruency | 0.00 | -0.02 – 0.03 | 0.41 | 0.680 | 0.01 | -0.02 – 0.04 | 0.48 | 0.628 | 0.00 | -0.03 – 0.03 | 0.07 | 0.940 |
| Interval Length | 0.00 | -0.02 – 0.02 | 0.01 | 0.994 | 0.01 | -0.02 – 0.04 | 0.78 | 0.434 | 0.01 | -0.04 – 0.06 | 0.35 | 0.726 |
| Interval Session Number | -0.04 *** | -0.06 – -0.02 | -3.52 | **<0.001** | -0.06 ** | -0.09 – -0.02 | -3.16 | **0.002** | -0.03 | -0.07 – 0.00 | -1.81 | 0.070 |
| Reward:Penalty | -0.03 ** | -0.06 – -0.01 | -2.79 | **0.005** | -0.05 *** | -0.08 – -0.02 | -3.32 | **0.001** | -0.01 | -0.04 – 0.02 | -0.54 | 0.591 |
| **Random Effects** | | | | | | | | | | | | |
| $\sigma^2$ | 1.56 | | | | 1.55 | | | | 1.44 | | | |
| $\tau_{00}$ | 0.08 $_{SubID}$ | | | | 0.13 $_{SubID}$ | | | | 0.09 $_{SubID}$ | | | |
| $\tau_{11}$ | 0.01 $_{SubID.catPenaltyLevel1}$ | | | | 0.04 $_{SubID.catPenaltyLevel1}$ | | | | 0.05 $_{SubID.catRewardLevel1}$ | | | |
| $\rho_{01}$ | -0.07 $_{SubID}$ | | | | 0.11 $_{SubID}$ | | | | -0.18 $_{SubID}$ | | | |
| N | 94 $_{SubID}$ | | | | 94 $_{SubID}$ | | | | 94 $_{SubID}$ | | | |
| Observations | 11551 | | | | 5998 | | | | 5553 | | | |
| Marginal $R^2$ / Conditional $R^2$ | 0.003 / 0.057 | | | | 0.006 / 0.104 | | | | 0.002 / 0.090 | | | |

** p<0.05 ** p<0.01 *** p<0.001*

**Dorsolateral PFC Interval**

| | ROI - All Trials | | | | ROI - Vary Reward | | | | ROI - Vary Penalty | | | |
|---|---|---|---|---|---|---|---|---|---|---|---|---|
| *Predictors* | *Estimates* | *CI* | *Statistic* | *p* | *Estimates* | *CI* | *Statistic* | *p* | *Estimates* | *CI* | *Statistic* | *p* |
| (Intercept) | -0.04 *** | -0.06 – -0.03 | -4.90 | **<0.001** | -0.04 *** | -0.06 – -0.02 | -4.48 | **<0.001** | -0.04 *** | -0.06 – -0.02 | -4.22 | **<0.001** |
| Reward | -0.00 | -0.01 – 0.00 | -1.18 | 0.239 | 0.00 | -0.00 – 0.01 | 0.58 | 0.563 | -0.01 | -0.02 – 0.00 | -1.43 | 0.153 |
| Penalty | 0.01 * | 0.00 – 0.01 | 2.27 | **0.023** | 0.01 ** | 0.00 – 0.01 | 2.98 | **0.003** | 0.00 | -0.01 – 0.01 | 0.72 | 0.469 |
| Interval Congruency | -0.00 | -0.00 – 0.00 | -0.23 | 0.815 | -0.00 | -0.01 – 0.00 | -0.85 | 0.393 | 0.00 | -0.00 – 0.01 | 1.06 | 0.289 |
| Interval Length | -0.00 | -0.00 – 0.00 | -0.22 | 0.827 | -0.00 | -0.01 – 0.01 | -0.09 | 0.928 | 0.00 | -0.01 – 0.01 | 0.60 | 0.545 |
| Interval Session Number | -0.02 *** | -0.03 – -0.02 | -10.57 | **<0.001** | -0.02 *** | -0.03 – -0.01 | -6.67 | **<0.001** | -0.02 *** | -0.03 – -0.02 | -7.71 | **<0.001** |
| Reward:Penalty | -0.00 * | -0.01 – -0.00 | -2.41 | **0.016** | -0.00 | -0.01 – 0.00 | -1.63 | 0.103 | -0.01 | -0.01 – 0.00 | -1.82 | 0.069 |
| **Random Effects** | | | | | | | | | | | | |
| $\sigma^2$ | 0.05 | | | | 0.05 | | | | 0.04 | | | |
| $\tau_{00}$ | 0.01 $_{SubID}$ | | | | 0.01 $_{SubID}$ | | | | 0.01 $_{SubID}$ | | | |
| $\tau_{11}$ | 0.00 $_{SubID.catPenaltyLevel1}$ | | | | 0.00 $_{SubID.catRewardLevel1}$ | | | | 0.00 $_{SubID.catRewardLevel1}$ | | | |
| $\rho_{01}$ | -0.12 $_{SubID}$ | | | | 0.79 $_{SubID}$ | | | | -0.02 $_{SubID}$ | | | |
| N | 94 $_{SubID}$ | | | | 94 $_{SubID}$ | | | | 94 $_{SubID}$ | | | |
| Observations | 11551 | | | | 5998 | | | | 5553 | | | |
| Marginal $R^2$ / Conditional $R^2$ | 0.010 / 0.140 | | | | 0.009 / 0.152 | | | | 0.013 / 0.186 | | | |

** p<0.05 ** p<0.01 *** p<0.001*

**Table S6:** *Inferior Frontal Gyrus Activity During the Cue and Interval Phase*. Linear mixed models of ROI during cue and interval task phases by reward and penalty incentive conditions, during all conditions, reward-varying conditions, and penalty-varying conditions.

**IFG Cue**

| | ROI - All Trials | | | | ROI - Vary Reward | | | | ROI - Vary Penalty | | | |
|---|---|---|---|---|---|---|---|---|---|---|---|---|
| *Predictors* | *Estimates* | *CI* | *Statistic* | *p* | *Estimates* | *CI* | *Statistic* | *p* | *Estimates* | *CI* | *Statistic* | *p* |
| (Intercept) | -0.11 *** | -0.16 – -0.06 | -4.15 | **<0.001** | -0.10 ** | -0.17 – -0.03 | -2.91 | **0.004** | -0.11 *** | -0.18 – -0.05 | -3.44 | **0.001** |
| Reward | -0.01 | -0.04 – 0.01 | -1.21 | 0.227 | -0.00 | -0.04 – 0.03 | -0.30 | 0.761 | -0.03 | -0.08 – 0.03 | -1.00 | 0.316 |
| Penalty | 0.04 ** | 0.01 – 0.07 | 2.85 | **0.004** | 0.07 ** | 0.02 – 0.11 | 2.71 | **0.007** | -0.01 | -0.06 – 0.04 | -0.34 | 0.731 |
| Interval Congruency | 0.01 | -0.01 – 0.04 | 1.09 | 0.276 | -0.00 | -0.03 – 0.03 | -0.01 | 0.990 | 0.03 | -0.00 – 0.06 | 1.77 | 0.076 |
| Interval Length | -0.01 | -0.03 – 0.02 | -0.41 | 0.683 | 0.00 | -0.03 – 0.03 | 0.17 | 0.865 | 0.02 | -0.02 – 0.07 | 1.00 | 0.316 |
| Interval Session Number | -0.05 *** | -0.07 – -0.03 | -4.12 | **<0.001** | -0.06 *** | -0.09 – -0.03 | -3.43 | **0.001** | -0.05 ** | -0.08 – -0.01 | -2.62 | **0.009** |
| Reward:Penalty | -0.01 | -0.03 – 0.02 | -0.54 | 0.588 | -0.01 | -0.04 – 0.02 | -0.58 | 0.563 | -0.00 | -0.04 – 0.03 | -0.29 | 0.769 |
| **Random Effects** | | | | | | | | | | | | |
| $\sigma^2$ | 1.54 | | | | 1.49 | | | | 1.48 | | | |
| $\tau_{00}$ | 0.05 SubID | | | | 0.09 SubID | | | | 0.07 SubID | | | |
| $\tau_{11}$ | 0.00 SubID.catRewardLevel1 | | | | 0.03 SubID.catPenaltyLevel1 | | | | 0.04 SubID.catRewardLevel1 | | | |
| | 0.01 SubID.catPenaltyLevel1 | | | | | | | | | | | |
| $\rho_{01}$ | 1.00 | | | | 0.20 SubID | | | | 0.15 SubID | | | |
| | -0.14 | | | | | | | | | | | |
| N | 94 SubID | | | | 94 SubID | | | | 94 SubID | | | |
| Observations | 11551 | | | | 5998 | | | | 5553 | | | |
| Marginal $R^2$ / Conditional $R^2$ | 0.003 / NA | | | | 0.005 / 0.077 | | | | 0.003 / 0.075 | | | |

** p<0.05 ** p<0.01 *** p<0.001*

**IFG Interval**

| | ROI - All Trials | | | | ROI - Vary Reward | | | | ROI - Vary Penalty | | | |
|---|---|---|---|---|---|---|---|---|---|---|---|---|
| *Predictors* | *Estimates* | *CI* | *Statistic* | *p* | *Estimates* | *CI* | *Statistic* | *p* | *Estimates* | *CI* | *Statistic* | *p* |
| (Intercept) | -0.16 *** | -0.17 – -0.14 | -19.67 | **<0.001** | -0.15 *** | -0.17 – -0.14 | -17.15 | **<0.001** | -0.16 *** | -0.18 – -0.14 | -18.62 | **<0.001** |
| Reward | -0.01 ** | -0.01 – -0.00 | -3.05 | **0.002** | -0.01 ** | -0.01 – -0.00 | -3.17 | **0.002** | -0.01 | -0.02 – 0.00 | -1.82 | 0.069 |
| Penalty | 0.01 *** | 0.00 – 0.02 | 3.39 | **0.001** | 0.01 ** | 0.01 – 0.02 | 3.03 | **0.002** | 0.00 | -0.01 – 0.01 | 0.40 | 0.692 |
| Interval Congruency | 0.00 | -0.00 – 0.01 | 1.76 | 0.079 | -0.00 | -0.01 – 0.01 | -0.08 | 0.940 | 0.01 ** | 0.00 – 0.01 | 2.70 | **0.007** |
| Interval Length | 0.00 | -0.00 – 0.01 | 0.42 | 0.675 | 0.00 | -0.00 – 0.01 | 0.80 | 0.422 | 0.01 | -0.00 – 0.01 | 1.28 | 0.202 |
| Interval Session Number | -0.01 *** | -0.02 – -0.01 | -6.18 | **<0.001** | -0.01 ** | -0.01 – -0.00 | -2.96 | **0.003** | -0.02 *** | -0.02 – -0.01 | -4.99 | **<0.001** |
| Reward:Penalty | 0.00 | -0.00 – 0.00 | 0.05 | 0.957 | 0.00 | -0.00 – 0.01 | 0.17 | 0.861 | -0.00 | -0.01 – 0.00 | -0.22 | 0.826 |
| **Random Effects** | | | | | | | | | | | | |
| $\sigma^2$ | 0.05 | | | | 0.04 | | | | 0.04 | | | |
| $\tau_{00}$ | 0.01 SubID | | | | 0.01 SubID | | | | 0.01 SubID | | | |
| $\tau_{11}$ | 0.00 SubID.catPenaltyLevel1 | | | | 0.00 SubID.catPenaltyLevel1 | | | | 0.00 SubID.catRewardLevel1 | | | |
| | 0.00 SubID.catRewardLevel1 | | | | 0.00 SubID.catRewardLevel1 | | | | | | | |
| $\rho_{01}$ | 0.25 | | | | 0.26 | | | | 0.01 SubID | | | |
| | 0.16 | | | | 0.72 | | | | | | | |
| N | 94 SubID | | | | 94 SubID | | | | 94 SubID | | | |
| Observations | 11551 | | | | 5998 | | | | 5553 | | | |
| Marginal $R^2$ / Conditional $R^2$ | 0.007 / 0.128 | | | | 0.007 / 0.164 | | | | 0.008 / 0.149 | | | |

** p<0.05 ** p<0.01 *** p<0.001*

**Table S7.** *Behavioral Task Performance*. Linear mixed models were conducted on response rate and log-transformed response time performance, and a generalized linear mixed effects model was conducted on accuracy performance. The models included congruency (at the interval and trial level), interval length, and interval or trial session number as covariates. Participants were faster and less accurate for high (relative to low) rewards, resulting in overall higher reward rates, whereas they were slower and more accurate for high (relative to low) penalties, resulting in overall lower reward rates.

**Behavioral Task Performance by Incentive Condition**

| | Response Rate | | | | Log RT | | | | Accuracy | | | |
|---|---|---|---|---|---|---|---|---|---|---|---|---|
| *Predictors* | *Estimates* | *CI* | *Statistic* | *p* | *Estimates* | *CI* | *Statistic* | *p* | *Odds Ratios* | *CI* | *Statistic* | *p* |
| (Intercept) | 1.50 *** | 1.44 – 1.56 | 52.29 | **<0.001** | 0.13 * | 0.03 – 0.23 | 2.54 | **0.011** | 24.81 *** | 21.60 – 28.50 | 45.38 | **<0.001** |
| Reward | 0.05 *** | 0.04 – 0.07 | 6.98 | **<0.001** | -0.12 *** | -0.16 – -0.09 | -7.47 | **<0.001** | 0.83 *** | 0.79 – 0.87 | -7.38 | **<0.001** |
| Penalty | -0.03 *** | -0.04 – -0.02 | -5.95 | **<0.001** | 0.09 *** | 0.06 – 0.11 | 7.28 | **<0.001** | 1.29 *** | 1.23 – 1.34 | 11.75 | **<0.001** |
| Interval Congruency | 0.02 *** | 0.01 – 0.02 | 6.36 | **<0.001** | | | | | | | | |
| Interval Length | 0.01 *** | 0.01 – 0.02 | 4.71 | **<0.001** | -0.02 *** | -0.03 – -0.02 | -7.98 | **<0.001** | 0.88 *** | 0.85 – 0.91 | -7.74 | **<0.001** |
| Interval Session Num | -0.00 | -0.01 – 0.00 | -1.26 | 0.208 | | | | | | | | |
| Reward:Penalty | 0.01 * | 0.00 – 0.02 | 2.33 | **0.020** | -0.04 *** | -0.06 – -0.02 | -3.50 | **<0.001** | 0.92 *** | 0.89 – 0.96 | -3.80 | **<0.001** |
| Reward:Interval Congruency | -0.00 | -0.01 – 0.00 | -1.11 | 0.269 | | | | | | | | |
| Penalty:Interval Congruency | -0.00 | -0.01 – 0.00 | -0.15 | 0.879 | | | | | | | | |
| Rew:Pen:Interval Congruency | -0.00 | -0.01 – 0.00 | -0.67 | 0.500 | | | | | | | | |
| Trial Congruency | | | | | -0.19 *** | -0.20 – -0.18 | -32.29 | **<0.001** | 1.13 *** | 1.06 – 1.20 | 3.59 | **<0.001** |
| Trial Session Num | | | | | -0.01 ** | -0.01 – -0.00 | -3.07 | **0.002** | 1.03 | 1.00 – 1.06 | 1.85 | 0.065 |
| Reward:Trial Congruency | | | | | 0.00 | -0.01 – 0.01 | 0.57 | 0.571 | 1.01 | 0.95 – 1.08 | 0.33 | 0.743 |
| Penalty:Trial Congruency | | | | | -0.00 | -0.01 – 0.01 | -0.11 | 0.912 | 0.94 | 0.88 – 1.00 | -1.87 | 0.061 |
| Rew:Pen:Trial Congruency | | | | | 0.00 | -0.01 – 0.02 | 0.77 | 0.439 | 1.00 | 0.94 – 1.07 | 0.09 | 0.925 |
| **Random Effects** | | | | | | | | | | | | |
| $\sigma^2$ | 0.08 | | | | 0.73 | | | | 3.29 | | | |
| $\tau_{00}$ | 0.08 $_{SubID}$ | | | | 0.24 $_{SubID}$ | | | | 0.42 $_{SubID}$ | | | |
| $\tau_{11}$ | 0.00 $_{SubID.catRewardLevel1}$ | | | | 0.02 $_{SubID.catRewardLevel1}$ | | | | 0.01 $_{SubID.catRewardLevel1}$ | | | |
| | 0.00 $_{SubID.catPenaltyLevel1}$ | | | | 0.01 $_{SubID.catPenaltyLevel1}$ | | | | | | | |
| | 0.00 $_{SubID.catRewardLevel1:catPenaltyLevel1}$ | | | | 0.01 $_{SubID.catRewardLevel1:catPenaltyLevel1}$ | | | | | | | |
| $\rho_{01}$ | -0.33 | | | | -0.34 | | | | -0.00 $_{SubID}$ | | | |
| | 0.11 | | | | 0.21 | | | | | | | |
| | -0.09 | | | | -0.25 | | | | | | | |
| N | 94 $_{SubID}$ | | | | 94 $_{SubID}$ | | | | 94 $_{SubID}$ | | | |
| Observations | 11552 | | | | 88901 | | | | 93152 | | | |
| Marginal $R^2$ / Conditional $R^2$ | 0.024 / 0.538 | | | | 0.030 / 0.304 | | | | 0.024 / 0.136 | | | |

** p<0.05 ** p<0.01 *** p<0.001*

**Table S8.** *Drift Diffusion Model Posterior Estimates.* We fit a drift diffusion model to RT and choice data, and estimated drift rate (v), response threshold (a), prepotent bias (z). The model estimated drift rate as a function of reward, penalty, and congruency, response threshold as a function of reward, penalty, and scaled linear running time (i.e., amount of time that passed within each interval which approximates an interval-level collapsing bound, z-scored), and bias as a function of congruency. Non-decision time was fitted as a free parameter.

Drift Diffusion Model Posterior Estimates

| parameter | condition | Mean | SD | pvalue | sig | CI_upper | CI_lower |
|---|---|---|---|---|---|---|---|
| Drift Rate | Intercept | 3.295 | 0.076 | 0.000 | *** | 3.445 | 3.146 |
| Drift Rate | Reward | 0.057 | 0.019 | 0.001 | ** | 0.094 | 0.021 |
| Drift Rate | Penalty | 0.029 | 0.016 | 0.032 | * | 0.061 | -0.002 |
| Drift Rate | Congruency | 0.197 | 0.017 | 0.000 | *** | 0.230 | 0.165 |
| Threshold | Intercept | 0.956 | 0.028 | 0.000 | *** | 1.014 | 0.903 |
| Threshold | Reward | -0.051 | 0.005 | 0.000 | *** | -0.041 | -0.062 |
| Threshold | Penalty | 0.051 | 0.004 | 0.000 | *** | 0.059 | 0.043 |
| Threshold | Scaled Running Time | -0.066 | 0.004 | 0.000 | *** | -0.059 | -0.073 |
| Bias | Congruency | 0.020 | 0.002 | 0.000 | *** | 0.025 | 0.015 |
| Drift Rate | Intertrial Variability | 0.708 | 0.028 | 0.000 | *** | 0.763 | 0.654 |
| NDT | Intertrial Variability | 0.177 | 0.002 | 0.000 | *** | 0.181 | 0.172 |
| NDT | Intercept | 0.272 | 0.006 | 0.000 | *** | 0.284 | 0.260 |

**Table S9:** *Drift Rate and Threshold Parameters by Reward and Penalty Incentive Condition.* Drift rate (v) and threshold (a) are plotted by each incentive condition.

Posterior Estimates by Incentive Condition

| parameter | Reward | Penalty | Mean | SD | CI_lower | CI_upper |
|---|---|---|---|---|---|---|
| Drift Rate | Low Reward | Low Penalty | 3.2097 | 0.0797 | 3.0540 | 3.3663 |
| Drift Rate | High Reward | Low Penalty | 3.3230 | 0.0805 | 3.1630 | 3.4799 |
| Drift Rate | Low Reward | High Penalty | 3.2679 | 0.0798 | 3.1122 | 3.4255 |
| Drift Rate | High Reward | High Penalty | 3.3812 | 0.0807 | 3.2240 | 3.5374 |
| Threshold | Low Reward | Low Penalty | 0.9559 | 0.0292 | 0.9013 | 1.0156 |
| Threshold | High Reward | Low Penalty | 0.8537 | 0.0291 | 0.7999 | 0.9135 |
| Threshold | Low Reward | High Penalty | 1.0574 | 0.0293 | 1.0033 | 1.1177 |
| Threshold | High Reward | High Penalty | 0.9552 | 0.0293 | 0.9010 | 1.0153 |

**Table S10:** Drift Diffusion Model with Ventral Striatum (Cue) Posterior Estimates.

Drift Diffusion Model with Ventral Striatum (Cue) Posterior Estimates

| parameter | condition | Mean | SD | pvalue | sig | CI_upper | CI_lower |
|---|---|---|---|---|---|---|---|
| Drift Rate | Intercept | 3.298 | 0.074 | 0.000 | *** | 3.444 | 3.153 |
| Drift Rate | Reward | 0.056 | 0.018 | 0.001 | ** | 0.092 | 0.020 |
| Drift Rate | Penalty | 0.028 | 0.016 | 0.038 | * | 0.059 | -0.003 |
| Drift Rate | Congruency | 0.197 | 0.017 | 0.000 | *** | 0.231 | 0.165 |
| Drift Rate | Striatum | 0.033 | 0.009 | 0.000 | *** | 0.051 | 0.014 |
| Drift Rate | Striatum x Reward | -0.010 | 0.008 | 0.111 | n.s. | 0.007 | -0.025 |
| Drift Rate | Striatum x Penalty | -0.001 | 0.008 | 0.457 | n.s. | 0.015 | -0.018 |
| Threshold | Intercept | 0.956 | 0.029 | 0.000 | *** | 1.014 | 0.901 |
| Threshold | Reward | -0.051 | 0.005 | 0.000 | *** | -0.040 | -0.061 |
| Threshold | Penalty | 0.051 | 0.004 | 0.000 | *** | 0.059 | 0.042 |
| Threshold | Striatum | 0.003 | 0.003 | 0.120 | n.s. | 0.008 | -0.002 |
| Threshold | Striatum x Reward | -0.007 | 0.002 | 0.000 | *** | -0.003 | -0.011 |
| Threshold | Striatum x Penalty | 0.004 | 0.002 | 0.013 | * | 0.008 | 0.001 |
| Threshold | Scaled Running Time | -0.067 | 0.004 | 0.000 | *** | -0.060 | -0.074 |
| Bias | Congruency | 0.020 | 0.002 | 0.000 | *** | 0.024 | 0.015 |
| Drift Rate | Intertrial Variability | 0.708 | 0.027 | 0.000 | *** | 0.763 | 0.655 |
| NDT | Intertrial Variability | 0.176 | 0.002 | 0.000 | *** | 0.181 | 0.172 |
| NDT | Intercept | 0.272 | 0.006 | 0.000 | *** | 0.284 | 0.260 |

**Table S11:** Drift Diffusion Model with Ventral Striatum (Interval) Posterior Estimates.

Drift Diffusion Model with Ventral Striatum (Interval) Posterior Estimates

| parameter | condition | Mean | SD | pvalue | sig | CI_upper | CI_lower |
|---|---|---|---|---|---|---|---|
| Drift Rate | Intercept | 3.268 | 0.076 | 0.000 | *** | 3.414 | 3.119 |
| Drift Rate | Reward | 0.063 | 0.020 | 0.001 | *** | 0.102 | 0.024 |
| Drift Rate | Penalty | 0.027 | 0.018 | 0.067 | n.s. | 0.062 | -0.008 |
| Drift Rate | Congruency | 0.200 | 0.017 | 0.000 | *** | 0.233 | 0.167 |
| Drift Rate | Striatum | 0.265 | 0.074 | 0.000 | *** | 0.411 | 0.121 |
| Drift Rate | Striatum x Reward | -0.045 | 0.052 | 0.185 | n.s. | 0.060 | -0.147 |
| Drift Rate | Striatum x Penalty | 0.006 | 0.051 | 0.454 | n.s. | 0.109 | -0.095 |
| Threshold | Intercept | 0.977 | 0.028 | 0.000 | *** | 1.034 | 0.923 |
| Threshold | Reward | -0.048 | 0.006 | 0.000 | *** | -0.037 | -0.060 |
| Threshold | Penalty | 0.051 | 0.005 | 0.000 | *** | 0.060 | 0.042 |
| Threshold | Striatum | -0.094 | 0.018 | 0.000 | *** | -0.061 | -0.130 |
| Threshold | Striatum x Reward | -0.020 | 0.015 | 0.090 | n.s. | 0.009 | -0.049 |
| Threshold | Striatum x Penalty | -0.001 | 0.012 | 0.476 | n.s. | 0.023 | -0.025 |
| Threshold | Scaled Running Time | -0.067 | 0.004 | 0.000 | *** | -0.060 | -0.074 |
| Bias | Congruency | 0.020 | 0.002 | 0.000 | *** | 0.024 | 0.015 |
| Drift Rate | Intertrial Variability | 0.711 | 0.027 | 0.000 | *** | 0.763 | 0.656 |
| NDT | Intertrial Variability | 0.176 | 0.002 | 0.000 | *** | 0.180 | 0.172 |
| NDT | Intercept | 0.272 | 0.006 | 0.000 | *** | 0.284 | 0.260 |

**Table S12:** Drift Diffusion Model with Anterior Insula (Cue) Posterior Estimates.

Drift Diffusion Model with Anterior Insula (Cue) Posterior Estimates

| parameter | condition | Mean | SD | pvalue | sig | CI_upper | CI_lower |
|---|---|---|---|---|---|---|---|
| Drift Rate | Intercept | 3.255 | 0.072 | 0.000 | *** | 3.398 | 3.114 |
| Drift Rate | Reward | 0.051 | 0.017 | 0.001 | ** | 0.084 | 0.019 |
| Drift Rate | Penalty | 0.008 | 0.015 | 0.284 | n.s. | 0.037 | -0.021 |
| Drift Rate | Congruency | 0.040 | 0.017 | 0.008 | ** | 0.074 | 0.007 |
| Drift Rate | AI | 0.011 | 0.008 | 0.072 | n.s. | 0.027 | -0.004 |
| Drift Rate | AI x Reward | -0.008 | 0.010 | 0.204 | n.s. | 0.011 | -0.027 |
| Drift Rate | AI x Penalty | 0.007 | 0.008 | 0.188 | n.s. | 0.021 | -0.009 |
| Threshold | Intercept | 0.919 | 0.026 | 0.000 | *** | 0.973 | 0.870 |
| Threshold | Reward | -0.049 | 0.005 | 0.000 | *** | -0.039 | -0.060 |
| Threshold | Penalty | 0.043 | 0.004 | 0.000 | *** | 0.051 | 0.036 |
| Threshold | AI | 0.003 | 0.002 | 0.107 | n.s. | 0.007 | -0.002 |
| Threshold | AI x Reward | -0.005 | 0.002 | 0.004 | ** | -0.002 | -0.009 |
| Threshold | AI x Penalty | 0.004 | 0.002 | 0.017 | * | 0.008 | 0.000 |
| Threshold | Scaled Running Time | -0.062 | 0.003 | 0.000 | *** | -0.056 | -0.069 |
| Bias | Congruency | 0.013 | 0.003 | 0.000 | *** | 0.018 | 0.008 |
| Drift Rate | Intertrial Variability | 0.697 | 0.026 | 0.000 | *** | 0.748 | 0.646 |
| NDT | Intertrial Variability | 0.168 | 0.002 | 0.000 | *** | 0.172 | 0.164 |
| NDT | Intercept | 0.264 | 0.006 | 0.000 | *** | 0.276 | 0.253 |

**Table S13:** Drift Diffusion Model with Anterior Insula (Interval) Posterior Estimates.

Drift Diffusion Model with Anterior Insula (Interval) Posterior Estimates

| parameter | condition | Mean | SD | pvalue | sig | CI_upper | CI_lower |
|---|---|---|---|---|---|---|---|
| Drift Rate | Intercept | 3.217 | 0.075 | 0.000 | *** | 3.368 | 3.070 |
| Drift Rate | Reward | 0.052 | 0.017 | 0.002 | ** | 0.086 | 0.018 |
| Drift Rate | Penalty | -0.006 | 0.016 | 0.360 | n.s. | 0.026 | -0.037 |
| Drift Rate | Congruency | 0.040 | 0.017 | 0.008 | ** | 0.073 | 0.007 |
| Drift Rate | AI | -0.222 | 0.057 | 0.000 | *** | -0.112 | -0.334 |
| Drift Rate | AI x Reward | 0.027 | 0.055 | 0.314 | n.s. | 0.136 | -0.080 |
| Drift Rate | AI x Penalty | -0.121 | 0.045 | 0.002 | ** | -0.036 | -0.210 |
| Threshold | Intercept | 0.920 | 0.026 | 0.000 | *** | 0.972 | 0.870 |
| Threshold | Reward | -0.052 | 0.005 | 0.000 | *** | -0.042 | -0.063 |
| Threshold | Penalty | 0.042 | 0.004 | 0.000 | *** | 0.050 | 0.034 |
| Threshold | AI | -0.006 | 0.013 | 0.319 | n.s. | 0.020 | -0.032 |
| Threshold | AI x Reward | -0.025 | 0.011 | 0.011 | * | -0.003 | -0.047 |
| Threshold | AI x Penalty | -0.008 | 0.010 | 0.224 | n.s. | 0.012 | -0.028 |
| Threshold | Scaled Running Time | -0.062 | 0.003 | 0.000 | *** | -0.056 | -0.069 |
| Bias | Congruency | 0.013 | 0.003 | 0.000 | *** | 0.018 | 0.008 |
| Drift Rate | Intertrial Variability | 0.687 | 0.028 | 0.000 | *** | 0.746 | 0.636 |
| NDT | Intertrial Variability | 0.168 | 0.002 | 0.000 | *** | 0.172 | 0.164 |
| NDT | Intercept | 0.264 | 0.006 | 0.000 | *** | 0.275 | 0.252 |

**Table S14:** Drift Diffusion Model with Caudal dACC (Cue) Posterior Estimates.

Drift Diffusion Model with Caudal dACC (Cue) Posterior Estimates

| parameter | condition | Mean | SD | pvalue | sig | CI_upper | CI_lower |
|---|---|---|---|---|---|---|---|
| Drift Rate | Intercept | 3.251 | 0.075 | 0.000 | *** | 3.400 | 3.105 |
| Drift Rate | Reward | 0.051 | 0.016 | 0.001 | ** | 0.083 | 0.018 |
| Drift Rate | Penalty | 0.005 | 0.015 | 0.356 | n.s. | 0.035 | -0.023 |
| Drift Rate | Congruency | 0.041 | 0.017 | 0.008 | ** | 0.075 | 0.007 |
| Drift Rate | Caudal dACC | 0.044 | 0.009 | 0.000 | *** | 0.062 | 0.027 |
| Drift Rate | Caudal dACC x Reward | 0.011 | 0.009 | 0.110 | n.s. | 0.028 | -0.006 |
| Drift Rate | Caudal dACC x Penalty | 0.002 | 0.008 | 0.403 | n.s. | 0.018 | -0.014 |
| Threshold | Intercept | 0.919 | 0.026 | 0.000 | *** | 0.971 | 0.870 |
| Threshold | Reward | -0.048 | 0.005 | 0.000 | *** | -0.038 | -0.058 |
| Threshold | Penalty | 0.042 | 0.004 | 0.000 | *** | 0.050 | 0.035 |
| Threshold | Caudal dACC | 0.000 | 0.002 | 0.498 | n.s. | 0.004 | -0.005 |
| Threshold | Caudal dACC x Reward | -0.003 | 0.002 | 0.020 | * | 0.000 | -0.007 |
| Threshold | Caudal dACC x Penalty | 0.004 | 0.002 | 0.018 | * | 0.008 | 0.000 |
| Threshold | Scaled Running Time | -0.062 | 0.003 | 0.000 | *** | -0.056 | -0.069 |
| Bias | Congruency | 0.013 | 0.003 | 0.000 | *** | 0.018 | 0.008 |
| Drift Rate | Intertrial Variability | 0.694 | 0.028 | 0.000 | *** | 0.749 | 0.640 |
| NDT | Intertrial Variability | 0.168 | 0.002 | 0.000 | *** | 0.172 | 0.164 |
| NDT | Intercept | 0.264 | 0.006 | 0.000 | *** | 0.276 | 0.253 |

**Table S15:** Drift Diffusion Model with Caudal dACC (Interval) Posterior Estimates.

Drift Diffusion Model with Caudal dACC (Interval) Posterior Estimates

| parameter | condition | Mean | SD | pvalue | sig | CI_upper | CI_lower |
|---|---|---|---|---|---|---|---|
| Drift Rate | Intercept | 3.254 | 0.075 | 0.000 | *** | 3.402 | 3.106 |
| Drift Rate | Reward | 0.059 | 0.019 | 0.000 | *** | 0.096 | 0.022 |
| Drift Rate | Penalty | 0.016 | 0.017 | 0.176 | n.s. | 0.050 | -0.017 |
| Drift Rate | Congruency | 0.039 | 0.017 | 0.011 | * | 0.073 | 0.006 |
| Drift Rate | Caudal dACC | -0.013 | 0.058 | 0.410 | n.s. | 0.103 | -0.128 |
| Drift Rate | Caudal dACC x Reward | -0.036 | 0.050 | 0.236 | n.s. | 0.060 | -0.135 |
| Drift Rate | Caudal dACC x Penalty | -0.042 | 0.041 | 0.140 | n.s. | 0.049 | -0.122 |
| Threshold | Intercept | 0.928 | 0.026 | 0.000 | *** | 0.981 | 0.878 |
| Threshold | Reward | -0.042 | 0.006 | 0.000 | *** | -0.031 | -0.054 |
| Threshold | Penalty | 0.042 | 0.004 | 0.000 | *** | 0.051 | 0.034 |
| Threshold | Caudal dACC | -0.028 | 0.015 | 0.034 | * | 0.002 | -0.059 |
| Threshold | Caudal dACC x Reward | -0.028 | 0.012 | 0.010 | * | -0.005 | -0.052 |
| Threshold | Caudal dACC x Penalty | -0.003 | 0.011 | 0.385 | n.s. | 0.019 | -0.023 |
| Threshold | Scaled Running Time | -0.063 | 0.003 | 0.000 | *** | -0.056 | -0.069 |
| Bias | Congruency | 0.013 | 0.003 | 0.000 | *** | 0.018 | 0.008 |
| Drift Rate | Intertrial Variability | 0.698 | 0.028 | 0.000 | *** | 0.753 | 0.644 |
| NDT | Intertrial Variability | 0.168 | 0.002 | 0.000 | *** | 0.172 | 0.164 |
| NDT | Intercept | 0.264 | 0.006 | 0.000 | *** | 0.276 | 0.253 |

**Table S16:** Drift Diffusion Model with Rostral dACC (Cue) Posterior Estimates.

Drift Diffusion Model with Rostral dACC (Cue) Posterior Estimates

| parameter | condition | Mean | SD | pvalue | sig | CI_upper | CI_lower |
|---|---|---|---|---|---|---|---|
| Drift Rate | Intercept | 3.247 | 0.073 | 0.000 | *** | 3.394 | 3.106 |
| Drift Rate | Reward | 0.051 | 0.016 | 0.002 | ** | 0.083 | 0.018 |
| Drift Rate | Penalty | 0.008 | 0.014 | 0.292 | n.s. | 0.036 | -0.020 |
| Drift Rate | Congruency | 0.040 | 0.017 | 0.008 | ** | 0.073 | 0.007 |
| Drift Rate | Rostral dACC | -0.003 | 0.009 | 0.349 | n.s. | 0.014 | -0.020 |
| Drift Rate | Rostral dACC x Reward | -0.014 | 0.009 | 0.058 | n.s. | 0.004 | -0.032 |
| Drift Rate | Rostral dACC x Penalty | 0.004 | 0.007 | 0.293 | n.s. | 0.018 | -0.011 |
| Threshold | Intercept | 0.920 | 0.026 | 0.000 | *** | 0.973 | 0.871 |
| Threshold | Reward | -0.049 | 0.005 | 0.000 | *** | -0.039 | -0.059 |
| Threshold | Penalty | 0.043 | 0.004 | 0.000 | *** | 0.050 | 0.035 |
| Threshold | Rostral dACC | 0.003 | 0.002 | 0.048 | * | 0.006 | -0.001 |
| Threshold | Rostral dACC x Reward | -0.004 | 0.002 | 0.020 | * | 0.000 | -0.007 |
| Threshold | Rostral dACC x Penalty | 0.002 | 0.002 | 0.099 | n.s. | 0.006 | -0.001 |
| Threshold | Scaled Running Time | -0.062 | 0.003 | 0.000 | *** | -0.056 | -0.069 |
| Bias | Congruency | 0.013 | 0.003 | 0.000 | *** | 0.018 | 0.008 |
| Drift Rate | Intertrial Variability | 0.693 | 0.027 | 0.000 | *** | 0.749 | 0.641 |
| NDT | Intertrial Variability | 0.168 | 0.002 | 0.000 | *** | 0.172 | 0.164 |
| NDT | Intercept | 0.264 | 0.006 | 0.000 | *** | 0.275 | 0.253 |

**Table S17:** Drift Diffusion Model with Rostral dACC (Interval) Posterior Estimates.

Drift Diffusion Model with Rostral dACC (Interval) Posterior Estimates

| parameter | condition | Mean | SD | pvalue | sig | CI_upper | CI_lower |
|---|---|---|---|---|---|---|---|
| Drift Rate | Intercept | 3.192 | 0.074 | 0.000 | *** | 3.336 | 3.047 |
| Drift Rate | Reward | 0.046 | 0.017 | 0.005 | ** | 0.079 | 0.012 |
| Drift Rate | Penalty | -0.001 | 0.016 | 0.480 | n.s. | 0.030 | -0.032 |
| Drift Rate | Congruency | 0.039 | 0.017 | 0.012 | * | 0.073 | 0.005 |
| Drift Rate | Rostral dACC | -0.250 | 0.058 | 0.000 | *** | -0.138 | -0.367 |
| Drift Rate | Rostral dACC x Reward | -0.019 | 0.050 | 0.350 | n.s. | 0.078 | -0.116 |
| Drift Rate | Rostral dACC x Penalty | -0.074 | 0.047 | 0.058 | n.s. | 0.016 | -0.168 |
| Threshold | Intercept | 0.920 | 0.026 | 0.000 | *** | 0.972 | 0.871 |
| Threshold | Reward | -0.052 | 0.005 | 0.000 | *** | -0.042 | -0.062 |
| Threshold | Penalty | 0.043 | 0.004 | 0.000 | *** | 0.051 | 0.036 |
| Threshold | Rostral dACC | 0.019 | 0.013 | 0.074 | n.s. | 0.044 | -0.007 |
| Threshold | Rostral dACC x Reward | -0.022 | 0.010 | 0.018 | * | -0.001 | -0.043 |
| Threshold | Rostral dACC x Penalty | 0.006 | 0.011 | 0.273 | n.s. | 0.027 | -0.014 |
| Threshold | Scaled Running Time | -0.063 | 0.003 | 0.000 | *** | -0.056 | -0.069 |
| Bias | Congruency | 0.013 | 0.003 | 0.000 | *** | 0.018 | 0.008 |
| Drift Rate | Intertrial Variability | 0.680 | 0.029 | 0.000 | *** | 0.736 | 0.624 |
| NDT | Intertrial Variability | 0.169 | 0.002 | 0.000 | *** | 0.172 | 0.164 |
| NDT | Intercept | 0.265 | 0.006 | 0.000 | *** | 0.276 | 0.254 |

**Table S18:** Drift Diffusion Model with Dorsolateral PFC (Cue) Posterior Estimates.

Drift Diffusion Model with Dorsolateral PFC (Cue) Posterior Estimates

| parameter | condition | Mean | SD | pvalue | sig | CI_upper | CI_lower |
|---|---|---|---|---|---|---|---|
| Drift Rate | Intercept | 3.244 | 0.072 | 0.000 | *** | 3.387 | 3.101 |
| Drift Rate | Reward | 0.053 | 0.016 | 0.001 | *** | 0.084 | 0.021 |
| Drift Rate | Penalty | 0.006 | 0.015 | 0.341 | n.s. | 0.035 | -0.023 |
| Drift Rate | Congruency | 0.040 | 0.017 | 0.008 | ** | 0.073 | 0.007 |
| Drift Rate | DLPFC | 0.026 | 0.008 | 0.000 | *** | 0.041 | 0.011 |
| Drift Rate | DLPFC x Reward | 0.013 | 0.009 | 0.078 | n.s. | 0.030 | -0.005 |
| Drift Rate | DLPFC x Penalty | 0.005 | 0.007 | 0.258 | n.s. | 0.019 | -0.010 |
| Threshold | Intercept | 0.917 | 0.026 | 0.000 | *** | 0.969 | 0.868 |
| Threshold | Reward | -0.049 | 0.005 | 0.000 | *** | -0.039 | -0.059 |
| Threshold | Penalty | 0.043 | 0.004 | 0.000 | *** | 0.050 | 0.035 |
| Threshold | DLPFC | 0.001 | 0.002 | 0.370 | n.s. | 0.004 | -0.003 |
| Threshold | DLPFC x Reward | -0.004 | 0.002 | 0.012 | * | -0.001 | -0.007 |
| Threshold | DLPFC x Penalty | 0.005 | 0.002 | 0.005 | ** | 0.008 | 0.001 |
| Threshold | Scaled Running Time | -0.062 | 0.003 | 0.000 | *** | -0.056 | -0.069 |
| Bias | Congruency | 0.013 | 0.003 | 0.000 | *** | 0.018 | 0.008 |
| Drift Rate | Intertrial Variability | 0.691 | 0.027 | 0.000 | *** | 0.744 | 0.637 |
| NDT | Intertrial Variability | 0.168 | 0.002 | 0.000 | *** | 0.172 | 0.164 |
| NDT | Intercept | 0.264 | 0.006 | 0.000 | *** | 0.276 | 0.253 |

**Table S19:** Drift Diffusion Model with Dorsolateral PFC (Interval) Posterior Estimates.

Drift Diffusion Model with Dorsolateral PFC (Interval) Posterior Estimates

| parameter | condition | Mean | SD | pvalue | sig | CI_upper | CI_lower |
|---|---|---|---|---|---|---|---|
| Drift Rate | Intercept | 3.240 | 0.072 | 0.000 | *** | 3.384 | 3.102 |
| Drift Rate | Reward | 0.048 | 0.016 | 0.002 | ** | 0.081 | 0.016 |
| Drift Rate | Penalty | 0.006 | 0.015 | 0.344 | n.s. | 0.036 | -0.023 |
| Drift Rate | Congruency | 0.039 | 0.017 | 0.010 | * | 0.073 | 0.006 |
| Drift Rate | DLPFC | -0.136 | 0.051 | 0.003 | ** | -0.039 | -0.237 |
| Drift Rate | DLPFC x Reward | -0.011 | 0.049 | 0.404 | n.s. | 0.087 | -0.106 |
| Drift Rate | DLPFC x Penalty | -0.034 | 0.042 | 0.216 | n.s. | 0.043 | -0.116 |
| Threshold | Intercept | 0.919 | 0.026 | 0.000 | *** | 0.972 | 0.871 |
| Threshold | Reward | -0.050 | 0.005 | 0.000 | *** | -0.040 | -0.060 |
| Threshold | Penalty | 0.043 | 0.004 | 0.000 | *** | 0.050 | 0.035 |
| Threshold | DLPFC | 0.008 | 0.013 | 0.269 | n.s. | 0.034 | -0.019 |
| Threshold | DLPFC x Reward | -0.027 | 0.012 | 0.010 | * | -0.004 | -0.050 |
| Threshold | DLPFC x Penalty | 0.012 | 0.011 | 0.125 | n.s. | 0.033 | -0.008 |
| Threshold | Scaled Running Time | -0.063 | 0.003 | 0.000 | *** | -0.056 | -0.069 |
| Bias | Congruency | 0.013 | 0.003 | 0.000 | *** | 0.018 | 0.008 |
| Drift Rate | Intertrial Variability | 0.692 | 0.026 | 0.000 | *** | 0.746 | 0.643 |
| NDT | Intertrial Variability | 0.168 | 0.002 | 0.000 | *** | 0.172 | 0.164 |
| NDT | Intercept | 0.264 | 0.006 | 0.000 | *** | 0.276 | 0.253 |

**Table S20:** Drift Diffusion Model with Inferior Frontal Gyrus (Cue) Posterior Estimates.

Drift Diffusion Model with Inferior Frontal Gyrus (Cue) Posterior Estimates

| parameter | condition | Mean | SD | pvalue | sig | CI_upper | CI_lower |
|---|---|---|---|---|---|---|---|
| Drift Rate | Intercept | 3.255 | 0.073 | 0.000 | *** | 3.399 | 3.112 |
| Drift Rate | Reward | 0.051 | 0.016 | 0.001 | *** | 0.082 | 0.019 |
| Drift Rate | Penalty | 0.007 | 0.015 | 0.312 | n.s. | 0.035 | -0.022 |
| Drift Rate | Congruency | 0.041 | 0.017 | 0.008 | ** | 0.075 | 0.007 |
| Drift Rate | IFG | -0.008 | 0.008 | 0.163 | n.s. | 0.008 | -0.023 |
| Drift Rate | IFG x Reward | 0.050 | 0.010 | 0.000 | *** | 0.070 | 0.029 |
| Drift Rate | IFG x Penalty | -0.005 | 0.008 | 0.260 | n.s. | 0.010 | -0.020 |
| Threshold | Intercept | 0.920 | 0.026 | 0.000 | *** | 0.972 | 0.870 |
| Threshold | Reward | -0.049 | 0.005 | 0.000 | *** | -0.039 | -0.059 |
| Threshold | Penalty | 0.043 | 0.004 | 0.000 | *** | 0.050 | 0.035 |
| Threshold | IFG | 0.003 | 0.002 | 0.066 | n.s. | 0.007 | -0.001 |
| Threshold | IFG x Reward | -0.004 | 0.002 | 0.026 | * | 0.000 | -0.007 |
| Threshold | IFG x Penalty | 0.002 | 0.002 | 0.178 | n.s. | 0.005 | -0.002 |
| Threshold | Scaled Running Time | -0.062 | 0.003 | 0.000 | *** | -0.056 | -0.069 |
| Bias | Congruency | 0.013 | 0.003 | 0.000 | *** | 0.018 | 0.008 |
| Drift Rate | Intertrial Variability | 0.698 | 0.027 | 0.000 | *** | 0.751 | 0.644 |
| NDT | Intertrial Variability | 0.168 | 0.002 | 0.000 | *** | 0.172 | 0.164 |
| NDT | Intercept | 0.264 | 0.006 | 0.000 | *** | 0.275 | 0.252 |

**Table S21:** Drift Diffusion Model with Inferior Frontal Gyrus (Interval) Posterior Estimates.

Drift Diffusion Model with Inferior Frontal Gyrus (Interval) Posterior Estimates

| parameter | condition | Mean | SD | pvalue | sig | CI_upper | CI_lower |
|---|---|---|---|---|---|---|---|
| Drift Rate | Intercept | 3.214 | 0.074 | 0.000 | *** | 3.359 | 3.068 |
| Drift Rate | Reward | 0.052 | 0.018 | 0.001 | ** | 0.087 | 0.017 |
| Drift Rate | Penalty | 0.001 | 0.016 | 0.482 | n.s. | 0.033 | -0.031 |
| Drift Rate | Congruency | 0.039 | 0.017 | 0.011 | * | 0.073 | 0.006 |
| Drift Rate | IFG | -0.185 | 0.056 | 0.000 | *** | -0.075 | -0.295 |
| Drift Rate | IFG x Reward | 0.265 | 0.053 | 0.000 | *** | 0.371 | 0.158 |
| Drift Rate | IFG x Penalty | -0.055 | 0.047 | 0.118 | n.s. | 0.032 | -0.148 |
| Threshold | Intercept | 0.926 | 0.026 | 0.000 | *** | 0.978 | 0.876 |
| Threshold | Reward | -0.050 | 0.005 | 0.000 | *** | -0.040 | -0.061 |
| Threshold | Penalty | 0.043 | 0.004 | 0.000 | *** | 0.051 | 0.035 |
| Threshold | IFG | 0.036 | 0.013 | 0.003 | ** | 0.063 | 0.011 |
| Threshold | IFG x Reward | -0.010 | 0.010 | 0.149 | n.s. | 0.009 | -0.030 |
| Threshold | IFG x Penalty | 0.005 | 0.010 | 0.301 | n.s. | 0.026 | -0.015 |
| Threshold | Scaled Running Time | -0.063 | 0.003 | 0.000 | *** | -0.056 | -0.069 |
| Bias | Congruency | 0.013 | 0.003 | 0.000 | *** | 0.018 | 0.008 |
| Drift Rate | Intertrial Variability | 0.688 | 0.028 | 0.000 | *** | 0.741 | 0.633 |
| NDT | Intertrial Variability | 0.168 | 0.002 | 0.000 | *** | 0.172 | 0.164 |
| NDT | Intercept | 0.264 | 0.006 | 0.000 | *** | 0.276 | 0.253 |

**Table S22:** *Task Performance Predicted by Ventral Striatum during Cue and Interval Phase.*

**Task Performance Predicted by Ventral Striatum (Cue)**

| | Response Rate | | | | Log RT | | | | Accuracy | | | |
|---|---|---|---|---|---|---|---|---|---|---|---|---|
| *Predictors* | *Estimates* | *CI* | *Statistic* | *p* | *Estimates* | *CI* | *Statistic* | *p* | *Odds Ratios* | *CI* | *Statistic* | *p* |
| (Intercept) | 1.50 *** | 1.44 – 1.56 | 52.44 | **<0.001** | 0.12 * | 0.02 – 0.22 | 2.39 | **0.017** | 24.65 *** | 21.46 – 28.31 | 45.34 | **<0.001** |
| Reward | 0.05 *** | 0.05 – 0.06 | 19.50 | **<0.001** | -0.11 *** | -0.11 – -0.10 | -36.24 | **<0.001** | 0.83 *** | 0.81 – 0.86 | -11.11 | **<0.001** |
| Penalty | -0.03 *** | -0.03 – -0.02 | -10.11 | **<0.001** | 0.08 *** | 0.07 – 0.08 | 24.18 | **<0.001** | 1.26 *** | 1.21 – 1.30 | 13.09 | **<0.001** |
| Ventral Striatum (VS) | 0.01 ** | 0.00 – 0.01 | 2.64 | **0.008** | -0.01 *** | -0.01 – -0.00 | -3.45 | **0.001** | 1.01 | 0.98 – 1.04 | 0.82 | 0.413 |
| Interval Congruency | 0.02 *** | 0.01 – 0.02 | 6.35 | **<0.001** | | | | | | | | |
| Interval Length | 0.01 *** | 0.01 – 0.02 | 5.06 | **<0.001** | -0.03 *** | -0.03 – -0.02 | -8.71 | **<0.001** | 0.88 *** | 0.85 – 0.91 | -7.73 | **<0.001** |
| Interval Session Number | -0.00 | -0.01 – 0.00 | -1.21 | 0.228 | -0.01 * | -0.01 – -0.00 | -2.54 | **0.011** | 1.03 * | 1.00 – 1.07 | 2.12 | **0.034** |
| Reward:Penalty | 0.01 ** | 0.00 – 0.01 | 3.22 | **0.001** | -0.02 *** | -0.03 – -0.02 | -7.74 | **<0.001** | 0.93 *** | 0.90 – 0.96 | -4.68 | **<0.001** |
| VS:Penalty | -0.00 * | -0.01 – -0.00 | -2.12 | **0.034** | 0.01 ** | 0.00 – 0.01 | 2.80 | **0.005** | 1.01 | 0.98 – 1.03 | 0.61 | 0.543 |
| VS:Reward | 0.00 | -0.00 – 0.01 | 1.29 | 0.196 | -0.01 ** | -0.01 – -0.00 | -2.87 | **0.004** | 0.98 | 0.95 – 1.01 | -1.58 | 0.115 |
| Congruency | | | | | -0.19 *** | -0.20 – -0.18 | -31.82 | **<0.001** | 1.14 *** | 1.07 – 1.21 | 3.89 | **<0.001** |
| **Random Effects** | | | | | | | | | | | | |
| $\sigma^2$ | 0.08 | | | | 0.76 | | | | 3.29 | | | |
| $\tau_{00}$ | 0.08 $_{SubID}$ | | | | 0.23 $_{SubID}$ | | | | 0.42 $_{SubID}$ | | | |
| N | 94 $_{SubID}$ | | | | 94 $_{SubID}$ | | | | 94 $_{SubID}$ | | | |
| Observations | 11551 | | | | 88901 | | | | 93150 | | | |
| Marginal $R^2$ / Conditional $R^2$ | 0.025 / 0.494 | | | | 0.026 / 0.254 | | | | 0.025 / 0.135 | | | |

* p<0.05 ** p<0.01 *** p<0.001

**Task Performance Predicted by Ventral Striatum (Interval)**

| | Response Rate | | | | Log RT | | | | Accuracy | | | |
|---|---|---|---|---|---|---|---|---|---|---|---|---|
| *Predictors* | *Estimates* | *CI* | *Statistic* | *p* | *Estimates* | *CI* | *Statistic* | *p* | *Odds Ratios* | *CI* | *Statistic* | *p* |
| (Intercept) | 1.48 *** | 1.42 – 1.53 | 51.75 | **<0.001** | 0.16 ** | 0.06 – 0.26 | 3.24 | **0.001** | 24.74 *** | 21.48 – 28.50 | 44.51 | **<0.001** |
| Reward | 0.05 *** | 0.05 – 0.06 | 15.12 | **<0.001** | -0.11 *** | -0.11 – -0.10 | -27.40 | **<0.001** | 0.84 *** | 0.81 – 0.88 | -8.04 | **<0.001** |
| Penalty | -0.03 *** | -0.04 – -0.02 | -8.08 | **<0.001** | 0.08 *** | 0.07 – 0.09 | 19.84 | **<0.001** | 1.24 *** | 1.19 – 1.30 | 9.70 | **<0.001** |
| Ventral Striatum (VS) | 0.12 *** | 0.10 – 0.15 | 9.25 | **<0.001** | -0.23 *** | -0.25 – -0.20 | -15.50 | **<0.001** | 0.98 | 0.84 – 1.14 | -0.27 | 0.791 |
| Interval Congruency | 0.02 *** | 0.01 – 0.02 | 6.41 | **<0.001** | | | | | | | | |
| Interval Length | 0.01 *** | 0.01 – 0.02 | 5.11 | **<0.001** | -0.03 *** | -0.03 – -0.02 | -8.84 | **<0.001** | 0.88 *** | 0.85 – 0.91 | -7.75 | **<0.001** |
| Interval Session Number | -0.00 | -0.01 – 0.00 | -1.47 | 0.142 | -0.01 * | -0.01 – -0.00 | -2.02 | **0.043** | 1.04 * | 1.00 – 1.07 | 2.19 | **0.029** |
| Reward:Penalty | 0.01 ** | 0.00 – 0.01 | 3.16 | **0.002** | -0.02 *** | -0.03 – -0.02 | -7.62 | **<0.001** | 0.93 *** | 0.90 – 0.96 | -4.74 | **<0.001** |
| VS:Penalty | -0.00 | -0.02 – 0.02 | -0.02 | 0.987 | -0.01 | -0.04 – 0.01 | -1.01 | 0.311 | 1.06 | 0.92 – 1.22 | 0.75 | 0.453 |
| VS:Reward | -0.00 | -0.03 – 0.02 | -0.39 | 0.695 | -0.00 | -0.03 – 0.03 | -0.07 | 0.941 | 0.96 | 0.84 – 1.11 | -0.54 | 0.590 |
| Congruency | | | | | -0.19 *** | -0.20 – -0.18 | -31.89 | **<0.001** | 1.14 *** | 1.07 – 1.21 | 3.90 | **<0.001** |
| **Random Effects** | | | | | | | | | | | | |
| $\sigma^2$ | 0.08 | | | | 0.76 | | | | 3.29 | | | |
| $\tau_{00}$ | 0.08 $_{SubID}$ | | | | 0.23 $_{SubID}$ | | | | 0.42 $_{SubID}$ | | | |
| N | 94 $_{SubID}$ | | | | 94 $_{SubID}$ | | | | 94 $_{SubID}$ | | | |
| Observations | 11551 | | | | 88901 | | | | 93150 | | | |
| Marginal $R^2$ / Conditional $R^2$ | 0.029 / 0.494 | | | | 0.028 / 0.254 | | | | 0.024 / 0.134 | | | |

* p<0.05 ** p<0.01 *** p<0.001

**Table S23:** *Task Performance Predicted by Anterior Insula during Cue and Interval Phase.*

**Task Performance Predicted by Anterior Insula (Cue)**

| | Response Rate | | | | Log RT | | | | Accuracy | | | |
|---|---|---|---|---|---|---|---|---|---|---|---|---|
| *Predictors* | *Estimates* | *CI* | *Statistic* | *p* | *Estimates* | *CI* | *Statistic* | *p* | *Odds Ratios* | *CI* | *Statistic* | *p* |
| (Intercept) | 1.50 *** | 1.45 – 1.56 | 52.45 | **<0.001** | 0.12 * | 0.02 – 0.22 | 2.37 | **0.018** | 24.83 *** | 21.62 – 28.52 | 45.40 | **<0.001** |
| Reward | 0.05 *** | 0.05 – 0.06 | 19.53 | **<0.001** | -0.11 *** | -0.11 – -0.10 | -36.47 | **<0.001** | 0.83 *** | 0.81 – 0.86 | -11.22 | **<0.001** |
| Penalty | -0.03 *** | -0.04 – -0.02 | -10.20 | **<0.001** | 0.08 *** | 0.07 – 0.08 | 24.23 | **<0.001** | 1.26 *** | 1.22 – 1.31 | 13.25 | **<0.001** |
| Anterior Insula (AI) | 0.00 | -0.00 – 0.01 | 1.41 | 0.160 | -0.00 | -0.01 – 0.00 | -0.51 | 0.612 | 1.04 *** | 1.02 – 1.07 | 3.42 | **0.001** |
| Interval Congruency | 0.02 *** | 0.01 – 0.02 | 6.34 | **<0.001** | | | | | | | | |
| Interval Length | 0.01 *** | 0.01 – 0.02 | 5.03 | **<0.001** | -0.03 *** | -0.03 – -0.02 | -8.67 | **<0.001** | 0.88 *** | 0.85 – 0.91 | -7.77 | **<0.001** |
| Interval Session Number | -0.00 | -0.01 – 0.00 | -1.21 | 0.226 | -0.01 * | -0.01 – -0.00 | -2.48 | **0.013** | 1.04 * | 1.01 – 1.07 | 2.30 | **0.021** |
| Reward:Penalty | 0.01 ** | 0.00 – 0.01 | 3.22 | **0.001** | -0.02 *** | -0.03 – -0.02 | -7.69 | **<0.001** | 0.93 *** | 0.90 – 0.96 | -4.63 | **<0.001** |
| AI:Penalty | -0.00 | -0.01 – 0.00 | -0.82 | 0.414 | 0.00 | -0.00 – 0.01 | 1.33 | 0.185 | 1.02 * | 1.00 – 1.05 | 2.00 | **0.045** |
| AI:Reward | 0.00 | -0.00 – 0.00 | 0.01 | 0.989 | -0.00 | -0.01 – 0.00 | -1.73 | 0.084 | 0.98 | 0.95 – 1.00 | -1.89 | 0.059 |
| Congruency | | | | | -0.19 *** | -0.20 – -0.18 | -31.80 | **<0.001** | 1.14 *** | 1.07 – 1.21 | 3.90 | **<0.001** |
| **Random Effects** | | | | | | | | | | | | |
| $\sigma^2$ | 0.08 | | | | 0.76 | | | | 3.29 | | | |
| $\tau_{00}$ | 0.08 $_{SubID}$ | | | | 0.23 $_{SubID}$ | | | | 0.42 $_{SubID}$ | | | |
| N | 94 $_{SubID}$ | | | | 94 $_{SubID}$ | | | | 94 $_{SubID}$ | | | |
| Observations | 11551 | | | | 88901 | | | | 93150 | | | |
| Marginal $R^2$ / Conditional $R^2$ | 0.024 / 0.493 | | | | 0.026 / 0.254 | | | | 0.026 / 0.136 | | | |

** p<0.05 ** p<0.01 *** p<0.001*

**Task Performance Predicted by Anterior Insula (Interval)**

| | Response Rate | | | | Log RT | | | | Accuracy | | | |
|---|---|---|---|---|---|---|---|---|---|---|---|---|
| *Predictors* | *Estimates* | *CI* | *Statistic* | *p* | *Estimates* | *CI* | *Statistic* | *p* | *Odds Ratios* | *CI* | *Statistic* | *p* |
| (Intercept) | 1.50 *** | 1.44 – 1.55 | 52.20 | **<0.001** | 0.13 * | 0.03 – 0.22 | 2.54 | **0.011** | 25.68 *** | 22.32 – 29.54 | 45.36 | **<0.001** |
| Reward | 0.06 *** | 0.05 – 0.06 | 18.29 | **<0.001** | -0.11 *** | -0.12 – -0.11 | -33.52 | **<0.001** | 0.82 *** | 0.79 – 0.86 | -10.15 | **<0.001** |
| Penalty | -0.03 *** | -0.04 – -0.03 | -9.87 | **<0.001** | 0.08 *** | 0.07 – 0.09 | 22.06 | **<0.001** | 1.25 *** | 1.20 – 1.30 | 11.13 | **<0.001** |
| Anterior Insula (AI) | -0.04 ** | -0.06 – -0.01 | -3.21 | **0.001** | 0.06 *** | 0.03 – 0.08 | 4.39 | **<0.001** | 1.32 *** | 1.15 – 1.52 | 3.89 | **<0.001** |
| Interval Congruency | 0.02 *** | 0.01 – 0.02 | 6.41 | **<0.001** | | | | | | | | |
| Interval Length | 0.01 *** | 0.01 – 0.02 | 5.02 | **<0.001** | -0.03 *** | -0.03 – -0.02 | -8.65 | **<0.001** | 0.88 *** | 0.85 – 0.91 | -7.73 | **<0.001** |
| Interval Session Number | -0.00 | -0.01 – 0.00 | -1.57 | 0.117 | -0.01 * | -0.01 – -0.00 | -1.99 | **0.047** | 1.04 * | 1.01 – 1.07 | 2.54 | **0.011** |
| Reward:Penalty | 0.01 ** | 0.00 – 0.01 | 3.04 | **0.002** | -0.02 *** | -0.03 – -0.02 | -7.50 | **<0.001** | 0.93 *** | 0.90 – 0.96 | -4.59 | **<0.001** |
| AI:Penalty | -0.02 * | -0.05 – -0.00 | -2.11 | **0.035** | 0.03 * | 0.00 – 0.05 | 2.07 | **0.038** | 0.98 | 0.86 – 1.12 | -0.33 | 0.741 |
| AI:Reward | 0.02 * | 0.00 – 0.05 | 2.06 | **0.039** | -0.05 *** | -0.07 – -0.03 | -3.96 | **<0.001** | 0.92 | 0.81 – 1.05 | -1.28 | 0.202 |
| Congruency | | | | | -0.19 *** | -0.20 – -0.18 | -31.84 | **<0.001** | 1.13 *** | 1.06 – 1.21 | 3.88 | **<0.001** |
| **Random Effects** | | | | | | | | | | | | |
| $\sigma^2$ | 0.08 | | | | 0.76 | | | | 3.29 | | | |
| $\tau_{00}$ | 0.08 $_{SubID}$ | | | | 0.23 $_{SubID}$ | | | | 0.42 $_{SubID}$ | | | |
| N | 94 $_{SubID}$ | | | | 94 $_{SubID}$ | | | | 94 $_{SubID}$ | | | |
| Observations | 11551 | | | | 88901 | | | | 93150 | | | |
| Marginal $R^2$ / Conditional $R^2$ | 0.025 / 0.494 | | | | 0.026 / 0.254 | | | | 0.025 / 0.136 | | | |

** p<0.05 ** p<0.01 *** p<0.001*

**Table S24:** *Task Performance Predicted by Caudal dACC during Cue and Interval Phase.*

**Task Performance Predicted by Caudal dACC (Cue)**

| | Response Rate | | | | Log RT | | | | Accuracy | | | |
|---|---|---|---|---|---|---|---|---|---|---|---|---|
| *Predictors* | *Estimates* | *CI* | *Statistic* | *p* | *Estimates* | *CI* | *Statistic* | *p* | *Odds Ratios* | *CI* | *Statistic* | *p* |
| (Intercept) | 1.50 *** | 1.44 – 1.56 | 52.47 | **<0.001** | 0.12 * | 0.02 – 0.22 | 2.41 | **0.016** | 24.61 *** | 21.42 – 28.27 | 45.27 | **<0.001** |
| Reward | 0.05 *** | 0.05 – 0.06 | 19.42 | **<0.001** | -0.11 *** | -0.11 – -0.10 | -36.09 | **<0.001** | 0.83 *** | 0.81 – 0.86 | -11.13 | **<0.001** |
| Penalty | -0.03 *** | -0.03 – -0.02 | -10.15 | **<0.001** | 0.08 *** | 0.07 – 0.08 | 24.16 | **<0.001** | 1.26 *** | 1.21 – 1.30 | 13.06 | **<0.001** |
| Caudal dACC | 0.01 *** | 0.01 – 0.02 | 6.18 | **<0.001** | -0.02 *** | -0.02 – -0.01 | -7.13 | **<0.001** | 1.03 * | 1.01 – 1.06 | 2.38 | **0.017** |
| Interval Congruency | 0.02 *** | 0.01 – 0.02 | 6.34 | **<0.001** | | | | | | | | |
| Interval Length | 0.01 *** | 0.01 – 0.02 | 4.96 | **<0.001** | -0.03 *** | -0.03 – -0.02 | -8.58 | **<0.001** | 0.88 *** | 0.85 – 0.91 | -7.77 | **<0.001** |
| Interval Session Number | -0.00 | -0.01 – 0.00 | -1.20 | 0.228 | -0.01 * | -0.01 – -0.00 | -2.48 | **0.013** | 1.04 * | 1.00 – 1.07 | 2.20 | **0.028** |
| Reward:Penalty | 0.01 *** | 0.00 – 0.01 | 3.33 | **0.001** | -0.02 *** | -0.03 – -0.02 | -7.90 | **<0.001** | 0.93 *** | 0.90 – 0.96 | -4.66 | **<0.001** |
| Caudal dACC:Penalty | -0.00 | -0.01 – 0.00 | -1.09 | 0.275 | 0.00 * | 0.00 – 0.01 | 2.00 | **0.045** | 1.01 | 0.99 – 1.04 | 1.03 | 0.301 |
| Caudal dACC:Reward | 0.00 * | 0.00 – 0.01 | 2.02 | **0.043** | -0.01 ** | -0.01 – -0.00 | -2.98 | **0.003** | 0.99 | 0.96 – 1.01 | -0.85 | 0.395 |
| Congruency | | | | | -0.19 *** | -0.20 – -0.18 | -31.81 | **<0.001** | 1.14 *** | 1.07 – 1.21 | 3.90 | **<0.001** |
| **Random Effects** | | | | | | | | | | | | |
| $\sigma^2$ | 0.08 | | | | 0.76 | | | | 3.29 | | | |
| $\tau_{00}$ | 0.08 SubID | | | | 0.23 SubID | | | | 0.42 SubID | | | |
| N | 94 SubID | | | | 94 SubID | | | | 94 SubID | | | |
| Observations | 11551 | | | | 88901 | | | | 93150 | | | |
| Marginal $R^2$ / Conditional $R^2$ | 0.026 / 0.495 | | | | 0.026 / 0.254 | | | | 0.025 / 0.135 | | | |

* p<0.05 ** p<0.01 *** p<0.001

**Task Performance Predicted by Caudal dACC (Interval)**

| | Response Rate | | | | Log RT | | | | Accuracy | | | |
|---|---|---|---|---|---|---|---|---|---|---|---|---|
| *Predictors* | *Estimates* | *CI* | *Statistic* | *p* | *Estimates* | *CI* | *Statistic* | *p* | *Odds Ratios* | *CI* | *Statistic* | *p* |
| (Intercept) | 1.50 *** | 1.44 – 1.55 | 51.92 | **<0.001** | 0.13 ** | 0.03 – 0.23 | 2.65 | **0.008** | 24.38 *** | 21.13 – 28.13 | 43.79 | **<0.001** |
| Reward | 0.05 *** | 0.04 – 0.06 | 12.80 | **<0.001** | -0.10 *** | -0.11 – -0.09 | -23.26 | **<0.001** | 0.85 *** | 0.81 – 0.89 | -6.91 | **<0.001** |
| Penalty | -0.03 *** | -0.04 – -0.02 | -7.09 | **<0.001** | 0.08 *** | 0.07 – 0.09 | 17.86 | **<0.001** | 1.27 *** | 1.21 – 1.33 | 9.95 | **<0.001** |
| Caudal dACC | 0.03 | -0.00 – 0.05 | 1.95 | 0.051 | -0.06 *** | -0.09 – -0.03 | -4.13 | **<0.001** | 1.05 | 0.90 – 1.22 | 0.66 | 0.507 |
| Interval Congruency | 0.02 *** | 0.01 – 0.02 | 6.38 | **<0.001** | | | | | | | | |
| Interval Length | 0.01 *** | 0.01 – 0.02 | 5.08 | **<0.001** | -0.03 *** | -0.03 – -0.02 | -8.79 | **<0.001** | 0.88 *** | 0.85 – 0.91 | -7.70 | **<0.001** |
| Interval Session Number | -0.00 | -0.01 – 0.00 | -1.08 | 0.278 | -0.01 ** | -0.01 – -0.00 | -2.78 | **0.005** | 1.04 * | 1.00 – 1.07 | 2.24 | **0.025** |
| Reward:Penalty | 0.01 ** | 0.00 – 0.01 | 3.23 | **0.001** | -0.02 *** | -0.03 – -0.02 | -7.75 | **<0.001** | 0.93 *** | 0.90 – 0.96 | -4.70 | **<0.001** |
| Caudal dACC:Penalty | -0.00 | -0.03 – 0.02 | -0.27 | 0.787 | -0.01 | -0.03 – 0.02 | -0.39 | 0.696 | 0.95 | 0.83 – 1.09 | -0.71 | 0.480 |
| Caudal dACC:Reward | 0.01 | -0.01 – 0.04 | 1.00 | 0.318 | -0.04 ** | -0.06 – -0.01 | -2.97 | **0.003** | 0.91 | 0.80 – 1.05 | -1.26 | 0.207 |
| Congruency | | | | | -0.19 *** | -0.20 – -0.18 | -31.85 | **<0.001** | 1.14 *** | 1.07 – 1.21 | 3.89 | **<0.001** |
| **Random Effects** | | | | | | | | | | | | |
| $\sigma^2$ | 0.08 | | | | 0.76 | | | | 3.29 | | | |
| $\tau_{00}$ | 0.08 SubID | | | | 0.23 SubID | | | | 0.42 SubID | | | |
| N | 94 SubID | | | | 94 SubID | | | | 94 SubID | | | |
| Observations | 11551 | | | | 88901 | | | | 93150 | | | |
| Marginal $R^2$ / Conditional $R^2$ | 0.024 / 0.494 | | | | 0.026 / 0.254 | | | | 0.024 / 0.135 | | | |

* p<0.05 ** p<0.01 *** p<0.001

**Table S25:** *Task Performance Predicted by Rostral dACC during Cue and Interval Phase.*

**Task Performance Predicted by Rostral dACC (Cue)**

| | Response Rate | | | | Log RT | | | | Accuracy | | | |
|---|---|---|---|---|---|---|---|---|---|---|---|---|
| *Predictors* | *Estimates* | *CI* | *Statistic* | *p* | *Estimates* | *CI* | *Statistic* | *p* | *Odds Ratios* | *CI* | *Statistic* | *p* |
| (Intercept) | 1.50 *** | 1.44 – 1.56 | 52.41 | **<0.001** | 0.12 * | 0.02 – 0.22 | 2.39 | **0.017** | 24.74 *** | 21.53 – 28.42 | 45.35 | **<0.001** |
| Reward | 0.05 *** | 0.05 – 0.06 | 19.52 | **<0.001** | -0.11 *** | -0.11 – -0.10 | -36.40 | **<0.001** | 0.83 *** | 0.81 – 0.86 | -11.24 | **<0.001** |
| Penalty | -0.03 *** | -0.03 – -0.02 | -10.14 | **<0.001** | 0.08 *** | 0.07 – 0.08 | 24.18 | **<0.001** | 1.26 *** | 1.22 – 1.30 | 13.22 | **<0.001** |
| Rostral dACC | -0.00 | -0.00 – 0.00 | -0.06 | 0.956 | 0.00 | -0.00 – 0.01 | 1.41 | 0.160 | 1.04 ** | 1.01 – 1.06 | 2.79 | **0.005** |
| Interval Congruency | 0.02 *** | 0.01 – 0.02 | 6.34 | **<0.001** | | | | | | | | |
| Interval Length | 0.01 *** | 0.01 – 0.02 | 5.03 | **<0.001** | -0.03 *** | -0.03 – -0.02 | -8.67 | **<0.001** | 0.88 *** | 0.85 – 0.91 | -7.78 | **<0.001** |
| Interval Session Number | -0.00 | -0.01 – 0.00 | -1.26 | 0.209 | -0.01 * | -0.01 – -0.00 | -2.36 | **0.018** | 1.04 * | 1.01 – 1.07 | 2.32 | **0.020** |
| Reward:Penalty | 0.01 ** | 0.00 – 0.01 | 3.25 | **0.001** | -0.02 *** | -0.03 – -0.02 | -7.75 | **<0.001** | 0.93 *** | 0.90 – 0.96 | -4.68 | **<0.001** |
| Rostral dACC:Penalty | 0.00 | -0.00 – 0.00 | 0.13 | 0.897 | -0.00 | -0.00 – 0.00 | -0.15 | 0.885 | 1.02 | 1.00 – 1.04 | 1.58 | 0.114 |
| Rostral dACC:Reward | -0.00 | -0.01 – 0.00 | -1.89 | 0.059 | 0.00 | -0.00 – 0.01 | 0.96 | 0.335 | 0.98 | 0.95 – 1.00 | -1.94 | 0.053 |
| Congruency | | | | | -0.19 *** | -0.20 – -0.18 | -31.81 | **<0.001** | 1.14 *** | 1.06 – 1.21 | 3.89 | **<0.001** |
| **Random Effects** | | | | | | | | | | | | |
| $\sigma^2$ | 0.08 | | | | 0.76 | | | | 3.29 | | | |
| $\tau_{00}$ | $0.08_{\text{SubID}}$ | | | | $0.23_{\text{SubID}}$ | | | | $0.42_{\text{SubID}}$ | | | |
| N | $94_{\text{SubID}}$ | | | | $94_{\text{SubID}}$ | | | | $94_{\text{SubID}}$ | | | |
| Observations | 11551 | | | | 88901 | | | | 93150 | | | |
| Marginal $R^2$ / Conditional $R^2$ | 0.024 / 0.493 | | | | 0.026 / 0.254 | | | | 0.025 / 0.136 | | | |

* $p<0.05$ ** $p<0.01$ *** $p<0.001$

**Task Performance Predicted by Rostral dACC (Interval)**

| | Response Rate | | | | Log RT | | | | Accuracy | | | |
|---|---|---|---|---|---|---|---|---|---|---|---|---|
| *Predictors* | *Estimates* | *CI* | *Statistic* | *p* | *Estimates* | *CI* | *Statistic* | *p* | *Odds Ratios* | *CI* | *Statistic* | *p* |
| (Intercept) | 1.49 *** | 1.44 – 1.55 | 51.99 | **<0.001** | 0.13 ** | 0.04 – 0.23 | 2.69 | **0.007** | 25.08 *** | 21.80 – 28.87 | 44.94 | **<0.001** |
| Reward | 0.05 *** | 0.05 – 0.06 | 16.95 | **<0.001** | -0.11 *** | -0.11 – -0.10 | -31.10 | **<0.001** | 0.83 *** | 0.80 – 0.86 | -9.81 | **<0.001** |
| Penalty | -0.03 *** | -0.04 – -0.02 | -9.31 | **<0.001** | 0.08 *** | 0.07 – 0.08 | 21.18 | **<0.001** | 1.26 *** | 1.21 – 1.31 | 11.40 | **<0.001** |
| Rostral dACC | -0.06 *** | -0.09 – -0.04 | -5.28 | **<0.001** | 0.11 *** | 0.08 – 0.13 | 8.04 | **<0.001** | 1.12 | 0.97 – 1.29 | 1.59 | 0.111 |
| Interval Congruency | 0.02 *** | 0.01 – 0.02 | 6.39 | **<0.001** | | | | | | | | |
| Interval Length | 0.01 *** | 0.01 – 0.02 | 5.07 | **<0.001** | -0.03 *** | -0.03 – -0.02 | -8.70 | **<0.001** | 0.88 *** | 0.85 – 0.91 | -7.77 | **<0.001** |
| Interval Session Number | -0.00 | -0.01 – 0.00 | -1.41 | 0.160 | -0.01 * | -0.01 – -0.00 | -2.19 | **0.029** | 1.04 * | 1.00 – 1.07 | 2.24 | **0.025** |
| Reward:Penalty | 0.01 ** | 0.00 – 0.01 | 3.08 | **0.002** | -0.02 *** | -0.03 – -0.02 | -7.60 | **<0.001** | 0.93 *** | 0.90 – 0.96 | -4.69 | **<0.001** |
| Rostral dACC:Penalty | -0.01 | -0.03 – 0.01 | -0.99 | 0.321 | 0.01 | -0.01 – 0.03 | 0.82 | 0.410 | 0.99 | 0.87 – 1.13 | -0.16 | 0.875 |
| Rostral dACC:Reward | 0.00 | -0.02 – 0.02 | 0.05 | 0.961 | -0.01 | -0.03 – 0.02 | -0.60 | 0.550 | 0.97 | 0.85 – 1.10 | -0.51 | 0.613 |
| Congruency | | | | | -0.19 *** | -0.20 – -0.18 | -31.84 | **<0.001** | 1.14 *** | 1.07 – 1.21 | 3.89 | **<0.001** |
| **Random Effects** | | | | | | | | | | | | |
| $\sigma^2$ | 0.08 | | | | 0.76 | | | | 3.29 | | | |
| $\tau_{00}$ | $0.08_{\text{SubID}}$ | | | | $0.23_{\text{SubID}}$ | | | | $0.42_{\text{SubID}}$ | | | |
| N | $94_{\text{SubID}}$ | | | | $94_{\text{SubID}}$ | | | | $94_{\text{SubID}}$ | | | |
| Observations | 11551 | | | | 88901 | | | | 93150 | | | |
| Marginal $R^2$ / Conditional $R^2$ | 0.026 / 0.495 | | | | 0.026 / 0.255 | | | | 0.024 / 0.135 | | | |

* $p<0.05$ ** $p<0.01$ *** $p<0.001$

**Table S26:** *Task Performance Predicted by Dorsolateral PFC during Cue and Interval Phase.*

**Task Performance Predicted by Dorsolateral PFC (Cue)**

| | Response Rate | | | | Log RT | | | | Accuracy | | | |
|---|---|---|---|---|---|---|---|---|---|---|---|---|
| *Predictors* | *Estimates* | *CI* | *Statistic* | *p* | *Estimates* | *CI* | *Statistic* | *p* | *Odds Ratios* | *CI* | *Statistic* | *p* |
| (Intercept) | 1.50 *** | 1.45 – 1.56 | 52.51 | **<0.001** | 0.12 * | 0.02 – 0.22 | 2.37 | **0.018** | 24.65 *** | 21.46 – 28.32 | 45.33 | **<0.001** |
| Reward | 0.05 *** | 0.05 – 0.06 | 19.68 | **<0.001** | -0.11 *** | -0.11 – -0.10 | -36.50 | **<0.001** | 0.83 *** | 0.81 – 0.86 | -11.11 | **<0.001** |
| Penalty | -0.03 *** | -0.04 – -0.02 | -10.22 | **<0.001** | 0.08 *** | 0.07 – 0.08 | 24.34 | **<0.001** | 1.26 *** | 1.22 – 1.30 | 13.20 | **<0.001** |
| Dorsolateral PFC | 0.01 *** | 0.00 – 0.01 | 3.78 | **<0.001** | -0.01 *** | -0.01 – -0.01 | -4.31 | **<0.001** | 1.02 | 1.00 – 1.05 | 1.68 | 0.092 |
| Interval Congruency | 0.02 *** | 0.01 – 0.02 | 6.33 | **<0.001** | | | | | | | | |
| Interval Length | 0.01 *** | 0.01 – 0.02 | 5.01 | **<0.001** | -0.03 *** | -0.03 – -0.02 | -8.66 | **<0.001** | 0.88 *** | 0.85 – 0.91 | -7.78 | **<0.001** |
| Interval Session Number | -0.00 | -0.01 – 0.00 | -1.07 | 0.285 | -0.01 ** | -0.01 – -0.00 | -2.65 | **0.008** | 1.04 * | 1.01 – 1.07 | 2.28 | **0.023** |
| Reward:Penalty | 0.01 ** | 0.00 – 0.01 | 3.22 | **0.001** | -0.02 *** | -0.03 – -0.02 | -7.74 | **<0.001** | 0.93 *** | 0.90 – 0.96 | -4.68 | **<0.001** |
| Dorsolateral PFC:Penalty | -0.00 | -0.01 – 0.00 | -1.15 | 0.249 | 0.01 ** | 0.00 – 0.01 | 2.70 | **0.007** | 1.03 ** | 1.01 – 1.06 | 2.66 | **0.008** |
| Dorsolateral PFC:Reward | 0.00 * | 0.00 – 0.01 | 2.28 | **0.023** | -0.01 *** | -0.01 – -0.00 | -3.37 | **0.001** | 1.00 | 0.97 – 1.02 | -0.41 | 0.684 |
| Congruency | | | | | -0.19 *** | -0.20 – -0.18 | -31.79 | **<0.001** | 1.14 *** | 1.07 – 1.21 | 3.91 | **<0.001** |
| **Random Effects** | | | | | | | | | | | | |
| $\sigma^2$ | 0.08 | | | | 0.76 | | | | 3.29 | | | |
| $\tau_{00}$ | 0.08 $_{SubID}$ | | | | 0.23 $_{SubID}$ | | | | 0.42 $_{SubID}$ | | | |
| N | 94 $_{SubID}$ | | | | 94 $_{SubID}$ | | | | 94 $_{SubID}$ | | | |
| Observations | 11551 | | | | 88901 | | | | 93150 | | | |
| Marginal $R^2$ / Conditional $R^2$ | 0.025 / 0.493 | | | | 0.026 / 0.254 | | | | 0.025 / 0.135 | | | |

* *p<0.05* ** *p<0.01* *** *p<0.001*

**Task Performance Predicted by Dorsolateral PFC (Interval)**

| | Response Rate | | | | Log RT | | | | Accuracy | | | |
|---|---|---|---|---|---|---|---|---|---|---|---|---|
| *Predictors* | *Estimates* | *CI* | *Statistic* | *p* | *Estimates* | *CI* | *Statistic* | *p* | *Odds Ratios* | *CI* | *Statistic* | *p* |
| (Intercept) | 1.50 *** | 1.44 – 1.56 | 52.33 | **<0.001** | 0.12 * | 0.02 – 0.22 | 2.42 | **0.015** | 24.69 *** | 21.49 – 28.36 | 45.32 | **<0.001** |
| Reward | 0.05 *** | 0.05 – 0.06 | 19.63 | **<0.001** | -0.11 *** | -0.12 – -0.10 | -36.54 | **<0.001** | 0.83 *** | 0.80 – 0.86 | -11.20 | **<0.001** |
| Penalty | -0.03 *** | -0.04 – -0.02 | -10.17 | **<0.001** | 0.08 *** | 0.07 – 0.08 | 24.03 | **<0.001** | 1.26 *** | 1.22 – 1.31 | 13.09 | **<0.001** |
| Dorsolateral PFC | -0.04 ** | -0.06 – -0.01 | -2.96 | **0.003** | 0.05 *** | 0.02 – 0.07 | 3.35 | **0.001** | 1.04 | 0.90 – 1.20 | 0.52 | 0.602 |
| Interval Congruency | 0.02 *** | 0.01 – 0.02 | 6.36 | **<0.001** | | | | | | | | |
| Interval Length | 0.01 *** | 0.01 – 0.02 | 5.03 | **<0.001** | -0.03 *** | -0.03 – -0.02 | -8.66 | **<0.001** | 0.88 *** | 0.85 – 0.91 | -7.74 | **<0.001** |
| Interval Session Number | -0.00 | -0.01 – 0.00 | -1.52 | 0.128 | -0.01 * | -0.01 – -0.00 | -2.12 | **0.034** | 1.04 * | 1.00 – 1.07 | 2.21 | **0.027** |
| Reward:Penalty | 0.01 ** | 0.00 – 0.01 | 3.06 | **0.002** | -0.02 *** | -0.03 – -0.02 | -7.53 | **<0.001** | 0.93 *** | 0.90 – 0.96 | -4.70 | **<0.001** |
| Dorsolateral PFC:Penalty | -0.02 | -0.04 – 0.00 | -1.78 | 0.075 | 0.03 * | 0.00 – 0.05 | 2.26 | **0.024** | 1.07 | 0.94 – 1.23 | 1.03 | 0.303 |
| Dorsolateral PFC:Reward | 0.02 * | 0.00 – 0.04 | 1.96 | **0.050** | -0.05 *** | -0.08 – -0.03 | -4.26 | **<0.001** | 0.91 | 0.80 – 1.04 | -1.37 | 0.170 |
| Congruency | | | | | -0.19 *** | -0.20 – -0.18 | -31.82 | **<0.001** | 1.14 *** | 1.07 – 1.21 | 3.90 | **<0.001** |
| **Random Effects** | | | | | | | | | | | | |
| $\sigma^2$ | 0.08 | | | | 0.76 | | | | 3.29 | | | |
| $\tau_{00}$ | 0.08 $_{SubID}$ | | | | 0.23 $_{SubID}$ | | | | 0.42 $_{SubID}$ | | | |
| N | 94 $_{SubID}$ | | | | 94 $_{SubID}$ | | | | 94 $_{SubID}$ | | | |
| Observations | 11551 | | | | 88901 | | | | 93150 | | | |
| Marginal $R^2$ / Conditional $R^2$ | 0.025 / 0.494 | | | | 0.026 / 0.254 | | | | 0.024 / 0.135 | | | |

* *p<0.05* ** *p<0.01* *** *p<0.001*

**Table S27:** *Task Performance Predicted by Inferior Frontal Gyrus during Cue and Interval Phase.*

**Task Performance Predicted by Inferior Frontal Gyrus (Interval)**

| | Response Rate | | | | Log RT | | | | Accuracy | | | |
|---|---|---|---|---|---|---|---|---|---|---|---|---|
| *Predictors* | *Estimates* | *CI* | *Statistic* | *p* | *Estimates* | *CI* | *Statistic* | *p* | *Odds Ratios* | *CI* | *Statistic* | *p* |
| (Intercept) | 1.49 *** | 1.43 – 1.55 | 51.79 | **<0.001** | 0.14 ** | 0.04 – 0.24 | 2.75 | **0.006** | 25.58 *** | 22.22 – 29.45 | 45.12 | **<0.001** |
| Reward | 0.05 *** | 0.05 – 0.06 | 16.44 | **<0.001** | -0.11 *** | -0.12 – -0.10 | -29.76 | **<0.001** | 0.83 *** | 0.80 – 0.86 | -9.14 | **<0.001** |
| Penalty | -0.03 *** | -0.04 – -0.02 | -8.86 | **<0.001** | 0.08 *** | 0.07 – 0.08 | 20.27 | **<0.001** | 1.27 *** | 1.22 – 1.33 | 11.35 | **<0.001** |
| Inferior Frontal Gyrus (IFG) | -0.06 *** | -0.09 – -0.04 | -5.20 | **<0.001** | 0.11 *** | 0.09 – 0.14 | 8.24 | **<0.001** | 1.24 ** | 1.07 – 1.44 | 2.92 | **0.004** |
| Interval Congruency | 0.02 *** | 0.01 – 0.02 | 6.46 | **<0.001** | | | | | | | | |
| Interval Length | 0.01 *** | 0.01 – 0.02 | 5.04 | **<0.001** | -0.03 *** | -0.03 – -0.02 | -8.68 | **<0.001** | 0.88 *** | 0.85 – 0.91 | -7.79 | **<0.001** |
| Interval Session Number | -0.00 | -0.01 – 0.00 | -1.56 | 0.118 | -0.01 | -0.01 – 0.00 | -1.91 | 0.056 | 1.04 * | 1.01 – 1.07 | 2.33 | **0.020** |
| Reward:Penalty | 0.01 ** | 0.00 – 0.01 | 3.14 | **0.002** | -0.02 *** | -0.03 – -0.02 | -7.70 | **<0.001** | 0.93 *** | 0.90 – 0.96 | -4.68 | **<0.001** |
| IFG:Penalty | -0.01 | -0.03 – 0.01 | -0.94 | 0.347 | 0.01 | -0.01 – 0.04 | 0.91 | 0.362 | 1.07 | 0.94 – 1.23 | 1.02 | 0.310 |
| IFG:Reward | 0.01 | -0.01 – 0.03 | 0.77 | 0.440 | -0.01 | -0.04 – 0.01 | -0.86 | 0.392 | 0.97 | 0.85 – 1.11 | -0.42 | 0.678 |
| Congruency | | | | | -0.19 *** | -0.20 – -0.18 | -31.87 | **<0.001** | 1.13 *** | 1.06 – 1.21 | 3.88 | **<0.001** |
| **Random Effects** | | | | | | | | | | | | |
| $\sigma^2$ | 0.08 | | | | 0.76 | | | | 3.29 | | | |
| $\tau_{00}$ | 0.08 $_{SubID}$ | | | | 0.23 $_{SubID}$ | | | | 0.42 $_{SubID}$ | | | |
| N | 94 $_{SubID}$ | | | | 94 $_{SubID}$ | | | | 94 $_{SubID}$ | | | |
| Observations | 11551 | | | | 88901 | | | | 93150 | | | |
| Marginal $R^2$ / Conditional $R^2$ | 0.025 / 0.496 | | | | 0.026 / 0.256 | | | | 0.025 / 0.135 | | | |

* p<0.05 ** p<0.01 *** p<0.001


**Task Performance Predicted by Inferior Frontal Gyrus (Interval)**

| | Response Rate | | | | Log RT | | | | Accuracy | | | |
|---|---|---|---|---|---|---|---|---|---|---|---|---|
| *Predictors* | *Estimates* | *CI* | *Statistic* | *p* | *Estimates* | *CI* | *Statistic* | *p* | *Odds Ratios* | *CI* | *Statistic* | *p* |
| (Intercept) | 1.49 *** | 1.43 – 1.55 | 51.79 | **<0.001** | 0.14 ** | 0.04 – 0.24 | 2.75 | **0.006** | 25.58 *** | 22.22 – 29.45 | 45.12 | **<0.001** |
| Reward | 0.05 *** | 0.05 – 0.06 | 16.44 | **<0.001** | -0.11 *** | -0.12 – -0.10 | -29.76 | **<0.001** | 0.83 *** | 0.80 – 0.86 | -9.14 | **<0.001** |
| Penalty | -0.03 *** | -0.04 – -0.02 | -8.86 | **<0.001** | 0.08 *** | 0.07 – 0.08 | 20.27 | **<0.001** | 1.27 *** | 1.22 – 1.33 | 11.35 | **<0.001** |
| Inferior Frontal Gyrus (IFG) | -0.06 *** | -0.09 – -0.04 | -5.20 | **<0.001** | 0.11 *** | 0.09 – 0.14 | 8.24 | **<0.001** | 1.24 ** | 1.07 – 1.44 | 2.92 | **0.004** |
| Interval Congruency | 0.02 *** | 0.01 – 0.02 | 6.46 | **<0.001** | | | | | | | | |
| Interval Length | 0.01 *** | 0.01 – 0.02 | 5.04 | **<0.001** | -0.03 *** | -0.03 – -0.02 | -8.68 | **<0.001** | 0.88 *** | 0.85 – 0.91 | -7.79 | **<0.001** |
| Interval Session Number | -0.00 | -0.01 – 0.00 | -1.56 | 0.118 | -0.01 | -0.01 – 0.00 | -1.91 | 0.056 | 1.04 * | 1.01 – 1.07 | 2.33 | **0.020** |
| Reward:Penalty | 0.01 ** | 0.00 – 0.01 | 3.14 | **0.002** | -0.02 *** | -0.03 – -0.02 | -7.70 | **<0.001** | 0.93 *** | 0.90 – 0.96 | -4.68 | **<0.001** |
| IFG:Penalty | -0.01 | -0.03 – 0.01 | -0.94 | 0.347 | 0.01 | -0.01 – 0.04 | 0.91 | 0.362 | 1.07 | 0.94 – 1.23 | 1.02 | 0.310 |
| IFG:Reward | 0.01 | -0.01 – 0.03 | 0.77 | 0.440 | -0.01 | -0.04 – 0.01 | -0.86 | 0.392 | 0.97 | 0.85 – 1.11 | -0.42 | 0.678 |
| Congruency | | | | | -0.19 *** | -0.20 – -0.18 | -31.87 | **<0.001** | 1.13 *** | 1.06 – 1.21 | 3.88 | **<0.001** |
| **Random Effects** | | | | | | | | | | | | |
| $\sigma^2$ | 0.08 | | | | 0.76 | | | | 3.29 | | | |
| $\tau_{00}$ | 0.08 $_{SubID}$ | | | | 0.23 $_{SubID}$ | | | | 0.42 $_{SubID}$ | | | |
| N | 94 $_{SubID}$ | | | | 94 $_{SubID}$ | | | | 94 $_{SubID}$ | | | |
| Observations | 11551 | | | | 88901 | | | | 93150 | | | |
| Marginal $R^2$ / Conditional $R^2$ | 0.025 / 0.496 | | | | 0.026 / 0.256 | | | | 0.025 / 0.135 | | | |

* p<0.05 ** p<0.01 *** p<0.001

**Table S28:** *Affect and Motivation Ratings by Reward and Penalty Incentive Conditions.* At the end of each session, participants rate each incentive cues (pleasantness, arousal, motivation, effort, attention, and difficult) are averaged by incentive cue (high/low reward/punishment).

| Question | Reward | Penalty | N | Rating | sd | se | ci |
|---|---|---|---|---|---|---|---|
| Pleasantness | Low Reward | Low Penalty | 94 | 0.298 | 1.310 | 0.135 | 0.268 |
| Pleasantness | Low Reward | High Penalty | 94 | -2.862 | 2.191 | 0.226 | 0.449 |
| Pleasantness | High Reward | Low Penalty | 94 | 3.553 | 1.722 | 0.178 | 0.353 |
| Pleasantness | High Reward | High Penalty | 94 | 1.936 | 1.458 | 0.150 | 0.299 |
| Arousal | Low Reward | Low Penalty | 94 | -1.277 | 2.710 | 0.280 | 0.555 |
| Arousal | Low Reward | High Penalty | 94 | 1.883 | 2.512 | 0.259 | 0.515 |
| Arousal | High Reward | Low Penalty | 94 | 2.064 | 2.436 | 0.251 | 0.499 |
| Arousal | High Reward | High Penalty | 94 | 1.383 | 1.984 | 0.205 | 0.406 |
| Motivation | Low Reward | Low Penalty | 94 | -0.447 | 2.095 | 0.216 | 0.429 |
| Motivation | Low Reward | High Penalty | 94 | 2.021 | 2.216 | 0.229 | 0.454 |
| Motivation | High Reward | Low Penalty | 94 | 3.606 | 1.694 | 0.175 | 0.347 |
| Motivation | High Reward | High Penalty | 94 | 2.596 | 1.629 | 0.168 | 0.334 |
| Effort | Low Reward | Low Penalty | 94 | 0.287 | 1.914 | 0.197 | 0.392 |
| Effort | Low Reward | High Penalty | 94 | 2.649 | 1.897 | 0.196 | 0.389 |
| Effort | High Reward | Low Penalty | 94 | 2.947 | 1.380 | 0.142 | 0.283 |
| Effort | High Reward | High Penalty | 94 | 2.574 | 1.279 | 0.132 | 0.262 |
| Attention | Low Reward | Low Penalty | 94 | 0.457 | 1.954 | 0.202 | 0.400 |
| Attention | Low Reward | High Penalty | 94 | 3.394 | 1.794 | 0.185 | 0.367 |
| Attention | High Reward | Low Penalty | 94 | 2.862 | 1.350 | 0.139 | 0.277 |
| Attention | High Reward | High Penalty | 94 | 2.468 | 1.407 | 0.145 | 0.288 |
| Difficult | Low Reward | Low Penalty | 94 | -1.266 | 1.944 | 0.200 | 0.398 |
| Difficult | Low Reward | High Penalty | 94 | 0.777 | 2.096 | 0.216 | 0.429 |
| Difficult | High Reward | Low Penalty | 94 | -0.564 | 1.683 | 0.174 | 0.345 |
| Difficult | High Reward | High Penalty | 94 | 0.000 | 1.616 | 0.167 | 0.331 |
| Motivation (Averaged) | Low Reward | Low Penalty | 94 | 0.099 | 1.682 | 0.173 | 0.344 |
| Motivation (Averaged) | Low Reward | High Penalty | 94 | 2.688 | 1.503 | 0.155 | 0.308 |
| Motivation (Averaged) | High Reward | Low Penalty | 94 | 3.138 | 1.172 | 0.121 | 0.240 |
| Motivation (Averaged) | High Reward | High Penalty | 94 | 2.546 | 1.170 | 0.121 | 0.240 |

**Table S29:** *Linear Mixed Models of Affect and Motivation Ratings*. Linear mixed models were conducted to test whether pleasantness, arousal, motivation, effort, attention, and difficulty ratings were predicted by reward and penalty incentive conditions, as well as their interactions. As the motivation, effort, and attention ratings are highly correlated and contain a similar interactive pattern and were combined into a 'Motivation (Averaged)' rating.

**Affect and Motivation Ratings Predicted by Incentive Conditions**

| | Pleasantness | | | | Arousal | | | | Difficulty | | | |
|---|---|---|---|---|---|---|---|---|---|---|---|---|
| *Predictors* | *Estimates* | *CI* | *Statistic* | *p* | *Estimates* | *CI* | *Statistic* | *p* | *Estimates* | *CI* | *Statistic* | *p* |
| (Intercept) | -0.01 | -0.07 – 0.06 | -0.17 | 0.864 | 0.02 | -0.07 – 0.11 | 0.38 | 0.707 | -0.01 | -0.14 – 0.13 | -0.08 | 0.938 |
| Reward | 0.68 *** | 0.63 – 0.74 | 22.92 | **<0.001** | 0.25 *** | 0.16 – 0.34 | 5.68 | **<0.001** | -0.01 | -0.09 – 0.07 | -0.20 | 0.845 |
| Penalty | -0.41 *** | -0.46 – -0.35 | -13.59 | **<0.001** | 0.22 *** | 0.13 – 0.31 | 4.95 | **<0.001** | 0.27 *** | 0.19 – 0.35 | 6.85 | **<0.001** |
| Reward:Penalty | 0.13 *** | 0.07 – 0.19 | 4.39 | **<0.001** | -0.34 *** | -0.43 – -0.25 | -7.68 | **<0.001** | -0.15 *** | -0.23 – -0.08 | -3.89 | **<0.001** |
| **Random Effects** | | | | | | | | | | | | |
| $\sigma^2$ | 0.34 | | | | 0.74 | | | | 0.59 | | | |
| $\tau_{00}$ | 0.01 $_{SubID}$ | | | | 0.02 $_{SubID}$ | | | | 0.33 $_{SubID}$ | | | |
| N | 94 $_{SubID}$ | | | | 94 $_{SubID}$ | | | | 94 $_{SubID}$ | | | |
| Observations | 376 | | | | 376 | | | | 376 | | | |
| Marginal $R^2$ / Conditional $R^2$ | 0.653 / 0.664 | | | | 0.231 / 0.253 | | | | 0.096 / 0.419 | | | |

* p<0.05 ** p<0.01 *** p<0.001

**Affect and Motivation Ratings Predicted by Incentive Conditions**

| | Motivation | | | | Effort | | | | Attention | | | | Motivation (Averaged) | | | |
|---|---|---|---|---|---|---|---|---|---|---|---|---|---|---|---|---|
| *Predictors* | *Estimates* | *CI* | *Statistic* | *p* | *Estimates* | *CI* | *Statistic* | *p* | *Estimates* | *CI* | *Statistic* | *p* | *Estimates* | *CI* | *Statistic* | *p* |
| (Intercept) | 0.00 | -0.11 – 0.12 | 0.04 | 0.966 | -0.00 | -0.14 – 0.13 | -0.07 | 0.947 | 0.00 | -0.13 – 0.13 | 0.06 | 0.955 | 0.00 | -0.13 – 0.13 | 0.01 | 0.991 |
| Reward | 0.43 *** | 0.36 – 0.50 | 11.65 | **<0.001** | 0.27 *** | 0.20 – 0.34 | 7.63 | **<0.001** | 0.16 *** | 0.09 – 0.23 | 4.36 | **<0.001** | 0.33 *** | 0.27 – 0.40 | 10.04 | **<0.001** |
| Penalty | 0.13 *** | 0.06 – 0.21 | 3.67 | **<0.001** | 0.21 *** | 0.14 – 0.28 | 5.87 | **<0.001** | 0.27 *** | 0.20 – 0.34 | 7.49 | **<0.001** | 0.23 *** | 0.16 – 0.29 | 6.92 | **<0.001** |
| Reward:Penalty | -0.32 *** | -0.39 – -0.25 | -8.76 | **<0.001** | -0.29 *** | -0.36 – -0.22 | -8.06 | **<0.001** | -0.35 *** | -0.42 – -0.28 | -9.81 | **<0.001** | -0.36 *** | -0.43 – -0.30 | -11.02 | **<0.001** |
| **Random Effects** | | | | | | | | | | | | | | | | |
| $\sigma^2$ | 0.51 | | | | 0.48 | | | | 0.49 | | | | 0.41 | | | |
| $\tau_{00}$ | 0.19 $_{SubID}$ | | | | 0.31 $_{SubID}$ | | | | 0.29 $_{SubID}$ | | | | 0.29 $_{SubID}$ | | | |
| N | 94 $_{SubID}$ | | | | 94 $_{SubID}$ | | | | 94 $_{SubID}$ | | | | 94 $_{SubID}$ | | | |
| Observations | 376 | | | | 376 | | | | 376 | | | | 376 | | | |
| Marginal $R^2$ / Conditional $R^2$ | 0.306 / 0.492 | | | | 0.205 / 0.513 | | | | 0.224 / 0.510 | | | | 0.296 / 0.589 | | | |

* p<0.05 ** p<0.01 *** p<0.001

**Table S30:** *Positive and Negative Arousal Values for Affective Circumplex.* Positive and Negative Arousal values were computed by dividing arousal and pleasantness (valence) ratings by √2, and either adding or subtracting for positive and negative valence. Positive Arousal (PA) = arousal√2 + valence√2, Negative Arousal (NA) = arousal√2 - valence√2 (See Knutson et al., 2005 for more details). We also computed the hypotenuse of the arousal and valence ratings in Figure 5 and confirmed they were within the expected range of positive and negative arousal values.

| Reward | Penalty | N | Pleasantness | Arousal | Hypotenuse | Pleasantness_mc | Arousal_mc | PosArousal | NegArousal |
|---|---|---|---|---|---|---|---|---|---|
| Low Reward | Low Penalty | 94 | 0.298 | -1.277 | -1.311 | -0.434 | -2.290 | -1.926 | -1.313 |
| Low Reward | High Penalty | 94 | -2.862 | 1.883 | 3.426 | -3.593 | 0.870 | -1.926 | 3.156 |
| High Reward | Low Penalty | 94 | 3.553 | 2.064 | 4.109 | 2.822 | 1.051 | 2.738 | -1.252 |
| High Reward | High Penalty | 94 | 1.936 | 1.383 | 2.379 | 1.205 | 0.370 | 1.113 | -0.591 |

**Table S31:** *Positive and Negative Arousal Predicted by Reward and Penalty.* Linear mixed models were conducted to test whether pleasantness, arousal, motivation, effort, attention, and difficulty ratings were predicted by reward and penalty incentive conditions, as well as their interactions. As the motivation, effort, and attention ratings are highly correlated and contain a similar interactive pattern and were combined into a 'Motivation (Averaged)' rating.

**Postive and Negative Arousal Predicted by Incentive Conditions**

| | **Positive Arousal** | | | | **Negative Arousal** | | | |
|---|---|---|---|---|---|---|---|---|
| *Predictors* | *Estimates* | *CI* | *Statistic* | *p* | *Estimates* | *CI* | *Statistic* | *p* |
| (Intercept) | -0.65 *** | -0.80 – -0.50 | -8.64 | **<0.001** | -0.47 *** | -0.62 – -0.32 | -6.08 | **<0.001** |
| Reward | 1.58 *** | 1.39 – 1.78 | 16.12 | **<0.001** | 0.02 | -0.19 – 0.24 | 0.20 | 0.844 |
| Penalty | -0.00 | -0.19 – 0.19 | -0.00 | 1.000 | 1.60 *** | 1.39 – 1.82 | 14.64 | **<0.001** |
| Reward:Penalty | -0.55 *** | -0.82 – -0.28 | -3.97 | **<0.001** | -1.37 *** | -1.67 – -1.06 | -8.82 | **<0.001** |
| **Random Effects** | | | | | | | | |
| $\sigma^2$ | 0.45 | | | | 0.56 | | | |
| $\tau_{00}$ | 0.08 $_{SubID}$ | | | | 0.00 $_{SubID}$ | | | |
| N | 94 $_{SubID}$ | | | | 94 $_{SubID}$ | | | |
| Observations | 376 | | | | 376 | | | |
| Marginal $R^2$ / Conditional $R^2$ | 0.465 / 0.549 | | | | 0.438 / NA | | | |

** p<0.05   ** p<0.01   *** p<0.001*

**Table S32:** P-values of correlation matrix of Self-Report Ratings and Model Parameters (Figure S9). The table includes subjective ratings (e.g., motivation, difficulty, positive arousal, negative rousal) vs. drift diffusion model parameters by incentive condition (v = drift, a = threshold). Ratings followed by _Avg reflect the average rating, ratings followed by _Rew reflect the difference between high vs. low reward, and ratings followed by _Pen reflect the difference between high vs low penalty. DDM parameters followed by _Reward reflects the difference between high vs. low reward, parameters followed by _Penalty reflects the difference between high vs. low penalty, and parameters followed by _Intercept reflect the intercept estimates. Incentive-related adjustments were most associated with motivation differences in high vs. low reward cues. P-values are Benjamini-Hochberg corrected.

P-Values of Correlation Matrix (FDR Corrected)

| | Mot_Avg | Mot_Rew | Mot_Pen | Diff_Avg | Diff_Rew | Diff_Pen | PA_Rew | PA_Pen | NA_Rew | NA_Pen |
|---|---|---|---|---|---|---|---|---|---|---|
| v_Reward | 0.416 | 0.000 | 0.671 | 0.275 | 0.009 | 0.671 | 0.416 | 0.909 | 0.416 | 0.875 |
| a_Reward | 0.214 | 0.000 | 0.909 | 0.435 | 0.134 | 0.909 | 0.618 | 0.655 | 0.951 | 0.628 |
| v_Penalty | 0.275 | 0.009 | 0.875 | 0.671 | 0.081 | 0.593 | 0.909 | 0.671 | 0.909 | 0.416 |
| a_Penalty | 0.416 | 0.014 | 0.909 | 0.593 | 0.606 | 0.432 | 0.909 | 0.416 | 0.909 | 0.960 |
| v_Intercept | 0.432 | 0.909 | 0.081 | 0.416 | 0.593 | 0.914 | 0.432 | 0.797 | 0.956 | 0.565 |
| a_Intercept | 0.593 | 0.432 | 0.009 | 0.432 | 0.914 | 0.275 | 0.099 | 0.441 | 0.930 | 0.593 |

**Table S33:** *Associations between Positive and Negative Ratings and Neural Activity During Cue Phase.* Linear mixed effects models with ROI activity during the cue phase revealed no associations between positive and negative arousal and neural activity and included experimental conditions such as interval length and interval session number to control for fatigue effects across the session.

**Cue Related ROI Activity by Positive and Negative Arousal Ratings**

| | Ventral Striatum | | | | Anterior Insula | | | | Caudal dACC | | | |
|---|---|---|---|---|---|---|---|---|---|---|---|---|
| *Predictors* | *Estimates* | *CI* | *Statistic* | *p* | *Estimates* | *CI* | *Statistic* | *p* | *Estimates* | *CI* | *Statistic* | *p* |
| (Intercept) | 0.08 ** | 0.02 – 0.14 | 2.70 | **0.007** | -0.14 *** | -0.20 – -0.08 | -4.37 | **<0.001** | 0.08 * | 0.02 – 0.14 | 2.45 | **0.014** |
| Positive Arousal | 0.03 * | 0.00 – 0.05 | 2.06 | **0.040** | -0.01 | -0.04 – 0.02 | -0.63 | 0.530 | 0.03 * | 0.00 – 0.06 | 2.08 | **0.038** |
| Negative Arousal | 0.04 ** | 0.01 – 0.07 | 2.93 | **0.003** | 0.02 | -0.01 – 0.05 | 1.45 | 0.147 | 0.02 | -0.01 – 0.05 | 1.52 | 0.128 |
| Interval Length | -0.00 | -0.02 – 0.02 | -0.12 | 0.907 | 0.01 | -0.02 – 0.03 | 0.60 | 0.548 | 0.02 | -0.01 – 0.04 | 1.49 | 0.135 |
| Interval Session Number | 0.01 | -0.01 – 0.03 | 1.16 | 0.244 | -0.04 *** | -0.07 – -0.02 | -3.75 | **<0.001** | -0.00 | -0.02 – 0.02 | -0.20 | 0.838 |
| Positive Arousal x Reward | -0.03 | -0.06 – 0.01 | -1.58 | 0.115 | 0.00 | -0.04 – 0.04 | 0.00 | 0.996 | -0.01 | -0.05 – 0.03 | -0.52 | 0.602 |
| Positive Arousal x Penalty | 0.00 | -0.02 – 0.03 | 0.08 | 0.940 | -0.01 | -0.04 – 0.02 | -0.52 | 0.602 | -0.01 | -0.03 – 0.02 | -0.38 | 0.702 |
| Negative Arousal x Reward | 0.01 | -0.03 – 0.04 | 0.45 | 0.651 | -0.00 | -0.04 – 0.04 | -0.06 | 0.953 | 0.01 | -0.03 – 0.04 | 0.31 | 0.758 |
| Negative Arousal x Penalty | -0.00 | -0.03 – 0.03 | -0.18 | 0.855 | -0.00 | -0.04 – 0.03 | -0.19 | 0.852 | 0.00 | -0.03 – 0.04 | 0.21 | 0.834 |
| **Random Effects** | | | | | | | | | | | | |
| $\sigma^2$ | 1.31 | | | | 1.56 | | | | 1.46 | | | |
| $\tau_{00}$ | 0.05 $_{\text{SubID}}$ | | | | 0.06 $_{\text{SubID}}$ | | | | 0.06 $_{\text{SubID}}$ | | | |
| N | 94 $_{\text{SubID}}$ | | | | 94 $_{\text{SubID}}$ | | | | 94 $_{\text{SubID}}$ | | | |
| Observations | 11551 | | | | 11551 | | | | 11551 | | | |
| Marginal $R^2$ / Conditional $R^2$ | 0.002 / 0.041 | | | | 0.002 / 0.038 | | | | 0.001 / 0.039 | | | |

** p<0.05 ** p<0.01 *** p<0.001*

**Cue Related ROI Activity by Positive and Negative Arousal Ratings**

| | Rostral dACC | | | | Dorsolateral PFC | | | | Inferior Frontal Gyrus | | | |
|---|---|---|---|---|---|---|---|---|---|---|---|---|
| *Predictors* | *Estimates* | *CI* | *Statistic* | *p* | *Estimates* | *CI* | *Statistic* | *p* | *Estimates* | *CI* | *Statistic* | *p* |
| (Intercept) | -0.06 * | -0.12 – -0.00 | -2.10 | **0.036** | -0.03 | -0.10 – 0.04 | -0.90 | 0.368 | -0.10 *** | -0.16 – -0.04 | -3.36 | **0.001** |
| Positive Arousal | -0.01 | -0.04 – 0.02 | -0.74 | 0.459 | 0.01 | -0.01 – 0.04 | 1.01 | 0.312 | -0.00 | -0.03 – 0.02 | -0.27 | 0.786 |
| Negative Arousal | 0.02 | -0.01 – 0.05 | 1.08 | 0.278 | 0.02 | -0.01 – 0.05 | 1.60 | 0.109 | 0.02 | -0.01 – 0.05 | 1.20 | 0.229 |
| Interval Length | 0.01 | -0.01 – 0.03 | 0.72 | 0.469 | 0.01 | -0.01 – 0.04 | 1.09 | 0.276 | 0.00 | -0.02 – 0.03 | 0.39 | 0.697 |
| Interval Session Number | -0.06 *** | -0.08 – -0.03 | -4.81 | **<0.001** | -0.04 *** | -0.06 – -0.02 | -3.58 | **<0.001** | -0.05 *** | -0.07 – -0.03 | -4.13 | **<0.001** |
| Positive Arousal x Reward | -0.00 | -0.04 – 0.03 | -0.19 | 0.846 | -0.00 | -0.04 – 0.04 | -0.01 | 0.990 | -0.01 | -0.04 – 0.03 | -0.31 | 0.755 |
| Positive Arousal x Penalty | 0.01 | -0.02 – 0.03 | 0.48 | 0.630 | 0.00 | -0.02 – 0.03 | 0.34 | 0.731 | 0.01 | -0.02 – 0.04 | 0.57 | 0.570 |
| Negative Arousal x Reward | -0.01 | -0.05 – 0.03 | -0.47 | 0.636 | -0.01 | -0.05 – 0.03 | -0.49 | 0.624 | -0.02 | -0.05 – 0.02 | -0.96 | 0.335 |
| Negative Arousal x Penalty | -0.01 | -0.04 – 0.03 | -0.52 | 0.602 | 0.02 | -0.02 – 0.05 | 0.89 | 0.375 | -0.02 | -0.05 – 0.02 | -0.86 | 0.390 |
| **Random Effects** | | | | | | | | | | | | |
| $\sigma^2$ | 1.55 | | | | 1.58 | | | | 1.55 | | | |
| $\tau_{00}$ | 0.05 $_{\text{SubID}}$ | | | | 0.08 $_{\text{SubID}}$ | | | | 0.05 $_{\text{SubID}}$ | | | |
| N | 94 $_{\text{SubID}}$ | | | | 94 $_{\text{SubID}}$ | | | | 94 $_{\text{SubID}}$ | | | |
| Observations | 11551 | | | | 11551 | | | | 11551 | | | |
| Marginal $R^2$ / Conditional $R^2$ | 0.002 / 0.036 | | | | 0.002 / 0.048 | | | | 0.002 / 0.035 | | | |

** p<0.05 ** p<0.01 *** p<0.001*

**Table S34:** *Associations between Positive and Negative Ratings and Neural Activity During Interval Phase*. Linear mixed effects models with ROI activity during the cue phase revealed no associations between positive and negative arousal and neural activity and included experimental conditions such as interval length and interval session number to control for fatigue effects across the session.

**Interval Related ROI Activity by Positive and Negative Arousal Ratings**

| | Ventral Striatum | | | | Anterior Insula | | | | Caudal dACC | | | |
|---|---|---|---|---|---|---|---|---|---|---|---|---|
| *Predictors* | *Estimates* | *CI* | *Statistic* | *p* | *Estimates* | *CI* | *Statistic* | *p* | *Estimates* | *CI* | *Statistic* | *p* |
| (Intercept) | 0.19 *** | 0.17 – 0.20 | 20.70 | **<0.001** | -0.13 *** | -0.14 – -0.11 | -14.53 | **<0.001** | 0.23 *** | 0.22 – 0.25 | 24.39 | **<0.001** |
| Positive Arousal | 0.00 * | 0.00 – 0.01 | 2.05 | **0.041** | -0.00 | -0.01 – 0.00 | -0.89 | 0.373 | 0.00 | -0.00 – 0.01 | 1.62 | 0.105 |
| Negative Arousal | 0.01 * | 0.00 – 0.01 | 2.44 | **0.015** | 0.01 *** | 0.01 – 0.02 | 3.79 | **<0.001** | 0.00 | -0.00 – 0.01 | 1.88 | 0.060 |
| Interval Length | -0.00 | -0.00 – 0.00 | -0.57 | 0.571 | -0.00 | -0.01 – 0.00 | -0.62 | 0.536 | -0.01 ** | -0.01 – -0.00 | -3.22 | **0.001** |
| Interval Session Number | 0.00 ** | 0.00 – 0.01 | 2.59 | **0.010** | -0.02 *** | -0.03 – -0.02 | -10.21 | **<0.001** | -0.02 *** | -0.02 – -0.01 | -8.76 | **<0.001** |
| Positive Arousal x Reward | 0.01 | -0.00 – 0.01 | 1.63 | 0.104 | 0.00 | -0.00 – 0.01 | 1.34 | 0.181 | 0.01 * | 0.00 – 0.01 | 2.20 | **0.028** |
| Positive Arousal x Penalty | 0.01 ** | 0.00 – 0.01 | 3.11 | **0.002** | 0.00 | -0.00 – 0.01 | 1.56 | 0.118 | 0.00 | -0.00 – 0.01 | 1.08 | 0.282 |
| Negative Arousal x Reward | -0.00 | -0.01 – 0.00 | -1.20 | 0.232 | -0.01 | -0.01 – 0.00 | -1.86 | 0.063 | -0.00 | -0.01 – 0.00 | -1.04 | 0.297 |
| Negative Arousal x Penalty | -0.01 ** | -0.01 – -0.00 | -2.82 | **0.005** | -0.01 * | -0.01 – -0.00 | -2.42 | **0.016** | -0.01 ** | -0.02 – -0.00 | -3.07 | **0.002** |
| **Random Effects** | | | | | | | | | | | | |
| $\sigma^2$ | 0.04 | | | | 0.05 | | | | 0.04 | | | |
| $\tau_{00}$ | 0.01 SubID | | | | 0.01 SubID | | | | 0.01 SubID | | | |
| N | 94 SubID | | | | 94 SubID | | | | 94 SubID | | | |
| Observations | 11551 | | | | 11551 | | | | 11551 | | | |
| Marginal $R^2$ / Conditional $R^2$ | 0.003 / 0.141 | | | | 0.010 / 0.116 | | | | 0.008 / 0.156 | | | |

** p<0.05 ** p<0.01 *** p<0.001*

**Interval Related ROI Activity by Positive and Negative Arousal Ratings**

| | Rostral dACC | | | | Dorsolateral PFC | | | | Inferior Frontal Gyrus | | | |
|---|---|---|---|---|---|---|---|---|---|---|---|---|
| *Predictors* | *Estimates* | *CI* | *Statistic* | *p* | *Estimates* | *CI* | *Statistic* | *p* | *Estimates* | *CI* | *Statistic* | *p* |
| (Intercept) | -0.14 *** | -0.16 – -0.12 | -14.45 | **<0.001** | -0.05 *** | -0.06 – -0.03 | -4.91 | **<0.001** | -0.16 *** | -0.18 – -0.14 | -18.70 | **<0.001** |
| Positive Arousal | -0.01 ** | -0.01 – -0.00 | -2.65 | **0.008** | -0.00 | -0.01 – 0.00 | -0.76 | 0.449 | -0.01 ** | -0.01 – -0.00 | -2.82 | **0.005** |
| Negative Arousal | 0.01 *** | 0.01 – 0.02 | 4.31 | **<0.001** | 0.01 ** | 0.00 – 0.01 | 3.20 | **0.001** | 0.01 *** | 0.01 – 0.02 | 4.28 | **<0.001** |
| Interval Length | 0.00 | -0.00 – 0.01 | 0.42 | 0.674 | 0.00 | -0.00 – 0.00 | 0.16 | 0.877 | 0.00 | -0.00 – 0.01 | 0.82 | 0.415 |
| Interval Session Number | -0.01 ** | -0.01 – -0.00 | -3.03 | **0.002** | -0.02 *** | -0.03 – -0.02 | -10.66 | **<0.001** | -0.01 *** | -0.02 – -0.01 | -6.26 | **<0.001** |
| Positive Arousal x Reward | 0.01 ** | 0.00 – 0.02 | 2.58 | **0.010** | 0.01 | -0.00 – 0.01 | 1.73 | 0.084 | 0.01 * | 0.00 – 0.01 | 1.96 | **0.050** |
| Positive Arousal x Penalty | 0.01 * | 0.00 – 0.01 | 2.02 | **0.044** | 0.00 | -0.00 – 0.01 | 0.73 | 0.466 | 0.01 ** | 0.00 – 0.01 | 3.26 | **0.001** |
| Negative Arousal x Reward | -0.01 | -0.01 – 0.00 | -1.51 | 0.130 | -0.01 | -0.01 – 0.00 | -1.53 | 0.126 | -0.01 * | -0.01 – -0.00 | -2.05 | **0.041** |
| Negative Arousal x Penalty | -0.01 ** | -0.01 – -0.00 | -2.59 | **0.010** | -0.01 * | -0.01 – -0.00 | -1.97 | **0.048** | -0.01 * | -0.01 – -0.00 | -2.18 | **0.030** |
| **Random Effects** | | | | | | | | | | | | |
| $\sigma^2$ | 0.05 | | | | 0.05 | | | | 0.05 | | | |
| $\tau_{00}$ | 0.01 SubID | | | | 0.01 SubID | | | | 0.01 SubID | | | |
| N | 94 SubID | | | | 94 SubID | | | | 94 SubID | | | |
| Observations | 11551 | | | | 11551 | | | | 11551 | | | |
| Marginal $R^2$ / Conditional $R^2$ | 0.004 / 0.133 | | | | 0.011 / 0.133 | | | | 0.008 / 0.117 | | | |

** p<0.05 ** p<0.01 *** p<0.001*

**Table S35:** *Motivation Ratings Predict Neural Activity During Cue Phase*. Linear mixed effects models with ROI activity during the cue phase predicted by self-report ratings of mean motivation (average across motivation, effort, and attention) and included experimental conditions such as interval length and interval session number to control for fatigue effects across the session.

**Cue Related ROI Activity by Mean Motivation Rating**

| | Ventral Striatum | | | | Anterior Insula | | | | Caudal dACC | | | |
|---|---|---|---|---|---|---|---|---|---|---|---|---|
| *Predictors* | *Estimates* | *CI* | *Statistic* | *p* | *Estimates* | *CI* | *Statistic* | *p* | *Estimates* | *CI* | *Statistic* | *p* |
| (Intercept) | 0.06 * | 0.01 – 0.11 | 2.18 | **0.030** | -0.13 *** | -0.19 – -0.08 | -4.78 | **<0.001** | 0.07 * | 0.01 – 0.12 | 2.37 | **0.018** |
| Mean Motivation | 0.04 * | 0.01 – 0.07 | 2.56 | **0.010** | 0.00 | -0.03 – 0.04 | 0.22 | 0.824 | 0.04 * | 0.01 – 0.07 | 2.38 | **0.017** |
| Interval Length | 0.00 | -0.02 – 0.02 | 0.09 | 0.925 | 0.01 | -0.01 – 0.03 | 0.88 | 0.380 | 0.02 | -0.00 – 0.04 | 1.53 | 0.126 |
| Interval Session Number | 0.01 | -0.01 – 0.03 | 1.18 | 0.238 | -0.04 *** | -0.07 – -0.02 | -3.73 | **<0.001** | -0.00 | -0.02 – 0.02 | -0.20 | 0.840 |
| Mean Motivation:Reward | -0.00 | -0.03 – 0.03 | -0.11 | 0.912 | -0.00 | -0.03 – 0.02 | -0.31 | 0.755 | 0.01 | -0.02 – 0.04 | 0.73 | 0.468 |
| Mean Motivation:Penalty | 0.00 | -0.02 – 0.03 | 0.10 | 0.917 | -0.00 | -0.03 – 0.02 | -0.34 | 0.731 | 0.00 | -0.02 – 0.03 | 0.10 | 0.921 |
| **Random Effects** | | | | | | | | | | | | |
| $\sigma^2$ | 1.31 | | | | 1.56 | | | | 1.46 | | | |
| $\tau_{00}$ | 0.05 $_{SubID}$ | | | | 0.06 $_{SubID}$ | | | | 0.06 $_{SubID}$ | | | |
| N | 94 $_{SubID}$ | | | | 94 $_{SubID}$ | | | | 94 $_{SubID}$ | | | |
| Observations | 11551 | | | | 11551 | | | | 11551 | | | |
| Marginal $R^2$ / Conditional $R^2$ | 0.001 / 0.041 | | | | 0.001 / 0.038 | | | | 0.001 / 0.039 | | | |

* p<0.05 ** p<0.01 *** p<0.001

**Cue Related ROI Activity by Mean Motivation Rating**

| | Rostral dACC | | | | Dorsolateral PFC | | | | Inferior Frontal Gyrus | | | |
|---|---|---|---|---|---|---|---|---|---|---|---|---|
| *Predictors* | *Estimates* | *CI* | *Statistic* | *p* | *Estimates* | *CI* | *Statistic* | *p* | *Estimates* | *CI* | *Statistic* | *p* |
| (Intercept) | -0.07 * | -0.12 – -0.02 | -2.55 | **0.011** | -0.02 | -0.09 – 0.04 | -0.79 | 0.428 | -0.11 *** | -0.16 – -0.05 | -3.94 | **<0.001** |
| Mean Motivation | 0.01 | -0.03 – 0.04 | 0.47 | 0.637 | 0.02 | -0.01 – 0.06 | 1.40 | 0.163 | 0.00 | -0.03 – 0.04 | 0.10 | 0.920 |
| Interval Length | 0.01 | -0.01 – 0.03 | 1.03 | 0.305 | 0.02 | -0.01 – 0.04 | 1.37 | 0.171 | 0.01 | -0.01 – 0.03 | 0.86 | 0.387 |
| Interval Session Number | -0.06 *** | -0.08 – -0.03 | -4.80 | **<0.001** | -0.04 *** | -0.06 – -0.02 | -3.58 | **<0.001** | -0.05 *** | -0.07 – -0.03 | -4.13 | **<0.001** |
| Mean Motivation:Reward | -0.00 | -0.03 – 0.03 | -0.13 | 0.893 | 0.01 | -0.02 – 0.04 | 0.40 | 0.686 | 0.00 | -0.03 – 0.03 | 0.02 | 0.984 |
| Mean Motivation:Penalty | 0.01 | -0.02 – 0.03 | 0.50 | 0.616 | 0.01 | -0.02 – 0.04 | 0.63 | 0.531 | -0.00 | -0.03 – 0.02 | -0.25 | 0.800 |
| **Random Effects** | | | | | | | | | | | | |
| $\sigma^2$ | 1.55 | | | | 1.58 | | | | 1.55 | | | |
| $\tau_{00}$ | 0.05 $_{SubID}$ | | | | 0.08 $_{SubID}$ | | | | 0.05 $_{SubID}$ | | | |
| N | 94 $_{SubID}$ | | | | 94 $_{SubID}$ | | | | 94 $_{SubID}$ | | | |
| Observations | 11551 | | | | 11551 | | | | 11551 | | | |
| Marginal $R^2$ / Conditional $R^2$ | 0.002 / 0.035 | | | | 0.002 / 0.047 | | | | 0.002 / 0.035 | | | |

* p<0.05 ** p<0.01 *** p<0.001

**Table S36:** *Motivation Ratings Predict Neural Activity During Interval Phase*. Linear mixed effects models with ROI activity during the interval phase predicted by self-report ratings of mean motivation (average across motivation, effort, and attention) and included experimental conditions such as interval length and interval session number to control for fatigue effects across the session.

**Interval Related ROI Activity by Mean Motivation Rating**

| | **Ventral Striatum** | | | | **Anterior Insula** | | | | **Caudal dACC** | | | |
|---|---|---|---|---|---|---|---|---|---|---|---|---|
| *Predictors* | *Estimates* | *CI* | *Statistic* | *p* | *Estimates* | *CI* | *Statistic* | *p* | *Estimates* | *CI* | *Statistic* | *p* |
| (Intercept) | 0.18 *** | 0.17 – 0.20 | 21.01 | **<0.001** | -0.13 *** | -0.14 – -0.11 | -15.13 | **<0.001** | 0.23 *** | 0.22 – 0.25 | 25.11 | **<0.001** |
| Mean Motivation | 0.01 | -0.00 – 0.01 | 1.94 | 0.052 | 0.00 | -0.01 – 0.01 | 0.15 | 0.878 | 0.00 | -0.00 – 0.01 | 0.82 | 0.411 |
| Interval Length | 0.00 | -0.00 – 0.00 | 0.10 | 0.922 | 0.00 | -0.00 – 0.01 | 0.50 | 0.614 | -0.01 ** | -0.01 – -0.00 | -2.61 | **0.009** |
| Interval Session Number | 0.00 * | 0.00 – 0.01 | 2.52 | **0.012** | -0.02 *** | -0.03 – -0.02 | -10.22 | **<0.001** | -0.02 *** | -0.02 – -0.01 | -8.81 | **<0.001** |
| Mean Motivation:Reward | 0.01 * | 0.00 – 0.01 | 2.39 | **0.017** | 0.00 | -0.00 – 0.01 | 0.25 | 0.804 | 0.01 * | 0.00 – 0.01 | 2.40 | **0.017** |
| Mean Motivation:Penalty | 0.00 | -0.00 – 0.01 | 0.88 | 0.381 | 0.00 | -0.00 – 0.01 | 0.27 | 0.784 | -0.00 | -0.01 – 0.00 | -1.36 | 0.175 |
| **Random Effects** | | | | | | | | | | | | |
| $\sigma^2$ | 0.04 | | | | 0.05 | | | | 0.04 | | | |
| $\tau_{00}$ | 0.01 $_{SubID}$ | | | | 0.01 $_{SubID}$ | | | | 0.01 $_{SubID}$ | | | |
| N | 94 $_{SubID}$ | | | | 94 $_{SubID}$ | | | | 94 $_{SubID}$ | | | |
| Observations | 11551 | | | | 11551 | | | | 11551 | | | |
| Marginal $R^2$ / Conditional $R^2$ | 0.001 / 0.141 | | | | 0.008 / 0.113 | | | | 0.007 / 0.155 | | | |

* p<0.05 ** p<0.01 *** p<0.001

**Interval Related ROI Activity by Mean Motivation Rating**

| | **Rostral dACC** | | | | **Dorsolateral PFC** | | | | **Inferior Frontal Gyrus** | | | |
|---|---|---|---|---|---|---|---|---|---|---|---|---|
| *Predictors* | *Estimates* | *CI* | *Statistic* | *p* | *Estimates* | *CI* | *Statistic* | *p* | *Estimates* | *CI* | *Statistic* | *p* |
| (Intercept) | -0.14 *** | -0.16 – -0.12 | -14.82 | **<0.001** | -0.04 *** | -0.06 – -0.03 | -4.88 | **<0.001** | -0.16 *** | -0.18 – -0.14 | -19.71 | **<0.001** |
| Mean Motivation | -0.00 | -0.01 – 0.01 | -0.33 | 0.739 | -0.00 | -0.01 – 0.00 | -0.61 | 0.543 | -0.00 | -0.01 – 0.01 | -0.33 | 0.745 |
| Interval Length | 0.00 | -0.00 – 0.01 | 1.81 | 0.070 | 0.00 | -0.00 – 0.01 | 1.31 | 0.191 | 0.00 * | 0.00 – 0.01 | 2.23 | **0.026** |
| Interval Session Number | -0.01 ** | -0.01 – -0.00 | -3.05 | **0.002** | -0.02 *** | -0.03 – -0.02 | -10.68 | **<0.001** | -0.01 *** | -0.02 – -0.01 | -6.27 | **<0.001** |
| Mean Motivation:Reward | 0.00 | -0.00 – 0.01 | 1.21 | 0.226 | 0.00 | -0.00 – 0.01 | 1.14 | 0.253 | 0.00 | -0.00 – 0.01 | 0.41 | 0.680 |
| Mean Motivation:Penalty | 0.00 | -0.00 – 0.01 | 1.48 | 0.139 | -0.00 | -0.01 – 0.00 | -0.55 | 0.583 | 0.01 * | 0.00 – 0.01 | 2.46 | **0.014** |
| **Random Effects** | | | | | | | | | | | | |
| $\sigma^2$ | 0.05 | | | | 0.05 | | | | 0.05 | | | |
| $\tau_{00}$ | 0.01 $_{SubID}$ | | | | 0.01 $_{SubID}$ | | | | 0.01 $_{SubID}$ | | | |
| N | 94 $_{SubID}$ | | | | 94 $_{SubID}$ | | | | 94 $_{SubID}$ | | | |
| Observations | 11551 | | | | 11551 | | | | 11551 | | | |
| Marginal $R^2$ / Conditional $R^2$ | 0.001 / 0.130 | | | | 0.009 / 0.130 | | | | 0.004 / 0.111 | | | |

* p<0.05 ** p<0.01 *** p<0.001

**Table S37:** *Difficulty Ratings Predict Neural Activity During Cue Phase*. Linear mixed effects models with ROI activity during the cue phase predicted by self-report ratings of difficulty and included experimental conditions such as interval length and interval session number to control for fatigue effects across the session.

**Cue Related ROI Activity by Difficulty Rating**

| | **Ventral Striatum** | | | | **Anterior Insula** | | | | **Caudal dACC** | | | |
|---|---|---|---|---|---|---|---|---|---|---|---|---|
| *Predictors* | *Estimates* | *CI* | *Statistic* | *p* | *Estimates* | *CI* | *Statistic* | *p* | *Estimates* | *CI* | *Statistic* | *p* |
| (Intercept) | 0.06 * | 0.01 – 0.12 | 2.38 | **0.017** | -0.14 *** | -0.19 – -0.08 | -4.95 | **<0.001** | 0.07 * | 0.02 – 0.12 | 2.51 | **0.012** |
| Difficult | 0.06 *** | 0.03 – 0.08 | 3.99 | **<0.001** | 0.04 ** | 0.01 – 0.07 | 2.68 | **0.007** | 0.03 * | 0.00 – 0.06 | 2.09 | **0.037** |
| Interval Length | -0.00 | -0.02 – 0.02 | -0.06 | 0.952 | 0.01 | -0.01 – 0.03 | 0.68 | 0.499 | 0.02 | -0.01 – 0.04 | 1.49 | 0.137 |
| Interval Session Number | 0.01 | -0.01 – 0.03 | 1.19 | 0.233 | -0.04 *** | -0.07 – -0.02 | -3.70 | **<0.001** | -0.00 | -0.02 – 0.02 | -0.18 | 0.856 |
| Difficult:Reward | 0.01 | -0.01 – 0.03 | 0.82 | 0.413 | 0.01 | -0.01 – 0.04 | 1.05 | 0.294 | 0.01 | -0.01 – 0.04 | 1.00 | 0.315 |
| Difficult:Penalty | -0.02 | -0.04 – 0.00 | -1.64 | 0.100 | 0.00 | -0.02 – 0.03 | 0.38 | 0.703 | 0.00 | -0.02 – 0.03 | 0.26 | 0.793 |
| **Random Effects** | | | | | | | | | | | | |
| $\sigma^2$ | 1.31 | | | | 1.55 | | | | 1.46 | | | |
| $\tau_{00}$ | $0.06_{\text{SubID}}$ | | | | $0.06_{\text{SubID}}$ | | | | $0.06_{\text{SubID}}$ | | | |
| N | $94_{\text{SubID}}$ | | | | $94_{\text{SubID}}$ | | | | $94_{\text{SubID}}$ | | | |
| Observations | 11551 | | | | 11551 | | | | 11551 | | | |
| Marginal $R^2$ / Conditional $R^2$ | 0.003 / 0.043 | | | | 0.002 / 0.039 | | | | 0.001 / 0.039 | | | |

* p<0.05 ** p<0.01 *** p<0.001

**Cue Related ROI Activity by Difficulty Rating**

| | **Rostral dACC** | | | | **Dorsolateral PFC** | | | | **Inferior Frontal Gyrus** | | | |
|---|---|---|---|---|---|---|---|---|---|---|---|---|
| *Predictors* | *Estimates* | *CI* | *Statistic* | *p* | *Estimates* | *CI* | *Statistic* | *p* | *Estimates* | *CI* | *Statistic* | *p* |
| (Intercept) | -0.07 * | -0.12 – -0.02 | -2.56 | **0.010** | -0.02 | -0.08 – 0.04 | -0.69 | 0.493 | -0.11 *** | -0.16 – -0.06 | -4.02 | **<0.001** |
| Difficult | 0.04 * | 0.01 – 0.06 | 2.38 | **0.018** | 0.04 ** | 0.01 – 0.07 | 2.72 | **0.007** | 0.02 | -0.00 – 0.05 | 1.64 | 0.102 |
| Interval Length | 0.01 | -0.01 – 0.03 | 0.86 | 0.392 | 0.01 | -0.01 – 0.04 | 1.17 | 0.242 | 0.01 | -0.01 – 0.03 | 0.68 | 0.498 |
| Interval Session Number | -0.06 *** | -0.08 – -0.03 | -4.78 | **<0.001** | -0.04 *** | -0.06 – -0.02 | -3.55 | **<0.001** | -0.05 *** | -0.07 – -0.02 | -4.11 | **<0.001** |
| Difficult:Reward | 0.01 | -0.02 – 0.03 | 0.71 | 0.480 | 0.01 | -0.02 – 0.03 | 0.71 | 0.479 | 0.00 | -0.02 – 0.03 | 0.22 | 0.829 |
| Difficult:Penalty | 0.00 | -0.02 – 0.03 | 0.26 | 0.797 | 0.00 | -0.02 – 0.03 | 0.11 | 0.916 | -0.00 | -0.03 – 0.02 | -0.03 | 0.976 |
| **Random Effects** | | | | | | | | | | | | |
| $\sigma^2$ | 1.55 | | | | 1.58 | | | | 1.55 | | | |
| $\tau_{00}$ | $0.05_{\text{SubID}}$ | | | | $0.08_{\text{SubID}}$ | | | | $0.05_{\text{SubID}}$ | | | |
| N | $94_{\text{SubID}}$ | | | | $94_{\text{SubID}}$ | | | | $94_{\text{SubID}}$ | | | |
| Observations | 11551 | | | | 11551 | | | | 11551 | | | |
| Marginal $R^2$ / Conditional $R^2$ | 0.003 / 0.037 | | | | 0.002 / 0.048 | | | | 0.002 / 0.035 | | | |

* p<0.05 ** p<0.01 *** p<0.001

**Table S38:** *Difficulty Ratings Predict Neural Activity During Interval Phase*. Linear mixed effects models with ROI activity during the interval phase predicted by self-report ratings of difficulty and included experimental conditions such as interval length and interval session number to control for fatigue effects across the session.

**Interval Related ROI Activity by Difficulty Rating**

| | **Ventral Striatum** | | | | **Anterior Insula** | | | | **Caudal dACC** | | | |
|---|---|---|---|---|---|---|---|---|---|---|---|---|
| *Predictors* | *Estimates* | *CI* | *Statistic* | *p* | *Estimates* | *CI* | *Statistic* | *p* | *Estimates* | *CI* | *Statistic* | *p* |
| (Intercept) | 0.19 *** | 0.17 – 0.20 | 21.48 | **<0.001** | -0.12 *** | -0.14 – -0.11 | -15.00 | **<0.001** | 0.24 *** | 0.22 – 0.25 | 25.54 | **<0.001** |
| Difficult | 0.01 * | 0.00 – 0.01 | 2.19 | **0.028** | 0.01 *** | 0.00 – 0.02 | 3.47 | **0.001** | 0.01 * | 0.00 – 0.01 | 2.15 | **0.032** |
| Interval Length | -0.00 | -0.00 – 0.00 | -0.31 | 0.757 | 0.00 | -0.00 – 0.00 | 0.06 | 0.955 | -0.01 ** | -0.01 – -0.00 | -3.17 | **0.002** |
| Interval Session Number | 0.00 * | 0.00 – 0.01 | 2.54 | **0.011** | -0.02 *** | -0.03 – -0.02 | -10.20 | **<0.001** | -0.02 *** | -0.02 – -0.01 | -8.79 | **<0.001** |
| Difficult:Reward | 0.00 | -0.00 – 0.01 | 0.80 | 0.426 | 0.00 | -0.00 – 0.01 | 0.74 | 0.458 | 0.00 | -0.00 – 0.01 | 1.24 | 0.217 |
| Difficult:Penalty | -0.00 | -0.01 – 0.00 | -1.16 | 0.248 | -0.00 | -0.01 – 0.00 | -1.43 | 0.154 | -0.00 | -0.01 – 0.00 | -1.53 | 0.125 |
| **Random Effects** | | | | | | | | | | | | |
| $\sigma^2$ | 0.04 | | | | 0.05 | | | | 0.04 | | | |
| $\tau_{00}$ | 0.01 $_{SubID}$ | | | | 0.01 $_{SubID}$ | | | | 0.01 $_{SubID}$ | | | |
| N | 94 $_{SubID}$ | | | | 94 $_{SubID}$ | | | | 94 $_{SubID}$ | | | |
| Observations | 11551 | | | | 11551 | | | | 11551 | | | |
| Marginal $R^2$ / Conditional $R^2$ | 0.001 / 0.141 | | | | 0.010 / 0.116 | | | | 0.007 / 0.155 | | | |

** p<0.05 ** p<0.01 *** p<0.001*

**Interval Related ROI Activity by Difficulty Rating**

| | **Rostral dACC** | | | | **Dorsolateral PFC** | | | | **Inferior Frontal Gyrus** | | | |
|---|---|---|---|---|---|---|---|---|---|---|---|---|
| *Predictors* | *Estimates* | *CI* | *Statistic* | *p* | *Estimates* | *CI* | *Statistic* | *p* | *Estimates* | *CI* | *Statistic* | *p* |
| (Intercept) | -0.14 *** | -0.15 – -0.12 | -14.62 | **<0.001** | -0.04 *** | -0.06 – -0.02 | -4.75 | **<0.001** | -0.16 *** | -0.17 – -0.14 | -19.42 | **<0.001** |
| Difficult | 0.00 | -0.00 – 0.01 | 1.55 | 0.121 | 0.01 ** | 0.00 – 0.01 | 2.99 | **0.003** | 0.01 ** | 0.00 – 0.01 | 2.64 | **0.008** |
| Interval Length | 0.00 | -0.00 – 0.01 | 1.41 | 0.159 | 0.00 | -0.00 – 0.01 | 0.67 | 0.501 | 0.00 | -0.00 – 0.01 | 1.80 | 0.071 |
| Interval Session Number | -0.01 ** | -0.01 – -0.00 | -3.02 | **0.003** | -0.02 *** | -0.03 – -0.02 | -10.65 | **<0.001** | -0.01 *** | -0.02 – -0.01 | -6.24 | **<0.001** |
| Difficult:Reward | 0.00 | -0.00 – 0.01 | 0.38 | 0.707 | 0.00 | -0.00 – 0.01 | 0.84 | 0.403 | -0.00 | -0.00 – 0.00 | -0.09 | 0.928 |
| Difficult:Penalty | -0.00 | -0.00 – 0.00 | -0.08 | 0.935 | -0.00 | -0.01 – 0.00 | -1.44 | 0.151 | -0.00 | -0.01 – 0.00 | -0.49 | 0.624 |
| **Random Effects** | | | | | | | | | | | | |
| $\sigma^2$ | 0.05 | | | | 0.05 | | | | 0.05 | | | |
| $\tau_{00}$ | 0.01 $_{SubID}$ | | | | 0.01 $_{SubID}$ | | | | 0.01 $_{SubID}$ | | | |
| N | 94 $_{SubID}$ | | | | 94 $_{SubID}$ | | | | 94 $_{SubID}$ | | | |
| Observations | 11551 | | | | 11551 | | | | 11551 | | | |
| Marginal $R^2$ / Conditional $R^2$ | 0.001 / 0.131 | | | | 0.010 / 0.131 | | | | 0.004 / 0.113 | | | |

** p<0.05 ** p<0.01 *** p<0.001*

**Table S39:** *Arousal Ratings Predict Neural Activity During Cue Phase*. Linear mixed effects models with ROI activity during the cue phase predicted by self-report ratings of arousal, including regressors for experimental conditions such as interval length and interval session number to control for fatigue effects across the session.

**Cue Related ROI Activity by Arousal Ratings**

| | Ventral Striatum | | | | Anterior Insula | | | | Caudal dACC | | | |
|---|---|---|---|---|---|---|---|---|---|---|---|---|
| *Predictors* | *Estimates* | *CI* | *Statistic* | *p* | *Estimates* | *CI* | *Statistic* | *p* | *Estimates* | *CI* | *Statistic* | *p* |
| (Intercept) | 0.06 * | 0.01 – 0.11 | 2.25 | **0.025** | -0.14 *** | -0.19 – -0.08 | -4.84 | **<0.001** | 0.07 * | 0.02 – 0.12 | 2.54 | **0.011** |
| Arousal | 0.04 ** | 0.02 – 0.07 | 3.15 | **0.002** | 0.01 | -0.02 – 0.04 | 0.75 | 0.450 | 0.03 * | 0.01 – 0.06 | 2.52 | **0.012** |
| Interval Length | -0.00 | -0.02 – 0.02 | -0.03 | 0.973 | 0.01 | -0.01 – 0.03 | 0.78 | 0.433 | 0.02 | -0.01 – 0.04 | 1.44 | 0.150 |
| Interval Session Number | 0.01 | -0.01 – 0.03 | 1.18 | 0.239 | -0.04 *** | -0.07 – -0.02 | -3.74 | **<0.001** | -0.00 | -0.02 – 0.02 | -0.20 | 0.841 |
| Arousal:Reward | -0.01 | -0.04 – 0.01 | -1.07 | 0.285 | -0.01 | -0.04 – 0.02 | -0.69 | 0.488 | -0.00 | -0.03 – 0.02 | -0.13 | 0.899 |
| Arousal:Penalty | 0.00 | -0.02 – 0.03 | 0.29 | 0.773 | 0.00 | -0.03 – 0.03 | 0.02 | 0.985 | -0.00 | -0.03 – 0.02 | -0.21 | 0.830 |
| **Random Effects** | | | | | | | | | | | | |
| $\sigma^2$ | 1.31 | | | | 1.56 | | | | 1.46 | | | |
| $\tau_{00}$ | 0.05 $_{SubID}$ | | | | 0.06 $_{SubID}$ | | | | 0.06 $_{SubID}$ | | | |
| N | 94 $_{SubID}$ | | | | 94 $_{SubID}$ | | | | 94 $_{SubID}$ | | | |
| Observations | 11551 | | | | 11551 | | | | 11551 | | | |
| Marginal $R^2$ / Conditional $R^2$ | 0.001 / 0.041 | | | | 0.001 / 0.038 | | | | 0.001 / 0.039 | | | |

* p<0.05 ** p<0.01 *** p<0.001

**Cue Related ROI Activity by Arousal Ratings**

| | Rostral dACC | | | | Dorsolateral PFC | | | | Inferior Frontal Gyrus | | | |
|---|---|---|---|---|---|---|---|---|---|---|---|---|
| *Predictors* | *Estimates* | *CI* | *Statistic* | *p* | *Estimates* | *CI* | *Statistic* | *p* | *Estimates* | *CI* | *Statistic* | *p* |
| (Intercept) | -0.07 * | -0.12 – -0.01 | -2.48 | **0.013** | -0.02 | -0.08 – 0.04 | -0.72 | 0.470 | -0.10 *** | -0.16 – -0.05 | -3.88 | **<0.001** |
| Arousal | 0.00 | -0.02 – 0.03 | 0.23 | 0.814 | 0.03 * | 0.00 – 0.06 | 2.21 | **0.027** | 0.01 | -0.02 – 0.04 | 0.68 | 0.498 |
| Interval Length | 0.01 | -0.01 – 0.03 | 0.95 | 0.341 | 0.01 | -0.01 – 0.04 | 1.13 | 0.259 | 0.01 | -0.02 – 0.03 | 0.61 | 0.540 |
| Interval Session Number | -0.06 *** | -0.08 – -0.03 | -4.80 | **<0.001** | -0.04 *** | -0.06 – -0.02 | -3.57 | **<0.001** | -0.05 *** | -0.07 – -0.03 | -4.13 | **<0.001** |
| Arousal:Reward | -0.01 | -0.04 – 0.01 | -0.93 | 0.351 | -0.01 | -0.04 – 0.02 | -0.86 | 0.389 | -0.02 | -0.05 – 0.01 | -1.37 | 0.170 |
| Arousal:Penalty | 0.01 | -0.02 – 0.03 | 0.51 | 0.608 | 0.02 | -0.01 – 0.04 | 1.19 | 0.233 | 0.00 | -0.02 – 0.03 | 0.24 | 0.811 |
| **Random Effects** | | | | | | | | | | | | |
| $\sigma^2$ | 1.55 | | | | 1.58 | | | | 1.55 | | | |
| $\tau_{00}$ | 0.05 $_{SubID}$ | | | | 0.08 $_{SubID}$ | | | | 0.05 $_{SubID}$ | | | |
| N | 94 $_{SubID}$ | | | | 94 $_{SubID}$ | | | | 94 $_{SubID}$ | | | |
| Observations | 11551 | | | | 11551 | | | | 11551 | | | |
| Marginal $R^2$ / Conditional $R^2$ | 0.002 / 0.035 | | | | 0.002 / 0.047 | | | | 0.002 / 0.035 | | | |

* p<0.05 ** p<0.01 *** p<0.001

**Table S40:** *Arousal Ratings Predict Neural Activity During Interval Phase*. Linear mixed effects models with ROI activity during the interval phase predicted by self-report ratings of arousal, including regressors for experimental conditions such as interval length and interval session number to control for fatigue effects across the session.

**Interval Related ROI Activity by Arousal Ratings**

| | Ventral Striatum | | | | Anterior Insula | | | | Caudal dACC | | | |
|---|---|---|---|---|---|---|---|---|---|---|---|---|
| *Predictors* | *Estimates* | *CI* | *Statistic* | *p* | *Estimates* | *CI* | *Statistic* | *p* | *Estimates* | *CI* | *Statistic* | *p* |
| (Intercept) | 0.19 *** | 0.17 – 0.20 | 21.27 | **<0.001** | -0.13 *** | -0.14 – -0.11 | -15.11 | **<0.001** | 0.23 *** | 0.22 – 0.25 | 25.35 | **<0.001** |
| Arousal | 0.01 ** | 0.00 – 0.01 | 3.06 | **0.002** | 0.01 * | 0.00 – 0.01 | 2.49 | **0.013** | 0.01 ** | 0.00 – 0.01 | 2.69 | **0.007** |
| Interval Length | -0.00 | -0.00 – 0.00 | -0.17 | 0.865 | 0.00 | -0.00 – 0.00 | 0.06 | 0.950 | -0.01 ** | -0.01 – -0.00 | -2.94 | **0.003** |
| Interval Session Number | 0.00 * | 0.00 – 0.01 | 2.54 | **0.011** | -0.02 *** | -0.03 – -0.02 | -10.21 | **<0.001** | -0.02 *** | -0.02 – -0.01 | -8.79 | **<0.001** |
| Arousal:Reward | 0.00 | -0.00 – 0.01 | 1.29 | 0.199 | -0.00 | -0.01 – 0.00 | -1.22 | 0.221 | 0.00 | -0.00 – 0.01 | 1.64 | 0.101 |
| Arousal:Penalty | 0.00 | -0.00 – 0.01 | 0.57 | 0.572 | 0.00 | -0.00 – 0.01 | 0.76 | 0.448 | -0.00 | -0.01 – 0.00 | -1.37 | 0.172 |
| **Random Effects** | | | | | | | | | | | | |
| $\sigma^2$ | 0.04 | | | | 0.05 | | | | 0.04 | | | |
| $\tau_{00}$ | 0.01 $_{SubID}$ | | | | 0.01 $_{SubID}$ | | | | 0.01 $_{SubID}$ | | | |
| N | 94 $_{SubID}$ | | | | 94 $_{SubID}$ | | | | 94 $_{SubID}$ | | | |
| Observations | 11551 | | | | 11551 | | | | 11551 | | | |
| Marginal $R^2$ / Conditional $R^2$ | 0.001 / 0.140 | | | | 0.009 / 0.114 | | | | 0.008 / 0.155 | | | |

* *p<0.05* ** *p<0.01* *** *p<0.001*

**Interval Related ROI Activity by Arousal Ratings**

| | Rostral dACC | | | | Dorsolateral PFC | | | | Inferior Frontal Gyrus | | | |
|---|---|---|---|---|---|---|---|---|---|---|---|---|
| *Predictors* | *Estimates* | *CI* | *Statistic* | *p* | *Estimates* | *CI* | *Statistic* | *p* | *Estimates* | *CI* | *Statistic* | *p* |
| (Intercept) | -0.14 *** | -0.15 – -0.12 | -14.76 | **<0.001** | -0.04 *** | -0.06 – -0.03 | -4.83 | **<0.001** | -0.16 *** | -0.17 – -0.14 | -19.62 | **<0.001** |
| Arousal | 0.00 | -0.00 – 0.01 | 1.77 | 0.077 | 0.01 * | 0.00 – 0.01 | 2.33 | **0.020** | 0.00 | -0.00 – 0.01 | 1.52 | 0.129 |
| Interval Length | 0.00 | -0.00 – 0.01 | 1.35 | 0.177 | 0.00 | -0.00 – 0.01 | 0.72 | 0.473 | 0.00 | -0.00 – 0.01 | 1.78 | 0.074 |
| Interval Session Number | -0.01 ** | -0.01 – -0.00 | -3.02 | **0.003** | -0.02 *** | -0.03 – -0.02 | -10.66 | **<0.001** | -0.01 *** | -0.02 – -0.01 | -6.24 | **<0.001** |
| Arousal:Reward | -0.00 | -0.01 – 0.00 | -0.33 | 0.740 | -0.00 | -0.01 – 0.00 | -0.63 | 0.528 | -0.00 | -0.01 – 0.00 | -1.28 | 0.199 |
| Arousal:Penalty | 0.00 | -0.00 – 0.01 | 1.57 | 0.115 | 0.00 | -0.00 – 0.01 | 0.19 | 0.853 | 0.01 ** | 0.00 – 0.01 | 2.97 | **0.003** |
| **Random Effects** | | | | | | | | | | | | |
| $\sigma^2$ | 0.05 | | | | 0.05 | | | | 0.05 | | | |
| $\tau_{00}$ | 0.01 $_{SubID}$ | | | | 0.01 $_{SubID}$ | | | | 0.01 $_{SubID}$ | | | |
| N | 94 $_{SubID}$ | | | | 94 $_{SubID}$ | | | | 94 $_{SubID}$ | | | |
| Observations | 11551 | | | | 11551 | | | | 11551 | | | |
| Marginal $R^2$ / Conditional $R^2$ | 0.001 / 0.130 | | | | 0.009 / 0.130 | | | | 0.005 / 0.112 | | | |

* *p<0.05* ** *p<0.01* *** *p<0.001*

**Table S41:** *Pleasantness Ratings Predict Neural Activity During Cue Phase*. Linear mixed effects models with ROI activity during the cue phase predicted by self-report ratings of pleasantness, including regressors for experimental conditions such as interval length and interval session number to control for fatigue effects across the session.

**Cue Related ROI Activity by Pleasantness Ratings**

| | Ventral Striatum | | | | Anterior Insula | | | | Caudal dACC | | | |
|---|---|---|---|---|---|---|---|---|---|---|---|---|
| *Predictors* | *Estimates* | *CI* | *Statistic* | *p* | *Estimates* | *CI* | *Statistic* | *p* | *Estimates* | *CI* | *Statistic* | *p* |
| (Intercept) | 0.06 * | 0.00 – 0.11 | 1.97 | **0.049** | -0.15 *** | -0.20 – -0.09 | -4.75 | **<0.001** | 0.06 * | 0.00 – 0.12 | 2.04 | **0.041** |
| Pleasant | -0.01 | -0.03 – 0.02 | -0.59 | 0.553 | -0.02 | -0.04 – 0.01 | -1.44 | 0.151 | 0.01 | -0.02 – 0.03 | 0.70 | 0.485 |
| Interval Length | 0.00 | -0.02 – 0.02 | 0.34 | 0.735 | 0.01 | -0.01 – 0.03 | 0.71 | 0.479 | 0.02 | -0.00 – 0.04 | 1.78 | 0.075 |
| Interval Session Number | 0.01 | -0.01 – 0.03 | 1.15 | 0.249 | -0.04 *** | -0.07 – -0.02 | -3.75 | **<0.001** | -0.00 | -0.02 – 0.02 | -0.21 | 0.833 |
| Pleasant:Reward | -0.00 | -0.04 – 0.04 | -0.17 | 0.868 | 0.01 | -0.04 – 0.05 | 0.26 | 0.797 | 0.01 | -0.04 – 0.05 | 0.25 | 0.800 |
| Pleasant:Penalty | -0.01 | -0.04 – 0.02 | -0.61 | 0.541 | -0.01 | -0.04 – 0.02 | -0.44 | 0.658 | -0.01 | -0.05 – 0.02 | -0.88 | 0.381 |
| **Random Effects** | | | | | | | | | | | | |
| $\sigma^2$ | 1.31 | | | | 1.56 | | | | 1.46 | | | |
| $\tau_{00}$ | 0.05 $_{SubID}$ | | | | 0.06 $_{SubID}$ | | | | 0.06 $_{SubID}$ | | | |
| N | 94 $_{SubID}$ | | | | 94 $_{SubID}$ | | | | 94 $_{SubID}$ | | | |
| Observations | 11551 | | | | 11551 | | | | 11551 | | | |
| Marginal $R^2$ / Conditional $R^2$ | 0.000 / 0.040 | | | | 0.002 / 0.039 | | | | 0.000 / 0.038 | | | |

* p<0.05 ** p<0.01 *** p<0.001

**Cue Related ROI Activity by Pleasantness Ratings**

| | Rostral dACC | | | | Dorsolateral PFC | | | | Inferior Frontal Gyrus | | | |
|---|---|---|---|---|---|---|---|---|---|---|---|---|
| *Predictors* | *Estimates* | *CI* | *Statistic* | *p* | *Estimates* | *CI* | *Statistic* | *p* | *Estimates* | *CI* | *Statistic* | *p* |
| (Intercept) | -0.07 * | -0.13 – -0.01 | -2.37 | **0.018** | -0.04 | -0.10 – 0.03 | -1.14 | 0.254 | -0.11 *** | -0.17 – -0.06 | -3.89 | **<0.001** |
| Pleasant | -0.02 | -0.05 – 0.00 | -1.58 | 0.114 | -0.01 | -0.04 – 0.01 | -0.89 | 0.374 | -0.02 | -0.04 – 0.01 | -1.29 | 0.199 |
| Interval Length | 0.01 | -0.01 – 0.03 | 0.83 | 0.404 | 0.01 | -0.01 – 0.04 | 1.27 | 0.205 | 0.01 | -0.02 – 0.03 | 0.61 | 0.543 |
| Interval Session Number | -0.06 *** | -0.08 – -0.03 | -4.81 | **<0.001** | -0.04 *** | -0.06 – -0.02 | -3.59 | **<0.001** | -0.05 *** | -0.07 – -0.03 | -4.13 | **<0.001** |
| Pleasant:Reward | 0.01 | -0.04 – 0.05 | 0.29 | 0.770 | 0.02 | -0.02 – 0.06 | 0.86 | 0.390 | 0.01 | -0.03 – 0.06 | 0.69 | 0.492 |
| Pleasant:Penalty | 0.01 | -0.02 – 0.04 | 0.50 | 0.614 | -0.01 | -0.04 – 0.02 | -0.60 | 0.545 | 0.01 | -0.02 – 0.04 | 0.59 | 0.552 |
| **Random Effects** | | | | | | | | | | | | |
| $\sigma^2$ | 1.55 | | | | 1.58 | | | | 1.55 | | | |
| $\tau_{00}$ | 0.05 $_{SubID}$ | | | | 0.08 $_{SubID}$ | | | | 0.05 $_{SubID}$ | | | |
| N | 94 $_{SubID}$ | | | | 94 $_{SubID}$ | | | | 94 $_{SubID}$ | | | |
| Observations | 11551 | | | | 11551 | | | | 11551 | | | |
| Marginal $R^2$ / Conditional $R^2$ | 0.002 / 0.036 | | | | 0.002 / 0.048 | | | | 0.002 / 0.035 | | | |

* p<0.05 ** p<0.01 *** p<0.001

**Table S42:** *Pleasantness Ratings Predict Neural Activity During Interval Phase*. Linear mixed effects models with ROI activity during the interval phase predicted by self-report ratings of pleasantness, including experimental conditions such as interval length and interval session number to control for fatigue effects across the session.

**Interval Related ROI Activity by Pleasantness Ratings**

| | Ventral Striatum | | | | Anterior Insula | | | | Caudal dACC | | | |
|---|---|---|---|---|---|---|---|---|---|---|---|---|
| *Predictors* | *Estimates* | *CI* | *Statistic* | *p* | *Estimates* | *CI* | *Statistic* | *p* | *Estimates* | *CI* | *Statistic* | *p* |
| (Intercept) | 0.18 *** | 0.17 – 0.20 | 20.63 | **<0.001** | -0.13 *** | -0.15 – -0.11 | -15.20 | **<0.001** | 0.23 *** | 0.21 – 0.25 | 24.28 | **<0.001** |
| Pleasant | 0.00 | -0.00 – 0.00 | 0.27 | 0.787 | -0.01 *** | -0.01 – -0.00 | -3.33 | **0.001** | 0.00 | -0.00 – 0.01 | 1.24 | 0.214 |
| Interval Length | -0.00 | -0.00 – 0.00 | -0.26 | 0.793 | -0.00 | -0.00 – 0.00 | -0.28 | 0.780 | -0.01 ** | -0.01 – -0.00 | -2.98 | **0.003** |
| Interval Session Number | 0.00 ** | 0.00 – 0.01 | 2.58 | **0.010** | -0.02 *** | -0.03 – -0.02 | -10.21 | **<0.001** | -0.02 *** | -0.02 – -0.01 | -8.75 | **<0.001** |
| Pleasant:Reward | 0.01 ** | 0.00 – 0.02 | 2.79 | **0.005** | 0.01 ** | 0.00 – 0.02 | 2.77 | **0.006** | 0.01 ** | 0.00 – 0.02 | 2.88 | **0.004** |
| Pleasant:Penalty | 0.01 *** | 0.00 – 0.02 | 3.63 | **<0.001** | 0.01 * | 0.00 – 0.01 | 2.14 | **0.032** | 0.01 * | 0.00 – 0.01 | 2.34 | **0.019** |
| **Random Effects** | | | | | | | | | | | | |
| $\sigma^2$ | 0.04 | | | | 0.05 | | | | 0.04 | | | |
| $\tau_{00}$ | 0.01 $_{SubID}$ | | | | 0.01 $_{SubID}$ | | | | 0.01 $_{SubID}$ | | | |
| N | 94 $_{SubID}$ | | | | 94 $_{SubID}$ | | | | 94 $_{SubID}$ | | | |
| Observations | 11551 | | | | 11551 | | | | 11551 | | | |
| Marginal $R^2$ / Conditional $R^2$ | 0.002 / 0.140 | | | | 0.010 / 0.116 | | | | 0.007 / 0.155 | | | |

* p<0.05 ** p<0.01 *** p<0.001

**Interval Related ROI Activity by Pleasantness Ratings**

| | Rostral dACC | | | | Dorsolateral PFC | | | | Inferior Frontal Gyrus | | | |
|---|---|---|---|---|---|---|---|---|---|---|---|---|
| *Predictors* | *Estimates* | *CI* | *Statistic* | *p* | *Estimates* | *CI* | *Statistic* | *p* | *Estimates* | *CI* | *Statistic* | *p* |
| (Intercept) | -0.14 *** | -0.16 – -0.12 | -14.80 | **<0.001** | -0.05 *** | -0.07 – -0.03 | -5.31 | **<0.001** | -0.16 *** | -0.18 – -0.15 | -19.12 | **<0.001** |
| Pleasant | -0.01 *** | -0.02 – -0.01 | -5.02 | **<0.001** | -0.01 * | -0.01 – -0.00 | -2.50 | **0.012** | -0.01 *** | -0.02 – -0.01 | -5.98 | **<0.001** |
| Interval Length | 0.00 | -0.00 – 0.01 | 0.53 | 0.595 | 0.00 | -0.00 – 0.00 | 0.41 | 0.683 | 0.00 | -0.00 – 0.01 | 0.95 | 0.341 |
| Interval Session Number | -0.01 ** | -0.01 – -0.00 | -3.03 | **0.002** | -0.02 *** | -0.03 – -0.02 | -10.66 | **<0.001** | -0.01 *** | -0.02 – -0.01 | -6.26 | **<0.001** |
| Pleasant:Reward | 0.01 ** | 0.00 – 0.02 | 3.05 | **0.002** | 0.01 ** | 0.00 – 0.02 | 2.69 | **0.007** | 0.01 ** | 0.00 – 0.02 | 2.94 | **0.003** |
| Pleasant:Penalty | 0.01 ** | 0.00 – 0.01 | 3.02 | **0.003** | 0.00 | -0.00 – 0.01 | 1.40 | 0.161 | 0.01 *** | 0.00 – 0.02 | 3.62 | **<0.001** |
| **Random Effects** | | | | | | | | | | | | |
| $\sigma^2$ | 0.05 | | | | 0.05 | | | | 0.05 | | | |
| $\tau_{00}$ | 0.01 $_{SubID}$ | | | | 0.01 $_{SubID}$ | | | | 0.01 $_{SubID}$ | | | |
| N | 94 $_{SubID}$ | | | | 94 $_{SubID}$ | | | | 94 $_{SubID}$ | | | |
| Observations | 11551 | | | | 11551 | | | | 11551 | | | |
| Marginal $R^2$ / Conditional $R^2$ | 0.004 / 0.133 | | | | 0.010 / 0.133 | | | | 0.008 / 0.118 | | | |

* p<0.05 ** p<0.01 *** p<0.001

**Table S43:** Regression Equations for Hierarchical Bayesian Estimation of drift diffusion model parameters. We estimated drift rate and threshold as a function of reward level, penalty level, and controlled for the interval-level collapsing bound (scaledRunningTime) and congruency. Non-decision time was fitted as a free parameter. The models additionally included intertrial variability in drift and non-decision time, and modeled p_outliers to downweight the influence of extreme RTs.

| **Models** | **Regression Equations** |
|---|---|
| Behavior Only | $a \sim Reward + Penalty + scaledRunningTime$<br>$v \sim Reward + Penalty + Congruency$<br>$z \sim Congruency$<br>$t \sim Intercept$<br>Includes: p_outlier, sv, st |
| VS<br>(Cue, Interval) | $a \sim Reward + Penalty + Rew{:}VS + Pen{:}VS + scaledRunningTime$<br>$v \sim Reward + Penalty + Congruency$<br>$z \sim Congruency$<br>$t \sim Intercept$<br>Includes: p_outlier, sv, st |
| AI<br>(Cue, Interval) | $a \sim Reward + Penalty + Rew{:}VS + Pen{:}VS + scaledRunningTime$<br>$v \sim Reward + Penalty + Congruency$<br>$z \sim Congruency$<br>$t \sim Intercept$<br>Includes: p_outlier, sv, st |
| Caudal dACC<br>(Cue, Interval) | $a \sim Reward + Penalty + Rew{:}VS + Pen{:}VS + scaledRunningTime$<br>$v \sim Reward + Penalty + Congruency$<br>$z \sim Congruency$<br>$t \sim Intercept$<br>Includes: p_outlier, sv, st |
| Rostral dACC<br>(Cue, Interval) | $a \sim Reward + Penalty + Rew{:}VS + Pen{:}VS + scaledRunningTime$<br>$v \sim Reward + Penalty + Congruency$<br>$z \sim Congruency$<br>$t \sim Intercept$<br>Includes: p_outlier, sv, st |
| DLPFC<br>(Cue, Interval) | $a \sim Reward + Penalty + Rew{:}VS + Pen{:}VS + scaledRunningTime$<br>$v \sim Reward + Penalty + Congruency$<br>$z \sim Congruency$<br>$t \sim Intercept$<br>Includes: p_outlier, sv, st |
| IFG<br>(Cue, Interval) | $a \sim Reward + Penalty + Rew{:}VS + Pen{:}VS + scaledRunningTime$<br>$v \sim Reward + Penalty + Congruency$<br>$z \sim Congruency$<br>$t \sim Intercept$<br>Includes: p_outlier, sv, st |

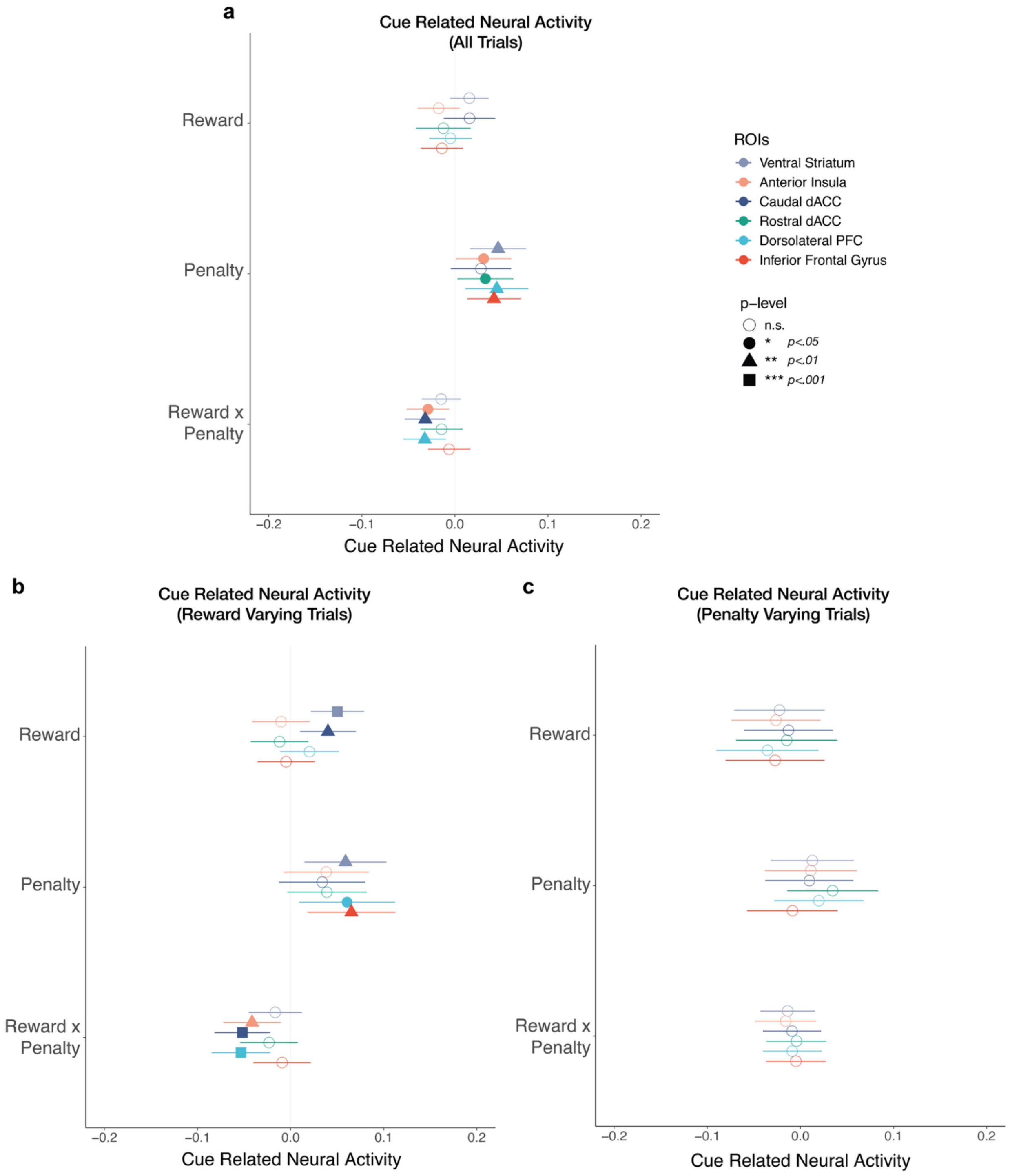


**Figure S1:** *Cue-Related Neural Activity by Reward and Penalty Incentives.* Linear mixed effects models were conducted of cue-related activity predicted by reward, penalty, controlling for nuisance regressors such as interval congruency, interval length, and interval session number. The plots correspond to the models in Tables S1-S6 with a) all intervals combined, b) intervals where reward value was varied and penalty was kept constant, and c) intervals where penalty value was varied and reward was kept constant.

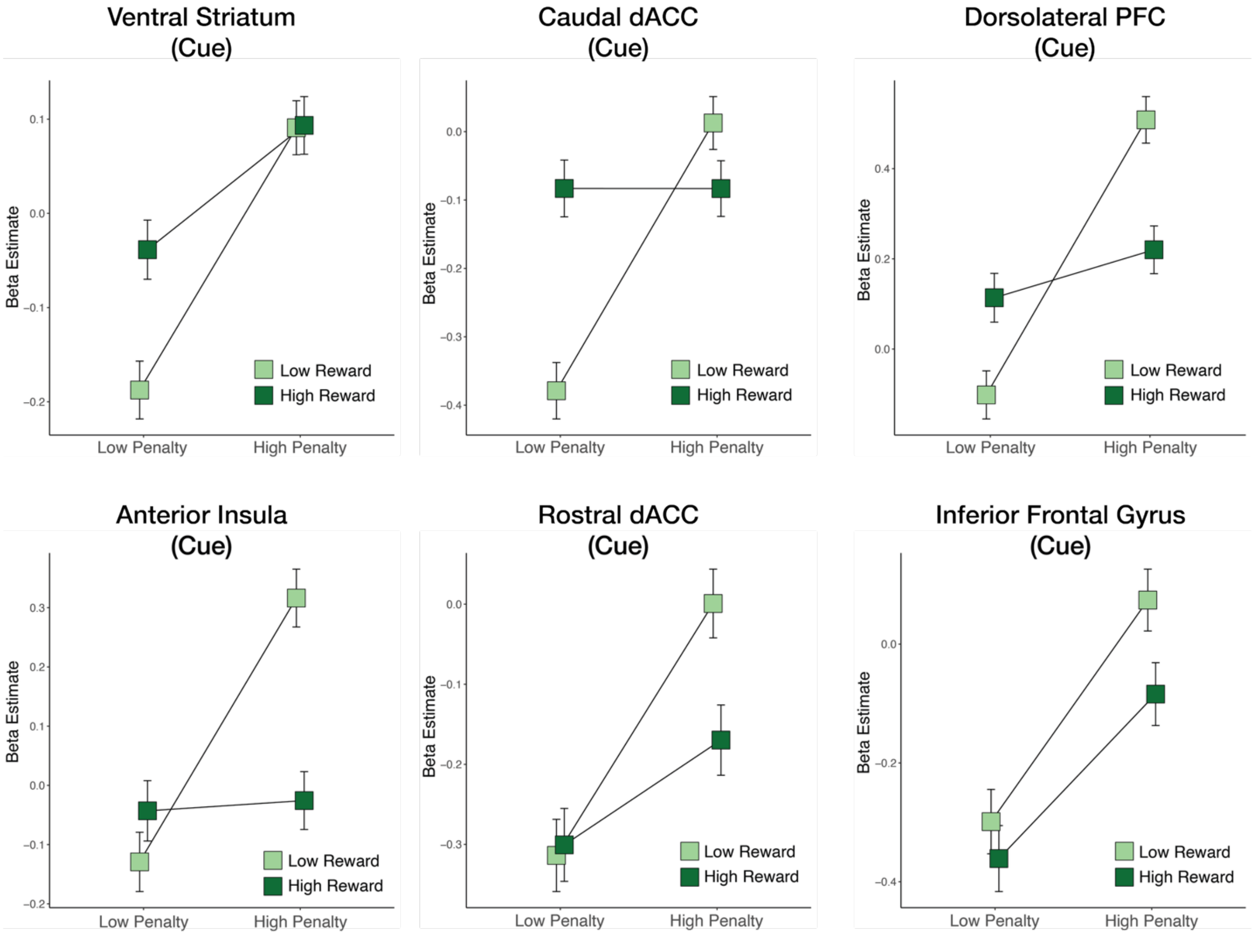


**Figure S2:** *Cue-Related Neural Activity by Reward and Penalty Incentives*. Error bars for the beta estimates represent standard error.

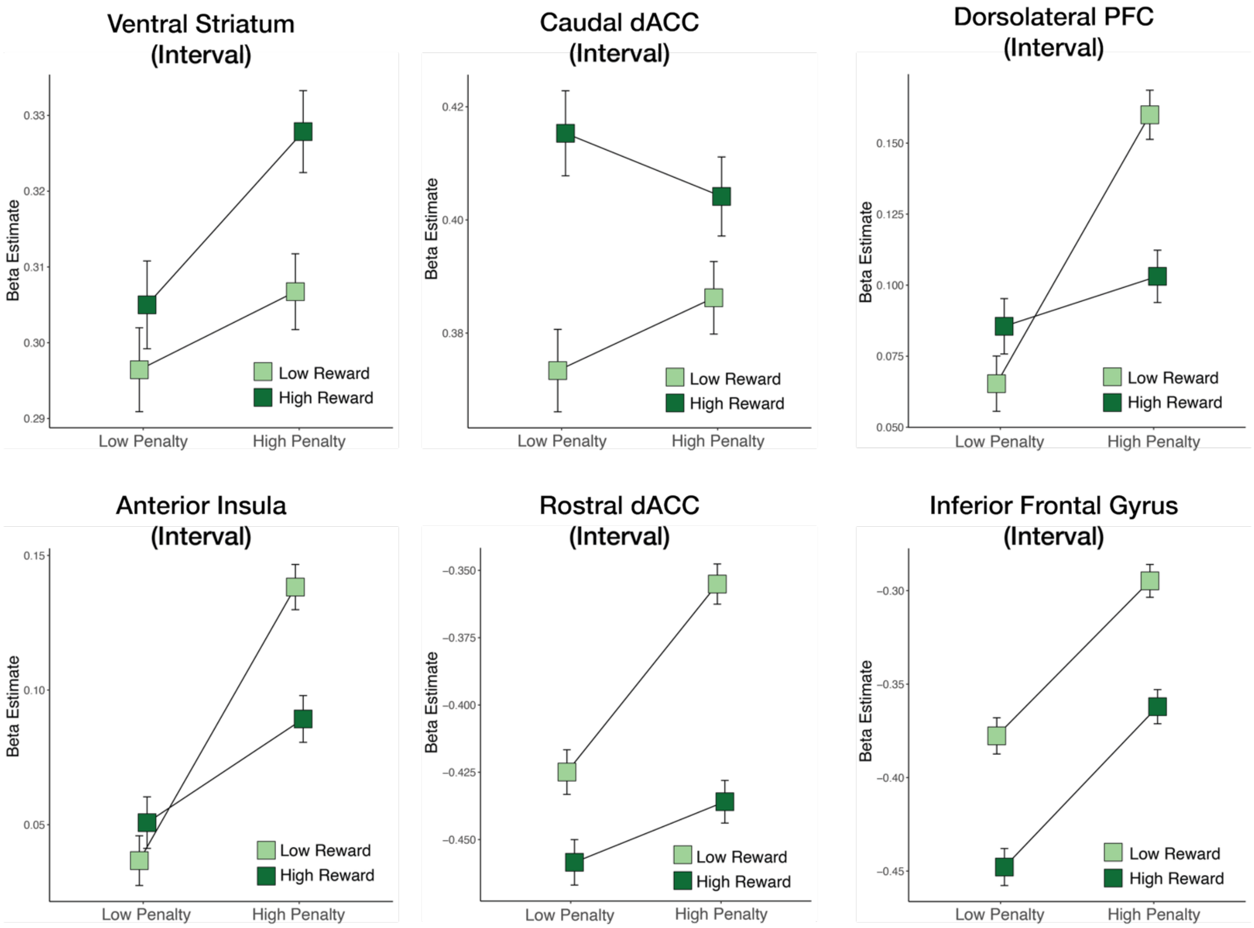


**Figure S3:** *Interval-Related Neural Activity by Reward and Penalty Incentives.* Error bars for the beta estimates represent standard error.

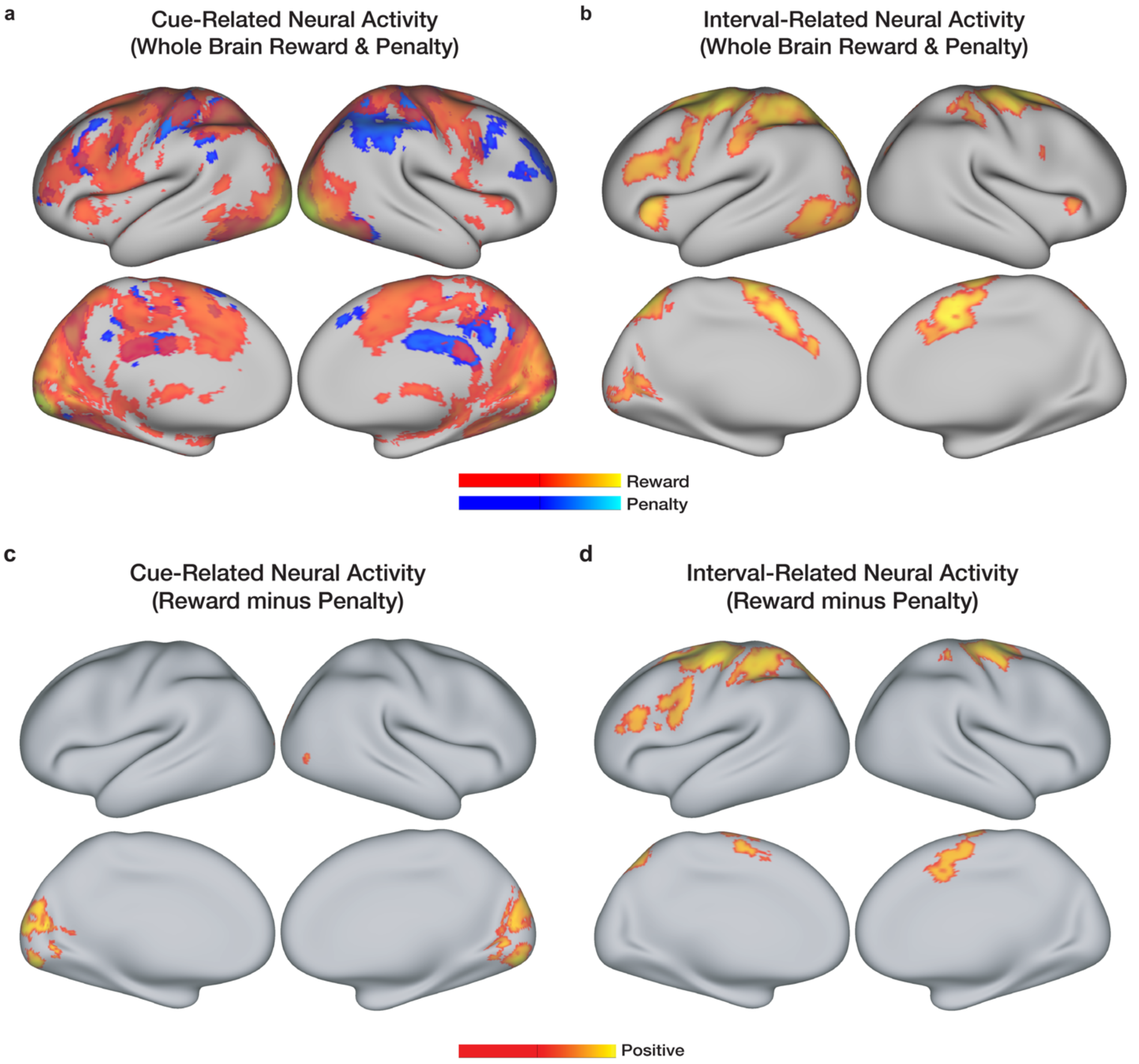


**Figure S4:** *Whole Brain Activation.* a) Whole Brain activity during the cue phase reveal convergence with ROI results, with reward associated with caudal dACC and striatum, whereas penalty was associated with rostral dACC. b) Whole brain activity during the interval phase revealed only reward-related activation in dACC, anterior insula. c) A Reward Minus Penalty contrast during the cue phase revealed greater activation for rewards in visual cortex. d) A Reward Minus Penalty contrast during the interval phase revealed greater activation for rewards in lateral prefrontal cortex and motor cortex. There was no significant activation for Penalty minus Reward. Unless otherwise specified, all reported activations survived whole-brain correction using threshold-free cluster enhancement[23] with familywise error correction at $p<.05$.

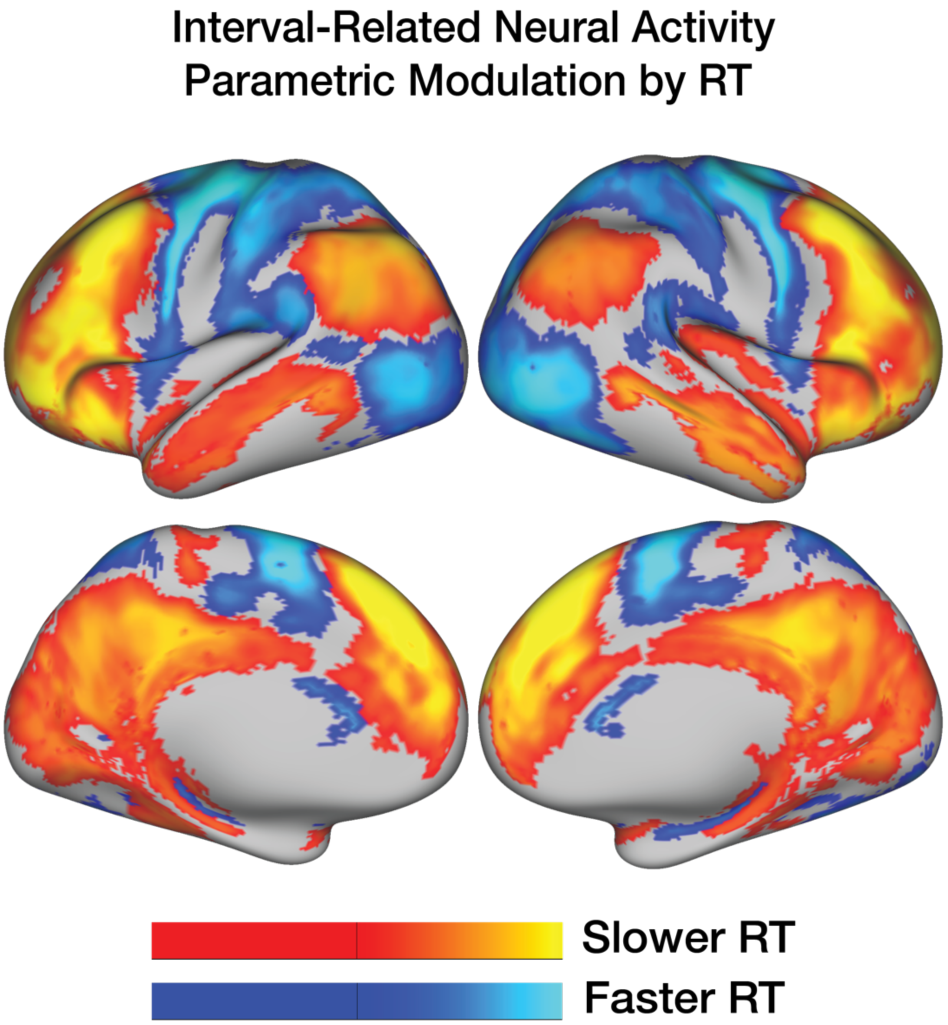


**Figure S5:** *Whole Brain Activation by Parametric Modulation of Response Time.* Parametric modulation of RT on interval-related neural activity revealed that slower RT was associated with greater activation in the medial prefrontal cortex, dorsolateral prefrontal cortex, temporal cortex, and parietal cortex, whereas faster RT was associated with supplementary motor cortex, ventral visual cortex, and dorsal striatum. Unless otherwise specified, all reported activations survived whole-brain correction using threshold-free cluster enhancement[23] with familywise error correction at $p<.05$.

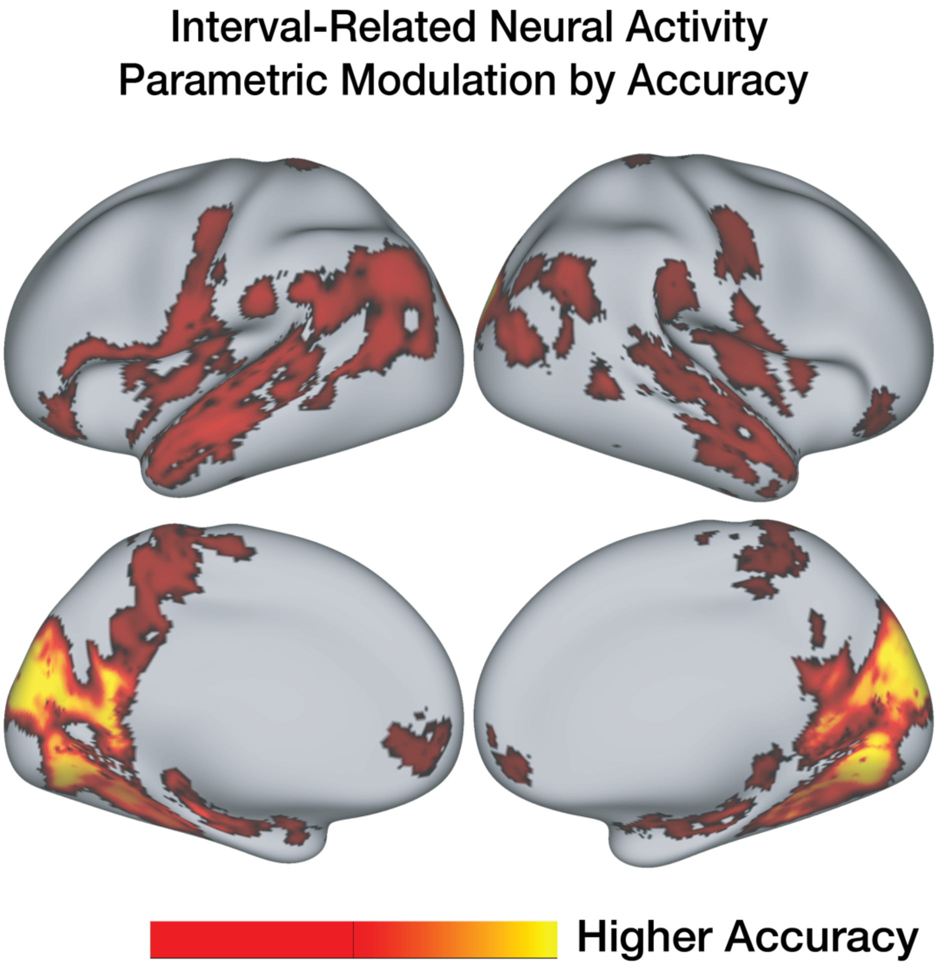


**Figure S6:** *Whole Brain Activation by Parametric Modulation of Accuracy*. Parametric modulation of accuracy on interval-related activity revealed higher accuracy was associated with ventromedial prefrontal cortex, as well as areas in ventral and occipital cortex. There was no significant activation for low accuracy. Unless otherwise specified, all reported activations survived whole-brain correction using threshold-free cluster enhancement[23] with familywise error correction at $p<.05$.

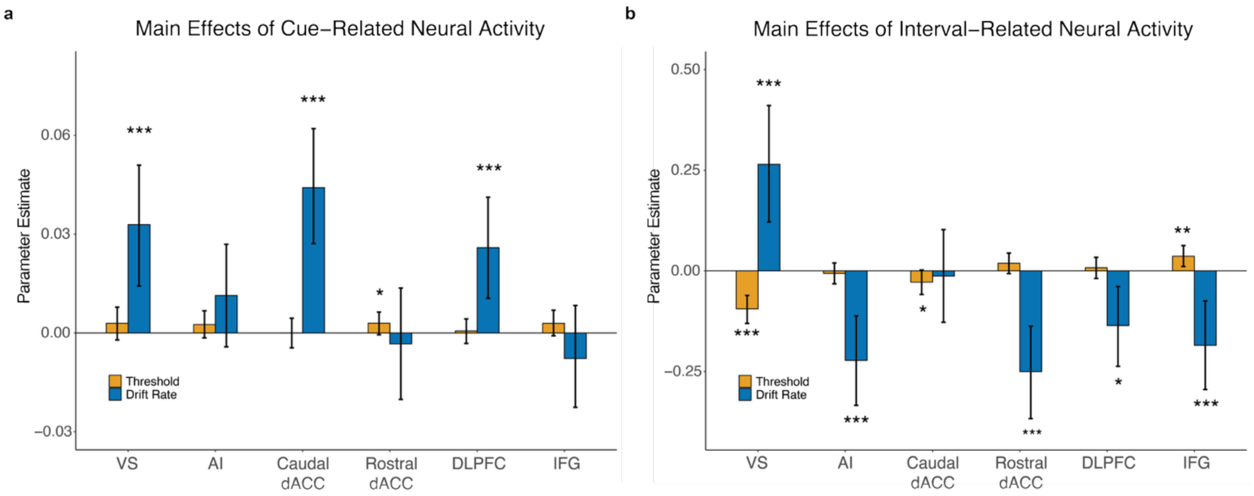


**Figure S7:** *Posterior Estimates of Main Effects of Neural Activity on drift rate and threshold*. Barplot of the posterior estimates of main effects of neural activity from model-based fMRI analyses (corresponding to Figure 3 2D scatterplot). Error bars represent 95% credible intervals.

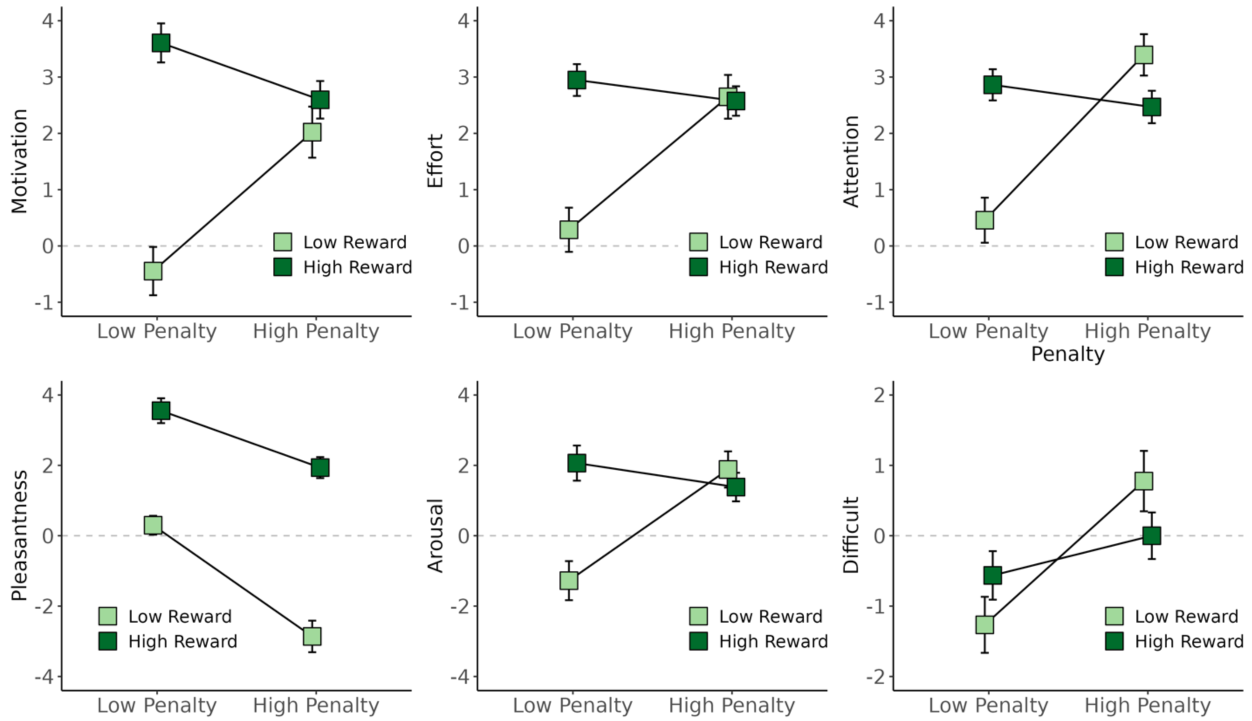


**Figure S8.** *Self-Report Affect and Motivation Ratings*. Individuals rated each of the four incentive cues (e.g., High Reward / High Penalty, Low Reward / Low Penalty, High Reward / Low Penalty, Low Reward / High Penalty). on a scale from -5 to -5 their motivation, effort, attention, pleasantness, arousal, and difficulty. Mean and standard error for all ratings for each incentive cue are shown.

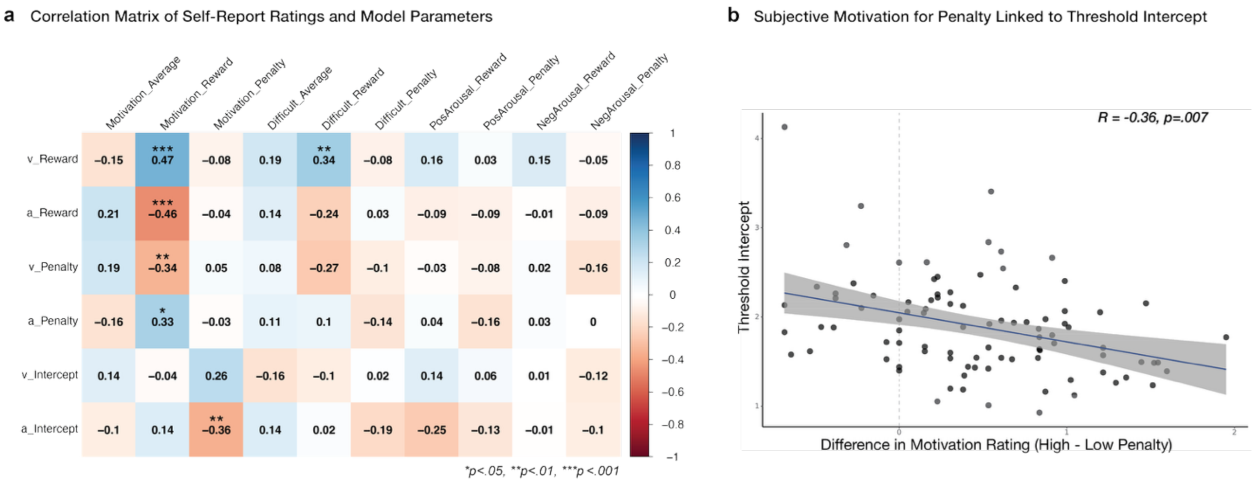


**Figure S9:** *Associations between Self-Report Ratings and DDM Parameters.* b) Correlation matrix of average and difference scores of subjective ratings (motivation, difficulty, positive and negative arousal) and model-based parameter estimates of task performance (drift rate v, threshold a). All p-values are FDR corrected using the Benjamini-Hochberg procedure. c) Individuals who reported greater differentiation in subjective motivation for high vs. low penalty also demonstrated greater degree of modulation of threshold intercept.

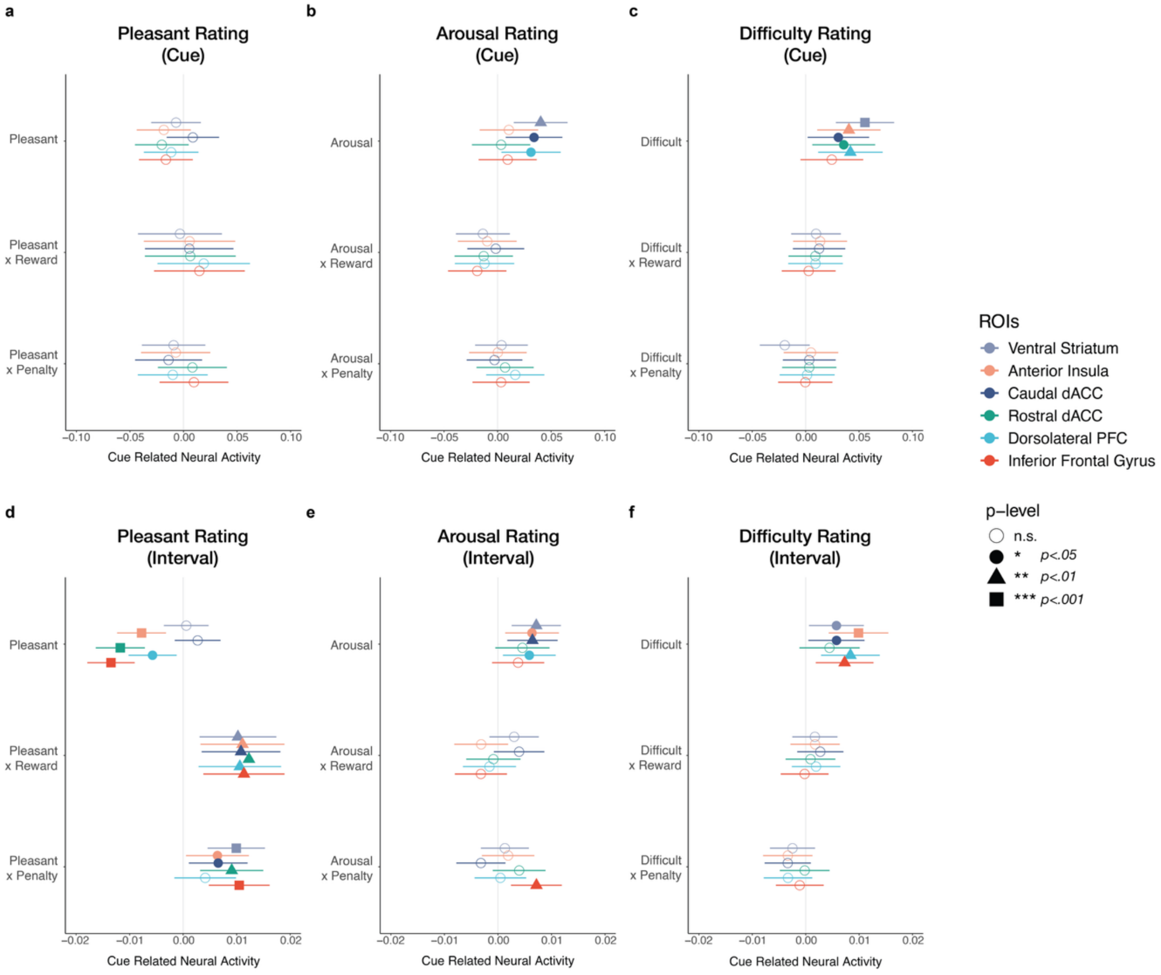


**Figure S10:** *Pleasantness, Arousal, and Difficulty Ratings Predicted by ROIs.* a-c) Cue-Related Neural Activity predicts self-reported ratings of pleasantness, arousal, and difficulty. B) Interval-Related Neural Activity predicts self-reported measures of pleasantness, arousal, and difficulty. We conducted linear mixed models with difficulty ratings predicted by ROI and plot the beta estimate associated with each ROI's fixed effect. Error bars represent 95% confidence intervals.

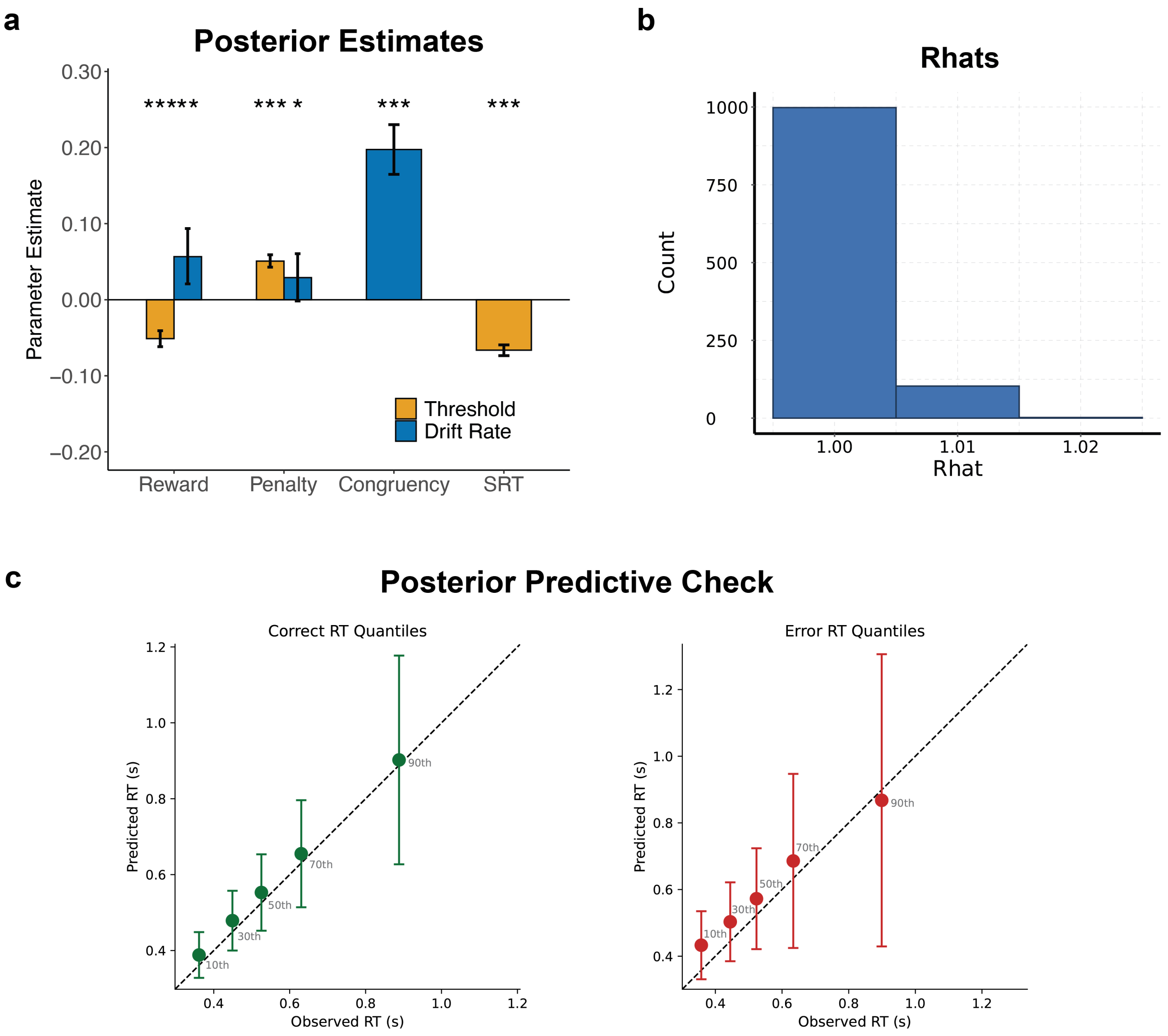


**Figure S11:** *Model Validation and Posterior Predictive Checks of Base Drift Diffusion Model.* A) Posterior estimates of drift diffusion model parameters fitted to choice and RT, as predicted by reward and penalty. The model included regressors for scaled linear running timing which tracked the interval-level (SRT) and Congruency (See Leng et al., 2021 for model optimization and selection). B) Model convergence was tested with the Gelman-Rubin statistic $Rhat$ , which compares between-chain and within-chain variances for model parameters across the five Markov chains. The maximum $Rhat$ across all parameters in was 1.022, indicating that all chains converged successfully. C) Posterior Predictive Checks demonstrate that the selected model reproduces key behavioral patterns from the reward and penalty incentive manipulations.
